# Plant demographic knowledge is biased towards short-term studies of temperate-region herbaceous perennials

**DOI:** 10.1101/2021.04.25.441327

**Authors:** Gesa Römer, Johan P. Dahlgren, Roberto Salguero-Gómez, Iain M. Stott, Owen R. Jones

**Author notes:** Corresponding authors: Gesa Römer; Owen R. Jones.

## Abstract

1. Plant population dynamics research has a long history, and data collection rates have increased through time. The inclusion of this information in databases enables researchers to investigate the drivers of demographic patterns globally and study life history evolution.
2. Studies aiming to generalise demographic patterns rely on data being derived from a representative sample of populations. However, the data are likely to be biased, both in terms of the species and ecoregions investigated and in how the original studies were conducted.
3. Matrix population models (MPMs) are a widely-used tool in plant demography, so an assessment of publications that have used MPMs is a convenient way to assess the distribution of plant demographic knowledge. We assessed bias in this knowledge using data from the COMPADRE Plant Matrix Database, which contains MPMs for almost 800 plant species.
4. We show that tree species and tropical ecoregions are under-represented, while herbaceous perennials and temperate ecoregions are over-represented. In addition, there is a positive association between the number of studies per country and the wealth of the country. Furthermore, we found a strong tendency towards low spatiotemporal replication: More than 50% of the studies were conducted over fewer than 4 years, and only 17% of the studies have replication across >3 sites. This limited spatiotemporal coverage means that the data may not be representative of the environmental conditions experienced by the species.
5. Synthesis: The biases and knowledge gaps we identify are a challenge for the progress of theory and limit the usefulness of current data for determining patterns that would be useful for conservation decisions, such as determining general responses to climate change. We urge researchers to close these knowledge gaps with novel data collection.

## Introduction

Population ecologists aim to understand and predict population dynamics using demographic data that includes the vital rates of survival, reproduction, and development. Their efforts include examining population responses to changes in climate, land use, and management (Silva *et al*., 1991; Buhler *et al*., 1997; Eriksson *et al.,* 2002; Morris *et al*., 2008; Colautti & Barrett, 2013). Demographic data are also crucial for robust population viability analyses of threatened and invasive species (Morris & Doak, 2002; Hansen & Wilson, 2006; Rueda-Cediel *et al*., 2019). Besides single-species studies, researchers have conducted comparative analyses investigating broad demographic and life history patterns among species. These comparative analyses have aided the development of general theories of life history variation, including r-K selection theory (MacArthur & Wilson, 1967; Gunderson, 1980), Grime’s C-S-R triangle (Grime, 1974; Silvertown *et al*. 1992), Stearns’ fast-slow continuum (Stearns, 1992; Franco & Silvertown, 1996; Salguero-Gómez *et al*., 2016) and reproductive strategies continuum (Salguero-Gómez 2017). The empirical exploration of these themes requires large quantities of data from diverse species experiencing a wide range of environmental conditions.

Comparative analyses often rely on the collation of published data to obtain sufficient sample sizes. There are numerous recent examples of this (*e.g.,* Iriondo, 2009; Dalgleish *et al*., 2010; Bullock *et al*., 2012; Burns *et al*., 2013), and large-scale collaborative efforts to collate global demographic and life history and related data are increasingly common (*e.g.,* GBIF: The Global Biodiversity Information Facility; Wright *et al*., 2004; Loh *et al*., 2005; NERC Centre for Population Biology, 2010; Kattge *et al*., 2011; Salguero-Gómez *et al*., 2015). These databases provide a rich resource for workers focussing on life history strategies and demographic performance.

One of the most frequently used tools to describe a species’ demography and life history are matrix population models (MPMs, Crone *et al*., 2011). MPMs depict a population’s life cycle in terms of survival, reproduction, and transitions among discrete life stages (Leslie, 1945; Lefkovitch, 1965; Caswell, 2001). MPMs are particularly useful because they have well-understood mathematical properties, and measures derived from MPMs are comparable across diverse species (Silvertown *et al*., 1993; Caswell, 2001; Salguero-Gómez & Kroon, 2010). The COMPADRE Plant Matrix Database (Salguero-Gómez *et al*., 2015) is the most comprehensive database of plant studies using MPMs and thus reflects our collective knowledge of plant demography. The contents of databases like COMPADRE were not explicitly collected for inclusion in large databases but rather for the disparate purposes of the many original studies. Although these large databases may be an unbiased (or even complete) sampling of the literature, researchers likely focus on species or geographical areas of particular interest. The resulting data collections are likely to be similarly taxonomically, geographically, or methodologically biased.

Bias of this nature has far-reaching consequences for our understanding of plant demography and could limit the usefulness of databases like COMPADRE for comparative analyses. To identify potential biases in plant demographic data and discuss their implications, we used COMPADRE to address the following questions: (1) When have the studies been published? (2) Where is the research done? (3) For which species and populations do we have demographic data? (4) How are the MPMs constructed? More precisely, we tested the following nine hypotheses (H1-H9):

### WHEN HAVE THE STUDIES BEEN PUBLISHED?

We expected (**H1**) to see that the proportion of published plant ecology articles that use MPMs has increased through time, reflecting the growing importance of demographic research within plant ecology.

### WHERE IS THE RESEARCH DONE?

To assess the potential geographic bias of plant demographic studies, we focussed on continental, ecoregion, and country-level biases. We also examined the relationship between the number of studies and the country’s wealth where the study was carried out (as indicated by gross domestic product, GDP). We expected (**H2**) Europe and North America to be over-represented and, likewise, that temperate ecoregions (which characterise these continents) would dominate. Further, we expected (**H3**) wealthier countries would be over-represented compared to their poorer counterparts since they have more funds for research.

### FOR WHICH SPECIES AND POPULATIONS DO WE HAVE DEMOGRAPHIC DATA?

We expected (**H4**) the representation of growth forms would not be proportional to their natural abundance, with herbaceous perennials being over-represented and some growth forms being almost absent. This potential bias is important because, if true, it would limit opportunities to make general inferences on the demography of poorly-represented growth forms. Furthermore, we expected to find (**H5**) a tendency to preferentially study threatened species because they are of particular interest in population ecology (Morris & Doak 2002). We also expected (**H6**) a trend towards choosing flourishing populations (*i.e.*, those with *λ*>1) for data collection, reflecting the researchers’ desire to ensure the long-term viability of their project.

### HOW ARE THE MPMS CONSTRUCTED?

The usefulness of individual demographic studies for comparative analyses is sometimes limited by the methods used to construct the MPMs. We explored this by examining within-study spatio-temporal replication and MPM dimension, which can influence demographic quantities calculated from MPMs, including asymptotic population growth rate (Salguero-Gómez & Plotkin, 2010). We expected (**H7**) low rates of temporal and spatial replication, meaning that the data may not represent adequately the environmental conditions experienced by the population/species. We expected (**H8**) that matrix dimension would vary widely, with a tendency for the MPMs of long-lived species such as trees to have a greater dimension.

Finally, we analysed the prevalence of a widespread simplification approach used in parameterisation: the assumption of transition constancy in two or more consecutive stages. This can occur, for example, when reproduction or survival in consecutive stages (*e.g.*, small, medium, large plants) is assumed to be constant. Although researchers may justify this simplifying assumption based on data limitations, estimates derived from such simplified MPMs may be inaccurate, limiting their usefulness in comparative work. Studies featuring analyses of life expectancy or generation time (Gaillard *et al*., 2005; Staerk *et al*., 2019), ageing trajectories (Baudisch, 2011; Baudisch & Stott, 2019), and transient population dynamics (Stott *et al*., 2011) are all likely to be marred by the widespread use of this assumption. Despite this problem, we expected (**H9**) a large proportion of studies to parameterise matrices using average transition probabilities and fecundity estimates across stages in this way.

We discuss the implications of the biases we identify for several applications. Our results highlight plant demographic knowledge gaps for assessing general patterns, and we encourage researchers to close these taxonomic, biogeographic, and methodological gaps going forward.

## Materials and methods

To quantify potential biases in our knowledge of plant demography, we used the COMPADRE Plant Matrix Database version 5.0.0 (Salguero-Gómez *et al*., 2015). Although COMPADRE also contains data on red and brown algae, and lichens, we restrict our analysis to plants (*i.e.*, land plants and green algae) (Cavalier-Smith, 1981). We analysed our data using R version 4.0.4 (R Core Team, 2021).

Data in COMPADRE are organised by research publication such that particular species can appear multiple times in different articles, and a single publication can include several species. We derived our sample from 641 articles on 746 species. Most articles (547) focussed on single species while 94 focussed on multiple species (2-30 species). In some cases, the archived MPM represents the element-by-element average across several transitions (*e.g.*, the average of 5 years of data). However, COMPADRE also often includes data for the individual transitions (*e.g.*, annual transitions are most commonly used, and COMPADRE thus often includes data on the transition from year 1 to 2, and another for year 2 to 3 and so on). Similarly, articles often include several MPMs representing different experimental treatments and/or different spatial areas for a given year or set of years. Our data set included 925 species-by-article combinations and a total of 9,022 MPMs. In addition to the MPMs, we use COMPADRE metadata on geolocation, ecoregion, growth form, taxonomy, and study timeframe, as well as the MPM projection interval, to examine potential temporal, biogeographic and taxonomic biases. We analyse this data in several ways. For most analyses, we use the entire data set but, due to specific requirements for some analyses, we subset the data for some parts of our study. For full transparency, we include the analysis code as supplementary information.

### WHEN HAVE THE STUDIES BEEN PUBLISHED?

To assess temporal trends in the publication of demography-focussed articles in plant ecology (**H1**), we examined the estimated proportion of articles published in the Journal of Ecology between 1991-2019 that used MPMs. We chose the *Journal of Ecology* as a proxy for the field of plant ecology because it is among the leading and the oldest journals for this field and is thus likely to reflect the temporal development of the discipline. To do this, we downloaded metadata for all of the journal articles for 1991-2019 from the Web of Science. We queried this dataset with the search terms [*projection model* OR *matrix model* OR *MPM*] to identify those that used MPMs. We then compared graphically the estimated percentage of articles that included MPMs.

To gauge the completeness of COMPADRE’s data holdings, we compared COMPADRE’s currently available data (to February 2019) to the COMPADRE team’s curated list of articles targeted for eventual inclusion (data provided by Haydee Hernández-Yañez, pers. comm., 2019).

### WHERE IS THE RESEARCH DONE?

#### Biogeography

We characterised biases in the distribution of studies among continents and ecoregions (**H2**). We first quantified the density of studied species (*i.e.*, number of studied species per unit area (n/km^2^) of each country). We then compared the distribution of species among ecoregions in COMPADRE with the estimated actual species distribution in nature among those ecoregions worldwide using Pearson’s Chi-squared tests (hereafter, *χ^2^*-tests) and *post-hoc* proportion z-tests (using `prop.test` in R). We could do this because COMPADRE assigns each studied population to one or more of Olson’s 14 ecoregions (Olson *et al*., 2001). In some cases, populations were assigned to multiple closely-related ecoregions (*e.g.*, different types of temperate forests). To simplify the analysis, we collapsed Olson’s ecoregions into five broader categories: tropical (Olson’s ecoregions TMB, TDB, TSC and TGV, see Table S1 for explanation), temperate (TBM, TCF, TGS), Mediterranean/desert (MED, DES), tundra/boreal (BOR, TUN, MON), and wetland (MAN, FGS). We extracted the estimated number of species naturally occurring in each region from Kier (Kier *et al*., 2005). Kier *et al*. (2005) did not include bryophytes or algae, so we excluded them from this comparison.

#### Country-specific wealth (GDP)

We used country-level per-capita GDP for 2017 (World Bank, 2018) to examine whether wealthy countries are overrepresented in COMPADRE (**H3**). To do this, we used a Poisson generalised linear model (GLM) (log-link) with log-transformed GDP as the explanatory variable and the number of demographic studies as the response variable. Log-transformation of GDP was necessary to improve the fit of the model. We included only countries with at least one demographic study on plants to avoid a zero-inflated model.

### FOR WHICH SPECIES AND POPULATIONS DO WE HAVE DEMOGRAPHIC DATA?

#### Taxonomy

To characterise potential biases in taxonomy and growth form (**H4**), we first analysed the distribution of taxa in COMPADRE among the taxonomic categories of angiosperm *vs.* gymnosperm, monocot *vs.* eudicot (for the 849 angiosperms only), and Family. We then used the COMPADRE database metadata variable `OrganismType` (hereafter, growth form), which includes a range of paraphyletic growth form categories such as “tree”, “herbaceous perennial”, and “shrub”. We compared the distribution of angiosperms *vs.* gymnosperms in COMPADRE with estimates of their diversity across all plant species from Campbell *et al*. (2018) and the numbers of eudicots and monocots with numbers derived from Evert *et al*. (2013). We linked numbers of species within families extracted from COMPADRE with the number of species per Family listed in The Plant List (2010) for the same families. FitzJohn *et al*. (2014) and Willis (2017) estimate how many plant species in the world are woody, which we compared with the number of trees and shrubs in COMPADRE. We compared the number of epiphytes with estimates from (Zotz, 2013). As above, we used *χ^2^*-tests and post-hoc proportion tests for these comparisons.

#### Conservation status

To characterise potential bias in the conservation statuses of species studied (**H5**), we obtained the IUCN Red List categories (IUCN 2019) for all species in our data set and in The Plant List, using the R package `rredlist` v. 0.4.0 (Chamberlain, 2017). We compared the distribution of COMPADRE species in each Red List category with the corresponding distribution of all species in The Plant List using a *χ^2^*-test.

#### Population growth rates

To assess whether researchers tend to collect demographic data on growing or declining populations and whether researchers tend to study populations that are in a “boom phase” (**H6**), we examined the asymptotic population growth rates (*λ*) calculated from each MPM. We first filtered the data to include only studies that spanned at least five years, were experimentally unmanipulated, and for which *λ* could be calculated (*i.e.*, the MPMs contained no missing values and did not violate ergodicity and irreducibility assumptions (Stott *et al*., 2010)). We then fitted an ordinary least squares regression with *λ* as the response variable and year as the explanatory variable. The slope of this model indicates the temporal trend in *λ*: a negative trend supports our hypothesis that researchers preferentially work on initially flourishing sites where population growth rates then decline over time.

### HOW ARE THE MPMS CONSTRUCTED?

#### Temporal and spatial replication

To explore potential biases in temporal and spatial replication (**H7**), we examined the frequency distributions of study length (years) and the number of spatially distinct populations (as defined by the original article authors) for all species-by-article combinations in COMPADRE.

#### Matrix model dimension

To examine how the MPM dimension chosen by modellers varies systematically among growth form and ecoregion (**H8**), we compared the frequency distribution of the MPM dimension across these variables. As above, we tested for an association statistically using Poisson GLMs.

#### Averaging matrix model elements

In COMPADRE, MPMs (**A**) are defined as the sum of three submatrices, **A** = **U** + **F** + **C**, where **U** represents survival-dependent transitions (*e.g.*, growth, stasis, shrinkage, ageing), **F** describes sexual reproduction, and **C** represents clonal reproduction. We used these three submatrices to address **H9** by characterising the prevalence of averaging across stage-/age-classes. This approach was only possible for matrices without missing values. We assessed the number of consecutive life cycle stages in each MPM that contained averaged rates of survival, reproduction and clonality, using the following approaches:

To estimate the degree of survival averaging, we calculated stage-specific survival probability as the column sums of the **U** matrix. When the survival probabilities across stages were all different, we categorised the MPM as not containing averaged rates (“no averaging”). When several survival probabilities were consecutively identical across up to 50% of stages, we assumed they had been averaged over those stages (“≤50% averaging”). Finally, when more than half of the stages have consecutively identical survival probabilities, we assumed that they had been averaged (“> 50% averaging”).

To estimate the degree of averaging for sexual reproduction, we calculated stage-specific sexual reproduction as the column sums of the **F** matrix. We classified the degree of averaging into three categories in the same way as for survival. Whereas every survival column must sum to >0, reproduction columns may include zero values. These may be before the stage of first reproduction (pre-reproductive) or after (post-reproductive). Post-reproductive stages with **F** = 0 are more likely to exist because researchers did not observe reproduction during fieldwork, rather than due to the method of averaging over multiple life-stages. Our assessments include only ‘reproducing’ stages (i.e., where reproduction >0). We estimated the degree of clonality averaging in the same way, using the **C** matrix.

## Results

### WHEN HAVE THE STUDIES BEEN PUBLISHED?

The proportion of articles focussing on plant demography (**H1**) in the *Journal of Ecology* has increased slightly in the past decades. However, the year-to-year variation is high, and the slope is not significantly different from zero (linear model: slope = 0.034±0.044, *F*_1,21_ = 0.601, *P* = 0.447; Fig. 1).

**Figure 1:**
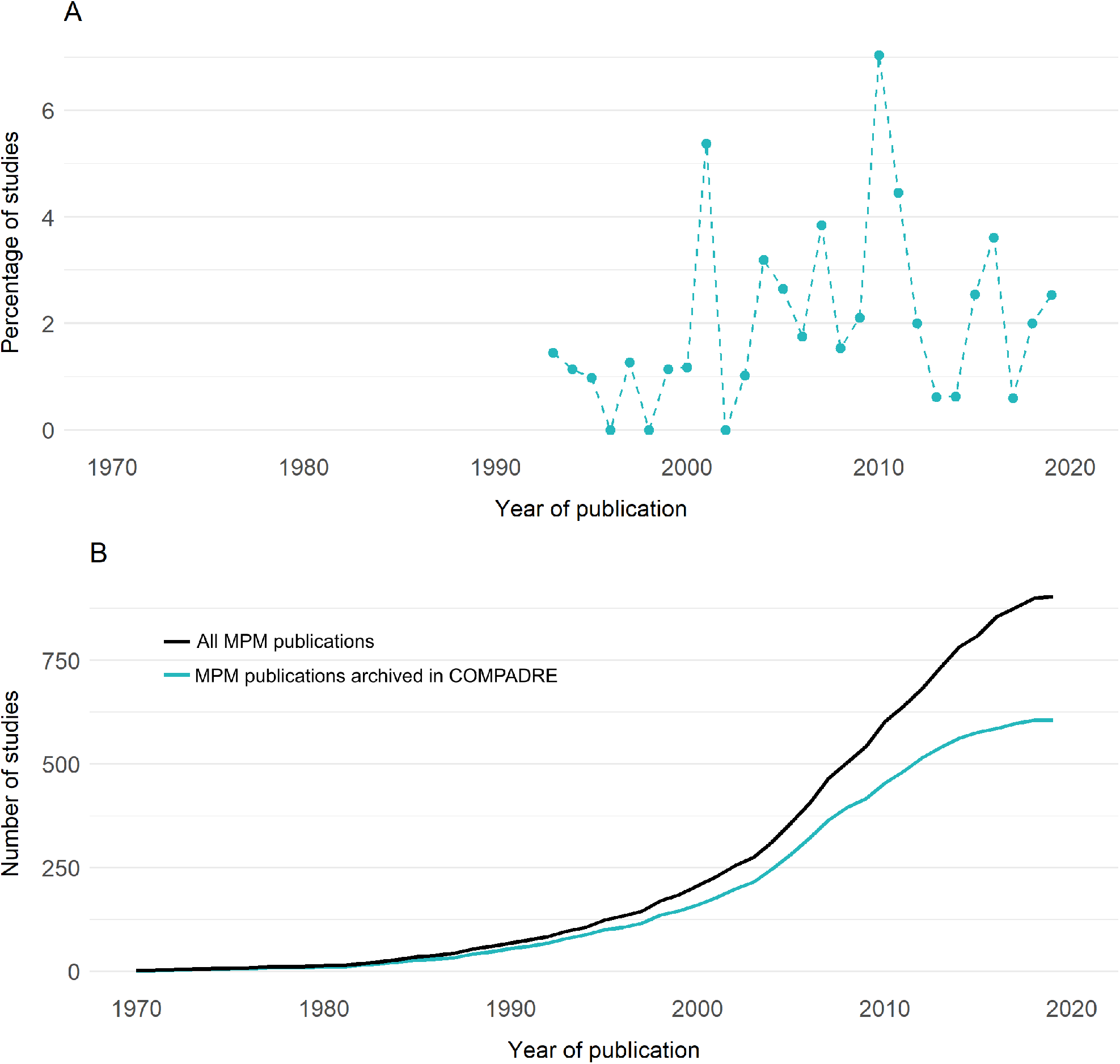
Publication trajectories in plant demography. **(A)** The percentage of articles in a sample of plant ecology literature that use matrix population models (MPMs). Note that the time-series starts in 1993 before which abstracts have been not digitised. **(B)** The cumulative number of MPM-based studies by year, archived in the COMPADRE Plant Matrix Database (at February 2019) (blue line) compared to the estimated cumulative number of all published studies containing plant MPMs (black line).

### WHERE IS THE RESEARCH DONE?

#### Biogeography

As expected (**H2**), geographical bias was obvious, with study density being greatest in Europe (23.67 studies per million km2) and North America (17.56 studies per million km^2^), while Oceania, South American, Asian, and African countries were relatively poorly-represented, with 6.69, 5.10, 1.66, 1.02 studies per million km^2^, respectively (see also Fig. 2B; Fig.S2). Interestingly, a post-hoc analysis showed that the dominance of Europe and North America in the COMPADRE database has increased since the year 2000: 75% of articles post-2000 focussed on these regions compared to 63% pre-2000 (Table S2).

**Figure 2.**
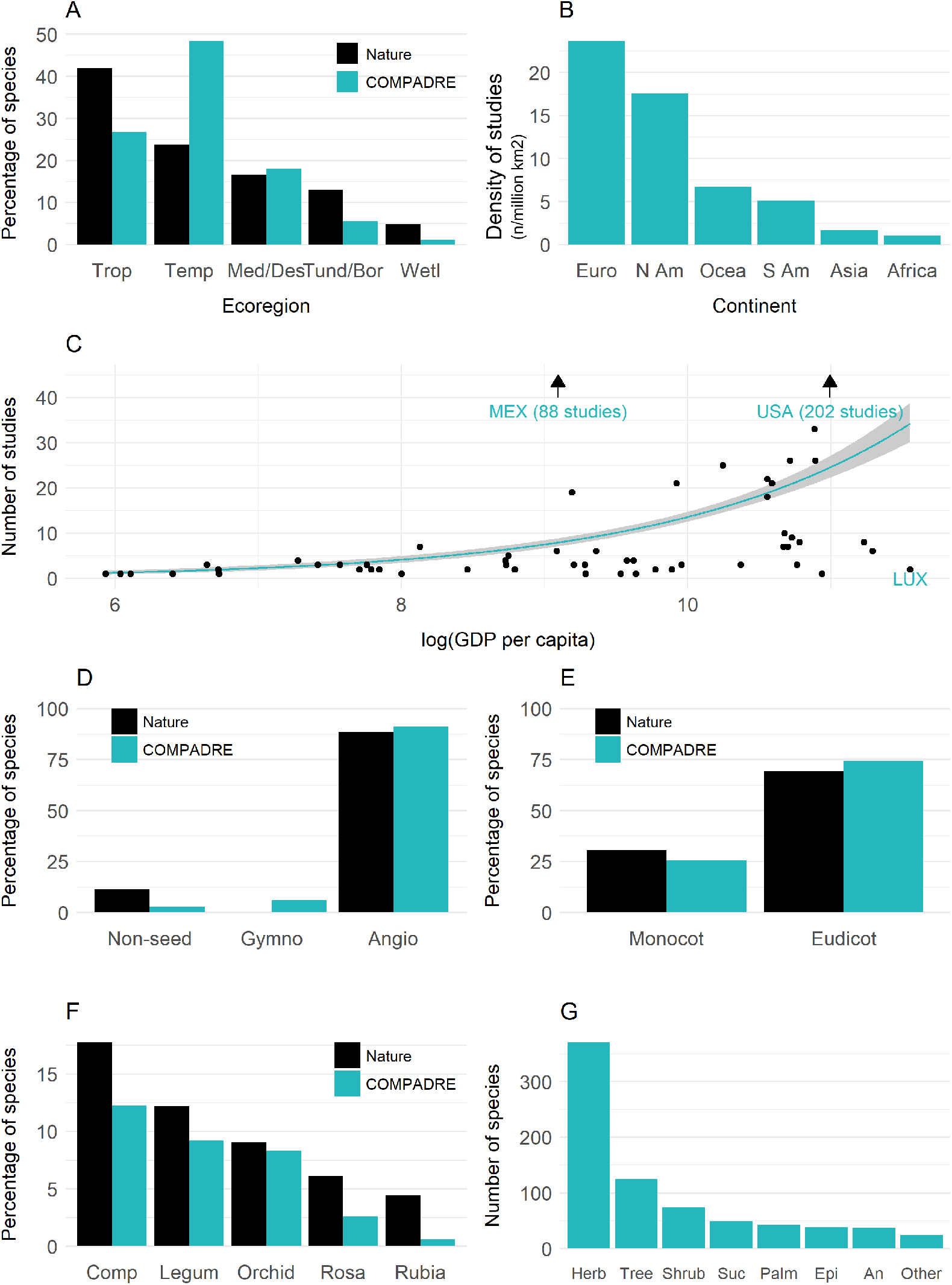
Geographic and taxonomic biases in the COMPADRE Plant Matrix Database. **(A)** The species distribution among ecoregions in COMPADRE compared to the natural distribution (Trop = tropical; temp = temperate; Med/Des = Mediterranean and deserts; Tund/Bor = tundra and boreal regions; Wetl = Wetlands). **(B)** The distribution of plant demography study density across continents. **(C)** The relationship between country per-capita GDP and the number of plant demography studies. The regression line represents a gamma-error GLM, conditioned on countries having at least one plant demography study. **(D)** Comparison of the species distribution among broad categories of angiosperms, gymnosperms and non-seed plants, in COMPADRE and in nature. **(E)** Comparison of the distribution of angiosperm species among monocot and eudicot categories, in COMPADRE and in nature. **(F)** Comparison of the distribution of species among the five largest dicot families, in COMPADRE and in nature (Comp = Compositae; Legum = Leguminosae; Orchid = Orchidaceae; Rosa = Rosaceae; Rubia = Rubiaceae). **(G)** The distribution of species among growth form categories (Herb = herbaceous perennials; Tree = trees; Shrub = shrubs; Suc = succulents; Palm = palms; An = annuals; Epi = epiphytes; Other = includes mosses and ferns).

As hypothesised (**H2**), there was a significant difference between the species distribution in COMPADRE compared to in nature (*χ^2^* -test: *χ^2^* = 327.79, d.f. = 4, *P <*0.001). Indeed, species from temperate ecoregions are significantly over-represented in COMPADRE (48% compared to an estimated 24% of species inhabiting these regions in nature; proportion test: *χ^2^* = 267.71, d.f. = 1, *P <*0.001; Fig. 2A). In contrast, tropical ecoregions are significantly under-represented (27% vs. 42%; proportion test: *χ^2^* = 77.432, d.f. = 1, *P <*0.001). Similar results were apparent for wetlands (1% vs. 5%; proportion test: *χ^2^* = 23.426, d.f. = 1, P <0.001) and tundra and boreal ecoregions (6% vs. 13%; proportion test: *χ^2^* = 40.335, d.f. = 1, *P <*0.001). Species from Mediterranean and desert ecoregions are represented in approximately the same proportion as in nature (18% in COMPADRE *vs*. an estimated 17%: proportion test: *χ^2^* = 1.273, d.f. = 1, *P* = 0.259).

#### Country-specific wealth (GDP)

As expected (**H3**), the numbers of articles per country and per capita GDP are positively correlated (Poisson GLM: Null Deviance = 1494.7, Residual Deviance = 1144.3, d.f. = 1, 54, *P <*0.001, Fig. 2C). Here, 73% of countries (161 out of 222) are not represented in COMPADRE. The United States of America, with 202 research articles, dominates COMPADRE, followed by Mexico with 88 articles, Sweden (33 articles), Australia and Canada (both 26 articles), Spain (25 articles), Japan (22 articles), Czech Republic and the United Kingdom (both 21 articles), and Brazil (19 articles). Interestingly, several of the wealthiest countries are not represented in COMPADRE (*e.g.*, China, Iceland, and Ireland). The positive correlation between the number of studies in COMPADRE and country GDP (Fig. 2C) remains statistically significant even when we remove the two outliers with most studies (USA and Mexico).

### FOR WHICH SPECIES AND POPULATIONS DO WE HAVE DEMOGRAPHIC DATA?

#### Taxonomy

As expected (**H4**), the representation of species in COMPADRE does not well-reflect natural diversity. COMPADRE categorises species as angiosperm (91%), gymnosperm (6%), or “non-seed plants” (3%) which includes ferns, mosses etc. (Fig. 2D). According to Campbell *et al*. (2018), the corresponding figures in nature are 88% (angiosperm), 1% (gymnosperm) and 11% (non-seed plants). Thus, COMPADRE’s taxonomic representativity is different than that found in nature (*χ^2^* test: *χ^2^* = 1282.7, d.f. = 2, *P <*0.001). In fact, COMPADRE over-represents gymnosperms (proportion test: *χ^2^* = 1127.1, d.f. = 1, *P <*0.001) and under-represents the non-seed plants (proportion test: *χ^2^* = 64.39, d.f. = 1, *P <*0.001).

Of COMPADRE’s angiosperms, 74% are eudicots and 26% monocots (Fig 2E), which approximately reflects Evert *et al*.’s (2013) estimate of their natural diversity distribution (69% eudicot, 31% monocot). Our *χ^2^* test nevertheless indicated the COMPADRE distribution was significantly different to the natural distribution (*χ^2^* test: *χ^2^* = 9.79, d.f. = 1, *P* = 0.002).

To better understand the distribution of COMPADRE species across plant families, we examined the five largest eudicot families (according to The Plant List, 2010): Compositae (Asteraceae), Leguminosae (Fabaceae), Orchidaceae, Rosaceae, and Rubiaceae (Fig. 2F). The *χ^2^* test showed that the distributions differed between COMPADRE and in nature patterns (*χ^2^*test: *χ^2^* = 24.161, d.f. = 4, *P <*0.001). However, this difference is driven by the Rubiaceae and Orchidaceae, which are significantly under-represented (Rubiaceae: proportion test: *χ^2^* = 13.616, d.f. = 1, *P <*0.001; Orchidaceae: proportion test: *χ^2^* = 7.0719, d.f. = 1, *P* = 0.008). The other families are fairly proportionately represented in COMPADRE (all *P* >0.05).

Half of the species in COMPADRE are herbaceous perennials (462 of 932 species), and only 26% are woody plants. This figure contrasts with current estimates that 45-48% of the world’s vascular plant species are woody (FitzJohn *et al*., 2014; Willis, 2017).

#### Conservation status

The IUCN has assessed only 29% (n = 220) of the species in COMPADRE. Of these, contrary to our expectations (**H5**), COMPADRE’s content with respect to the Red List status reflects current Red List assessments (*χ^2^* test: *χ^2^* = 4.054, d.f. = 4, *P* = 0.399). Most of COMPADRE’s species are assessed as Least Concern (62%), with the rest falling into one of the threatened categories (Vulnerable, Endangered, or Critically Endangered) (Fig. 3A).

**Figure 3.**
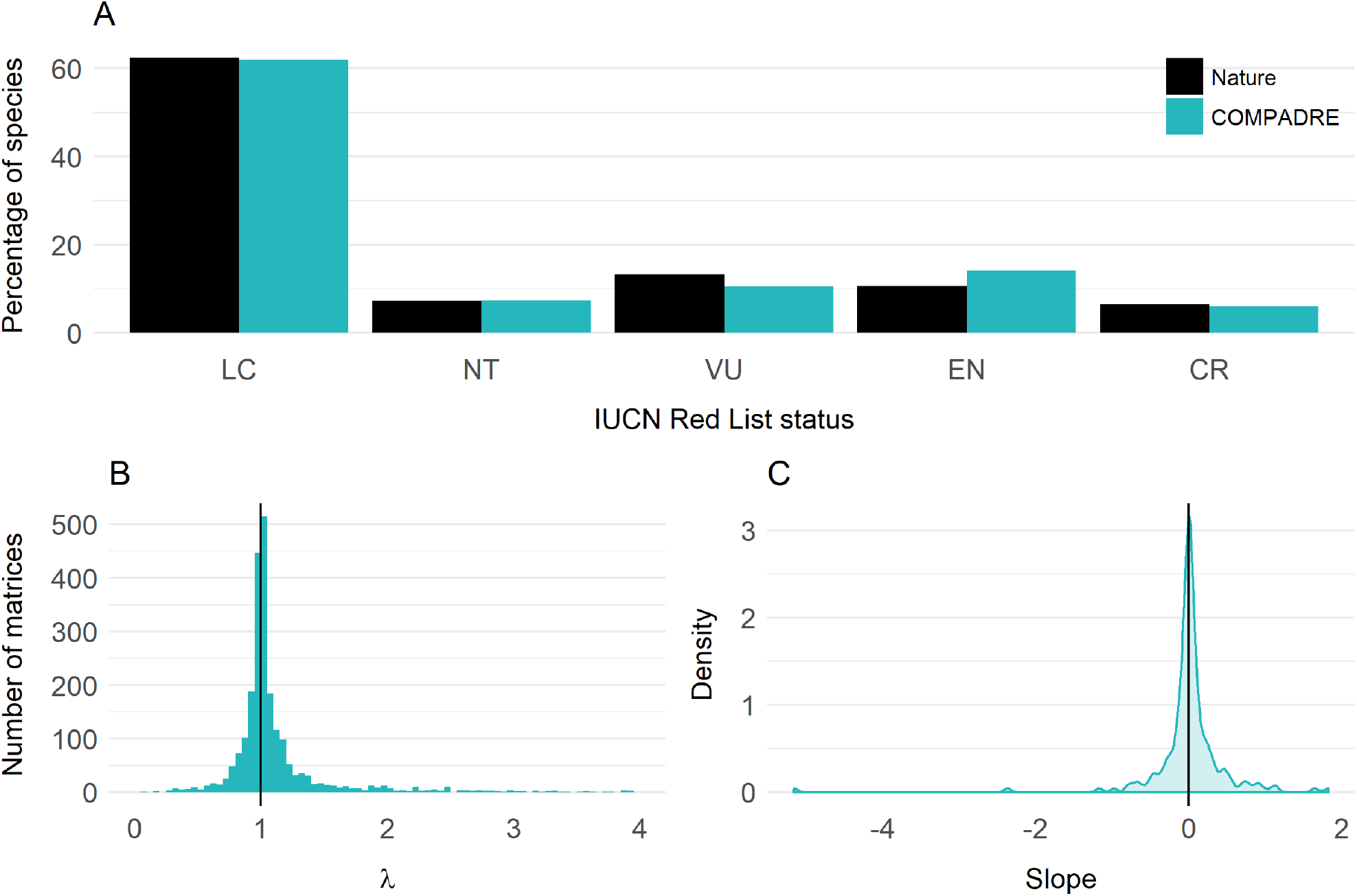
Conservation status and population trend biases in the COMPADRE Plant Matrix Database. **(A)** Comparison of the distribution of species among IUCN Red List conservation status in COMPADRE (blue) and in nature (black) (LC = Least Concern, NT = Near Threatened, VU = Vulnerable, EN = Endangered, CR = Critically Endangered). **(B)** The distribution of population growth rates (*λ*) for MPMs in COMPADRE. The graph is limited to *λ*-values between 0 and 4 to show the interesting area around *λ* = 1 (and *λ* > 4 seems biologically unreasonable and may represent errors). **(C)** The density distribution of the slope of the linear *λ* ∼ year relationship for studies with >5 years of data.

#### Population growth rates

As predicted (**H6**), there is a slight but statistically significant tendency to study growing populations (*i.e.*, *λ* > 1), the effect size is small and driven by the skewed nature of the data (Fig. 3B; two-sided t-test on *λ* = 1: *t* = 10.941, d.f. = 2312, *P* = 0.001). The overall mean value for log *λ* was 0.013 (standard deviation = 0.45). Our analysis of ordinary least-squares regression slopes from the subset of populations with at least a 5-year time-series shows no tendency for researchers to select populations where *λ* is initially high but then decreases, leading to negative slope values (Fig. 3C; t-test: *t* = −0.020, d.f. = 192, *P* = 0.984).

### HOW ARE THE MPMS CONSTRUCTED?

#### Temporal and spatial replication

As expected (**H7**), most studies in COMPADRE are short-term (Fig. 4A). The modal study duration is three years, while the median is four years. The mean is slightly longer (5.48 years), reflecting the skewed nature of the distribution. Some long-term exceptions include studies of the shrub *Cassia nemophila* (Silander, 1983) and the tree *Acer saccharum* (Lin & Augspurger, 2008), which both span 51-years. The study duration varies by ecoregion, with tropical and marine studies tending to be shorter than those from other ecoregions (Fig. S3A). There was minimal variation in study duration among growth forms and, contrary to expectation, trees (a typically long-lived growth form) are not studied for longer periods than typically shorter-lived growth forms (Fig. S4A).

**Figure 4:**
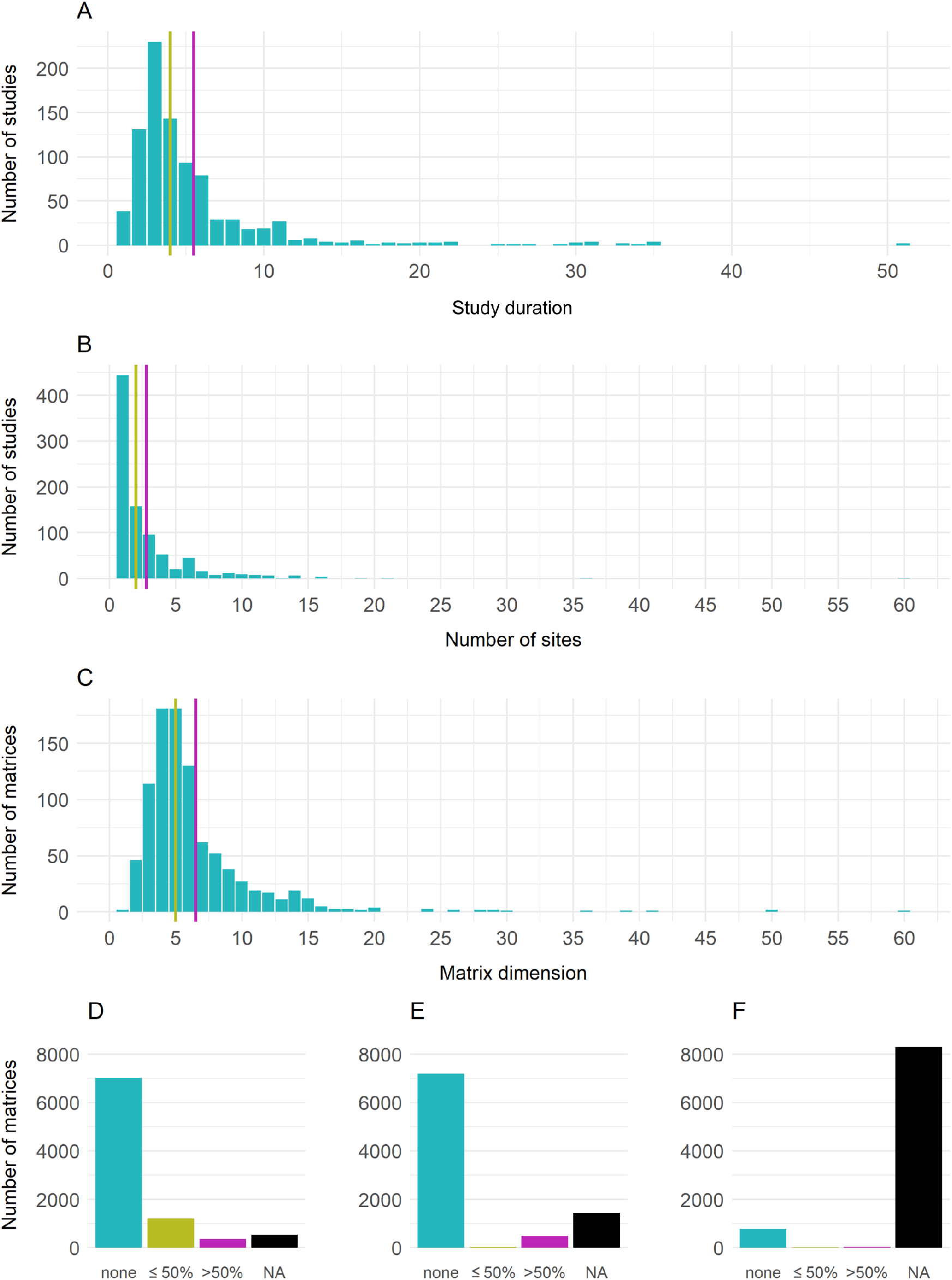
Spatiotemporal replication and MPM construction. **(A)** The distribution of study duration in COMPADRE. **(B)** The distribution of the number of study sites (spatial replication) in COMPADRE. **(C)** The distribution of matrix dimension across MPMs in COMPADRE. The magenta and yellow lines show the mean and median, respectively. **(D, E, F)** Summaries of element averaging in MPM submatrices of **(D)** survival, **(E)** fecundity and **(F)** clonality. *none =* all stages have different survival / fertility / clonality estimates; *≤50% =* apparent averaging with two or more consecutive values are the same, but the number of stages apparent averaging does not exceed 50%; *>50%* = more than half of the stages have the same value.

As expected, most studies have a low degree of spatial replication (Fig 4B). Most studies are carried out at a single site, though the distribution is skewed (mean = 2.82, median = 2, range = 1-60). Interestingly, species studied at four or more sites are mainly herbaceous perennials (57%).

Each MPM is obtained from a different study site or year. Therefore, the number of MPMs present in a study may be an indicator of the range of environmental conditions captured by the study. However, the number of MPMs per study does not vary much among ecoregions (Fig S3A and S3B) or growth form (Fig S4A and B).

#### Matrix model dimension

We expected (**H8**) that the MPM dimension would vary widely and would be greater for long-lived groups like trees. MPM dimension ranges between two and 60 (for *Rhododendron ponticum*, (Travis *et al*., 2011) but is left-skewed (mean = 6.51, median = 5; Fig. 4C). Furthermore, typical matrix dimension varied significantly among ecoregions (ANOVA on log matrix dimension: *F* = 17.967, d.f. = 5 and 884, *P <*0.001) (Fig. S3C), with tropical species tending to have slightly larger matrices than temperate or Mediterranean ones (t-test on log matrix dimension: *t* = 6.855, d.f. = 375.36, *P <*0.001; though this effect is likely driven by large tropical tree matrices). Finally, MPM dimension varies systematically across growth forms (ANOVA on log matrix dimension: *F* = 22.718, d.f. = 6 and 730, *P <*0.001) with tree MPMs tending to be larger (mean = 9.48, median = 8, range = 2-60) than other growth forms (Fig. S4C), thus supporting our initial hypothesis (**H8**).

#### Averaging matrix model elements

Contrary to our expectation (**H9**), researchers do not often appear to average survival rates across life cycle stages when parameterising MPMs. In COMPADRE, 77% of MPMs have no averaging of survival across consecutive stages at all, and only 4% have more than half of their stage-specific survival rates averaged across consecutive stages (Fig. 4D). A similar pattern follows for fecundity: 79% have no averaging and a smaller proportion (5.1%) have over half of their stage/age-specific fecundity rates averaged (Fig. 4E). Fewer than 1% of the MPMs showed averaging in clonality rates (Fig. 4F).

## Discussion

Before analysing large heterogeneous databases, it is essential to understand their potential biases and inconsistencies. Under-representation of particular ecoregions or taxonomic groups may lead to incorrect generalisations if those under-represented regions or groups have distinct demographic behaviour. Improved knowledge of systematic biases in databases like COMPADRE, and a recognition of their impact on inferences, will improve our understanding of the natural world. Researchers should carefully consider potential systematic biases in these large-scale datasets, especially when conducting comparative studies that seek to generalise across disparate taxa and geographic regions.

COMPADRE v.5.0.0. contains data for 746 species, representing 0.002% of the ∼370,000 extant plant species (not including green algae) (The Plant List, 2010). Although COMPADRE covers only a fraction of plant diversity, we show that it contains the majority of published MPM-based plant demographic work (Fig. 1B). Thus, this database is a valid indicator of our knowledge of MPM-based plant demography and reveals demographic knowledge gaps for most species. We note, however, that the vast majority of these studies are published in the English language literature. Amano *et al*. (2016) found that a third of the literature in biodiversity conservation was non-English, and that half provide neither the title nor the abstract in English. About 16% of this corpus is unsearchable using English keywords, thus remaining hidden. Assuming that there is a similar pattern for plant demography literature, some knowledge gaps in the English-speaking research community could undoubtedly be closed by engaging researchers familiar with non-English language literature to assist with contributions to COMPADRE.

### WHEN HAVE THE STUDIES BEEN PUBLISHED?

The number of articles that focus on plant demography has steadily accumulated since the 1970s (Fig. 1B). Our hypothesis that an increasing proportion of plant ecology research would be demography-focussed (**H1**) was supported, based on our survey of articles published in the *Journal of Ecology* (Fig 1A), although the relationship was rather noisy. The downturn in the rate of increase in MPM-related publication in recent years (since about 2015) may be due to the increasingly important role that integral projection models (IPMs, Easterling *et al*., 2000) play in plant demographic research.

### WHERE IS THE RESEARCH DONE?

#### Biogeography

Plant biodiversity is unevenly distributed. Equatorial regions are usually relatively species-rich, with declining biodiversity towards the poles for most plant groups (Gaston, 2000). Biodiversity and endemism hotspots are concentrated in the tropics, on equatorial islands, and in the southern hemisphere (Myers *et al*., 2000; Kier *et al*., 2009). Our finding that ∼73% of demographic studies are carried out in the mainly temperate regions of the western northern hemisphere (Fig. 2A & B) contrasts with those hotspots, thus supporting **H2**. This result is not surprising given similar findings for population dynamics (Amano & Sutherland, 2013; McRae *et al*., 2017), biodiversity time-series (Dornelas *et al*., 2018), and the distribution of ecological study sites (Martin *et al*., 2012). Collectively, these patterns highlight important knowledge gaps for some of the planet’s most threatened ecosystems. For example, sub-Saharan Africa and Southeast Asia have among the least ecological data yet show the most rapid decline of terrestrial ecosystems (MEA, 2005). Given that demographic data is important for assessing extinction risk, *e.g.*, by assessing population trends and population viability analyses (IUCN 2019; Rodrigues, *et al*., 2006), the lack of demographic data is a concerning impediment to species-level conservation. Beyond conservation, this biogeographic bias limits our understanding of global ecological trends and drivers of population dynamics and life history evolution.

#### Country-specific wealth (GDP)

As anticipated (**H3**), the number of demographic articles was positively associated with the per-capita GDP of the country where the work was carried out (Fig. 2C). We expected this because high-GDP countries can invest more in research (van Noorden & Butler, 2019; World Bank, 2019), and researchers tend to conduct research near their home institution for logistical reasons (Coutts *et al*., 2016). The fact that the relationship is relatively loose reflects the international networks and mobility of some researchers who carry out research away from their home institution. Nevertheless, one way to correct this bias is for funding bodies to encourage plant demographic research, and collaboration with researchers, in understudied and threatened regions.

### FOR WHICH SPECIES AND POPULATIONS DO WE HAVE DEMOGRAPHIC DATA?

#### Taxonomy

As expected (**H4**), most of COMPADRE’s species are angiosperms, reflecting the high diversity of this group. However, COMPADRE over-represents gymnosperms and under-represents the non-seed plants. Within the angiosperms, most of COMPADRE’s species are eudicots. Although this reflects the natural distribution of eudicot *vs.* monocot species, we did detect a statistically significant bias towards the study of eudicots. COMPADRE’s species distribution among the major eudicot plant Families approximated the natural distribution, except for Rubiaceae and Orchidaceae, which were under-represented. The pattern for Rubiaceae may be explained by geographical bias since it mainly occurs in the (sub)tropics, which are not well-represented in COMPADRE. As expected (**H4**), we know more about the demography of herbaceous perennials than any other growth form. Trees, which are important both economically (Poore, 2013) and ecologically (Chambers *et al*., 2001), are under-represented, probably due to the logistical difficulties of studying large, long-lived organisms, but perhaps also due to the relatively low tree species diversity in temperate regions.

Although some of these biases may be overcome statistically (*e.g.*, by resampling or rarefaction), the scarcity of demographic data on several growth forms, including ferns, lianas, and bryophytes, drastically reduces our ability to draw general patterns for these growth forms and set them in context with more commonly-studied forms. This issue is particularly troublesome for comparative studies of the evolution of plant life history.

#### Conservation status

Demographic models are an indispensable tool to guide management decisions for threatened species (Norris, 2004). Contrary to our expectation (**H5**) that researchers may collectively focus on threatened species, the distribution of demographic studies in COMPADRE well-approximates the distribution of Red List statuses of plants in general: There is no tendency to favour studies of threatened species. However, we should regard this result with caution because it is based on the subset of ∼25% of COMPADRE species that have been assessed for the IUCN Red List. The true distribution of species among IUCN Red List categories may be quite different, especially since species endemic to biodiversity hotspots are less likely to have been assessed.

#### Population growth rates

There was a slight tendency for researchers to preferentially study growing populations (supporting **H6**) but there was no evidence for a “regression to the mean” effect whereby population growth rates decline along the time-series (contrary to Buckley *et al*., 2010). We initially expected this tendency because we expected researchers to select obviously flourishing populations to avoid the risk and logistical cost of local extinction. The differences between our results and those of Buckley *et al*. (2010) could be due to differences in data or methods. This effect warrants further investigation because biased sampling (*e.g.*, towards growing populations) could lead to incorrect conclusions about population dynamics in comparative research.

### HOW ARE THE MPMS CONSTRUCTED?

#### Temporal and spatial replication

The fact that demographic studies tend to have low spatial and temporal replication supports our original hypothesis (**H7**) and confirms previous findings (Crone *et al*., 2011; Ehrlén *et al*., 2016). Limited spatial replication may affect confidence in inferences made from those models. The geographic distribution of plants varies widely, with some only occurring in specific small areas (*e.g.*, *Iliamna remota* is endemic to the ∼8 hectare Langham Island, Illinois, USA; Swinehart & Jacobs, 1998) and others even spanning continents (*e.g.*, *Plantago major*, Sagar & Harper, 1964). Widely distributed species are likely to experience a greater range of environmental conditions than those with small ranges, and demographic data should ideally be collected in representative parts of this range to understand the species’ demography more fully. Work by Doak & Morris (2010), Wardle *et al*. (2014), and Römer *et al*. (2021) are good examples of such efforts.

The limited temporal extent in most studies is also a concern. Researchers have argued that accurate forecasting of population dynamics typically requires time-series extending well beyond three years, especially because of the demographic impacts of rare extreme weather events (Doak & Morris, 2010; Ehrlén *et al*., 2016; Teller *et al*., 2016; Pérez-Llorca, *et al*., 2018). Given the cost and effort required for long-term research, it is not surprising that the temporal extent of studies in COMPADRE is short. In some settings, researchers could use an alternative space-for-time substitution approach to resolve this problem. The approach enables a rapid accumulation of data representing a large range of environmental conditions allowing the modelling of responses to future climate scenarios without the need for long time-series (Blois *et al*., 2013; Teller *et al*., 2016; Damgaard, 2019; Römer *et al*., 2021). However, the approach assumes that drivers of demographic variation across space are equivalent to those that drive temporal variation, which may not be the case (Pickett, 1989). In any case, the low spatial replication in COMPADRE may currently limit the application of this approach.

#### Matrix model dimension

Researchers constructing MPMs decide an appropriate dimension for their model, based on factors including species life-history (including longevity or life cycle complexity), the study’s purpose, and the amount of data available to parameterise each stage. As expected (**H8**), matrix dimension varies hugely, with a substantial proportion (∼20%) having a low dimension of 3 or less. This low dimension could limit utility in some cases. For example, this is likely to be too low for the derivation of measurements relying on the calculation of age trajectories from stage-based MPMs (Cochran & Ellner, 1992; Caswell, 2001) such as Keyfitz’s entropy (Keyfitz, 1968). Furthermore, other derived metrics, including elasticities (Salguero-Gómez & Plotkin, 2010) and some transient measures (Stott *et al*., 2010), are sensitive to the MPM dimension. Besides influencing individual metrics, the systematic bias in model dimension among growth forms and ecoregion could lead to spurious inferences in multi-species comparative studies if not taken into account.

#### Averaging matrix model elements

MPM-derived metrics of population dynamics such as transient dynamics metrics (Stott *et al*., 2010) and measures of life-history (*e.g.*, survival inequality or entropy) are sensitive to homogeneity among the vital rates of stages because peaks and troughs of survival or mortality in certain life stages might be undetected. Despite this, researchers sometimes have no option but to parameterise MPMs with average vital rates across consecutive stages. We expected this would be common (**H9**), but our results show that plant ecology researchers seldom take this approach. This finding is good news because averaging of reproduction or survival could lead to an underestimation of the effects of temporal variation in the environment underlying the vital rates (Stott *et al*., 2010). This underestimation would be a challenge for applications that are not based on asymptotic properties of the MPM, such as calculations of extinction risk, the stochastic growth rate in population viability analyses, and short-term predictions of population fate (transient dynamics; Stott *et al*., 2011). Thus, inference from MPMs that have been parameterised in this way could lead to misguided management strategies that miss opportunities to influence the population dynamics in the desired way by manipulating vital rates with strong influences on short-term or stochastic population growth. This averaging of vital rates would also problematic for comparative life-history research, and in particular, work that relies on age-from-stage methods (Cochran & Ellner, 1992; Caswell, 2001) to calculate demographic trajectories and derived measures. We, therefore, encourage researchers to, wherever possible, avoid parameterisation using averaging over consecutive stages.

## Conclusions

Our current knowledge of global plant demography is based on geographically biased data heavily focused on herbaceous perennials, leaving important knowledge gaps. Demographic studies are constrained by funding, with the temporal length reflecting the typical grant and PhD tenure, with most work concentrated in wealthy countries. We did not find significant bias in conservation status or population growth rate, which indicates that researchers do not focus on species of conservation concern nor growing or shrinking populations. To close the aforementioned knowledge gaps and better understand generalities in life-history strategy and population dynamics, research targeting neglected growth forms and ecoregions is desirable, as is increased spatial and temporal replication within species. Furthermore, an improved understanding of the impact of these biases on model predictions and methodological developments to account for known biases would be helpful.

## Supporting information

Appendix S2: Analysis code

## Acknowledgements

We thank the researchers that have contributed data to the COMPADRE Plant Matrix Database. We also thank T. Knight and attendees of the German Centre for Integrative Biodiversity Research (iDiv) sApropos (Analysis of PROjections of POpulationS) workshops in Leipzig led by RS-G, for valuable discussions on this topic. GR was supported to attend the sAPROPOS workshops by German Centre for Integrative Biodiversity Research (iDiv), ORJ was supported by the Danish Council for Independent Research (grant number DFF - 6108-00467). RS-G was also supported by NERC-IRF NE/M018458/1.

## Author’s contributions

GR, RS-G and ORJ conceived the ideas and designed methodology; ORJ and RS-G provided additional data; GR, ORJ and IS analysed the data; GR led the writing of the manuscript. All authors contributed to manuscript writing and gave final approval for publication.

## Data availability

Most of the used data is open access and can be downloaded on the following webpages: The COMPADRE Plant Matrix Database: http://www.compadre-db.org (Salguero-Gómez *et al*., 2015); IUCN Red List data: http://www.iucnredlist.org (IUCN, 2019); The Plant List: http://www.theplantlist.org (The Plant List, 2010); GDP data was obtained from the World Bank: https://data.worldbank.org/indicator/NY.GDP.PCAP.CD (World Bank, 2018). Code for the analyses is included in the supplementary information.

## Supporting information

Appendix S1: Supplementary figures and tables

Appendix S2: Analysis code

## References

Amano, T., González-Varo, J. P., & Sutherland, W. J. (2016). Languages are still a major barrier to global science. PLoS Biology, 14(12), e2000933.

Amano, T., & Sutherland, W. J. (2013). Four barriers to the global understanding of biodiversity conservation: Wealth, language, geographical location and security. Proceedings of the Royal Society Series B. Biological Sciences, 280(1756), 20122649.

Baudisch, A. (2011). The pace and shape of ageing. Methods in Ecology and Evolution, 2(4), 375–382.

Baudisch, A., & Stott, I. (2019). A pace and shape perspective on fertility. Methods in Ecology and Evolution, 10(11), 1941–1951.

Blois, J. L., Williams, J. W., Fitzpatrick, M. C., Jackson, S. T., & Ferrier, S. (2013). Space can substitute for time in predicting climate-change effects on biodiversity. Proceedings of the National Academy of Sciences, 110(23), 9374–9379.

Buckley, Y. M., Ramula, S., Blomberg, S. P., Burns, J. H., Crone, E. E., Ehrlén, J., … Wardle, G. M. (2010). Causes and consequences of variation in plant population growth rate: A synthesis of matrix population models in a phylogenetic context. Ecology Letters, 13(9), 1182–1197.

Buhler, D. D., Hartzler, R. G., & Forcella, F. (1997). Implications of weed seedbank dynamics to weed management. Weed Science, 45(3), 329–336.

Bullock, J. M., White, S. M., Prudhomme, C., Tansey, C., Perea, R., & Hooftman, D. A. P. (2012). Modelling spread of British wind-dispersed plants under future wind speeds in a changing climate. Journal of Ecology, 100(1), 104–115.

Burns, J. H. (2008). Demographic performance predicts invasiveness of species in the Commelinaceae under high-nutrient conditions. Ecological Applications, 18(2), 335–346.

Burns, J. H., Pardini, E. A., Schutzenhofer, M. R., Chung, Y. A., Seidler, K. J., & Knight, T. M. (2013). Greater sexual reproduction contributes to differences in demography of invasive plants and their noninvasive relatives. Ecology, 94(5), 995–1004.

Campbell, N. A., Urry, L. A., Cain, M. L., Wasserman, S. A., Minorsky, P. V., & Reece, J. B. (2018). Biology: A global approach (Eleventh edition, global edition). Boston: Pearson.

Caswell, H. (2001). Matrix population models: Construction, analysis, and interpretation. (2nd ed.). Sunderland, Mass.: Sinauer Associates.

Cavalier-Smith T (1981). Eukaryote kingdoms: seven or nine? Bio Systems. 14 (3–4): 461–81.

Chamberlain, S. (2017). rredlist: ‘IUCN’ Red List Client: R package version 0.4.0. Retrieved from https://CRAN.R-project.org/package=rredlist

Chambers, J. Q., Higuchi, N., Tribuzy, E. S., & Trumbore, S. E. (2001). Carbon sink for a century. Nature, 410(6827), 429.

Cochran, M. E., & Ellner, S. (1992). Simple methods for calculating age-based life history parameters for stage-structured populations. Ecological Monographs, 62, 345–364.

Colautti, R. I., & Barrett, S. C. H. (2013). Rapid adaptation to climate facilitates range expansion of an invasive plant. Science (New York, N.Y.), 342(6156), 364–366.

Conlisk, E., Castanha, C., Germino, M. J., Veblen, T. T., Smith, J. M., & Kueppers, L. M. (2017). Declines in low-elevation subalpine tree populations outpace growth in high-elevation populations with warming. Journal of Ecology, 105(5), 1347–1357.

Coutts, S. R., Salguero-Gómez, R., Csergő, A. M., & Buckley, Y. M. (2016). Extrapolating demography with climate, proximity and phylogeny: Approach with caution. Ecology Letters, 19(12), 1429–1438.

Crone, E. E., Menges, E. S., Ellis, M. M., Bell, T., Bierzychudek, P., Ehrlén, J., … Williams, J. L. (2011). How do plant ecologists use matrix population models? Ecology Letters, 14(1), 1–8.

Dalgleish, H. J., Koons, D. N., & Adler, P. B. (2010). Can life-history traits predict the response of forb populations to changes in climate variability? Journal of Ecology, 98(1), 209–217.

Damgaard, C. (2019). A critique of the space-for-time substitution practice in community ecology. Trends in Ecology & Evolution, 34(5), 416–421.

Doak, D. F., & Morris, W. F. (2010). Demographic compensation and tipping points in climate-induced range shifts. Nature, 467(7318), 959–962.

Dornelas, M., Antão, L. H., Moyes, F., Bates, A. E., Magurran, A. E., Adam, D., … Hickler, T. (2018). Biotime: A database of biodiversity time series for the Anthropocene. Global Ecology and Biogeography, 27(7), 760–786.

Easterling, M. R., Ellner, S. P., & Dixon, P. M. (2000). Size-specific sensitivity: Applying a new structured population model. Ecology, 81(3), 694–708.

Ehrlén, J., Morris, W. F., Euler, T. von, & Dahlgren, J. P. (2016). Advancing environmentally explicit structured population models of plants. Journal of Ecology, 104(2), 292–305.

Eriksson, O., Cousins, S. A.O., & Bruun, H. H. (2002). Land-use history and fragmentation of traditionally managed grasslands in Scandinavia. Journal of Vegetation Science, 13(5), 743–748.

Evert, R. F., Raven, P. H. B. o. p., & Eichhorn, S. E. (2013). Raven biology of plants (Eighth edition). New York: W.H. Freeman and Company Publishers.

FitzJohn, R. G., Pennell, M. W., Zanne, A. E., Stevens, P. F., Tank, D. C., & Cornwell, W. K. (2014). How much of the world is woody? Journal of Ecology, 102(5), 1266–1272.

Fowler, N. L., Overath, R. D., & Pease, C. M. (2006). Detection of density dependence requires density manipulations and calculation of lambda. Ecology, 87(3), 655–664.

Franco, M., & Silvertown, J. (1996). Life history variation in plants: an exploration of the fast-slow continuum hypothesis. Philosophical Transactions of the Royal Society of London. Series B, Biological Sciences, 351(1345), 1341–1348.

Gaillard, J.-M., Yoccoz, N. G., Lebreton, J.-D., Bonenfant, C., Devillard, S., Loison, A., … Allaine, D. (2005). Generation time: A reliable metric to measure life-history variation among mammalian populations. American Naturalist, 166(1), 119–23; discussion 124-8.

Gaston, K. J. (2000). Global patterns in biodiversity. Nature, 405(6783), 220–227.

GBIF: The Global Biodiversity Information Facility. What is GBIF?. Retrieved from https://www.gbif.org/what-is-gbif

Grime, J. P. (1974). Vegetation classification by reference to strategies. Nature, 250(5461), 26–31.

Gunderson, D. R. (1980). Using r-K selection theory to predict natural mortality. Canadian Journal of Fisheries and Aquatic Sciences, 37(12), 2266–2271.

Hansen, M. J., & Wilson, S. D. (2006). Is management of an invasive grass *Agropyron cristatum* contingent on environmental variation? Journal of Applied Ecology, 43(2), 269–280.

Albert, M.J., Giménez Benavides, L., Domínguez Lozano, F. & Escudero, A. (Ed.) (2009). Poblaciones en peligro: Viabilidad demográfica de la flora vascular amenazada de España: Populations in peril: Demographic viability of threatened Spanish vascular flora. Madrid: Dirección General de Medio Natural y Política Forestal.

IUCN 2019. The IUCN Red List of Threatened Species. Version 2019-2. Retrieved from http://www.iucnredlist.org

Kattge, J., Díaz, S., Lavorel, S., Prentice, I. C., Leadley, P., Bönisch, G., … Wirth, C. (2011). TRY - a global database of plant traits. Global Change Biology, 17(9), 2905–2935.

Keyfitz, N. (1968). Introduction to the mathematics of population. Addison-Wesley series in behavioral science: quantitative methods. Reading, Mass.: Addison-Wesley Pub. Co.

Kier, G., Kreft, H., Lee, T. M., Jetz, W., Ibisch, P. L., Nowicki, C., … Barthlott, W. (2009). A global assessment of endemism and species richness across island and mainland regions. Proceedings of the National Academy of Sciences, 106(23), 9322–9327.

Kier, G., Mutke, J., Dinerstein, E., Ricketts, T. H., Küper, W., Kreft, H., & Barthlott, W. (2005). Global patterns of plant diversity and floristic knowledge. Journal of Biogeography, 32(7), 1107–1116.

Lefkovitch, L. P. (1965). The study of population growth in organisms grouped by stages. Biometrics, 21(1), 1.

Leslie, P. H. (1945). On the Use of Matrices in Certain Population Mathematics. Biometrika, 33(3), 183.

Lin, Y., & Augspurger, C. K. (2008). Long-term spatial dynamics of *Acer saccharum* during a population explosion in an old-growth remnant forest in Illinois. Forest Ecology and Management, 256(5), 922–928.

Loh, J., Green, R. E., Ricketts, T., Lamoreux, J., Jenkins, M., Kapos, V., & Randers, J. (2005). The Living Planet Index: Using species population time series to track trends in biodiversity. Philosophical Transactions of the Royal Society of London. Series B, Biological Sciences, 360(1454), 289–295.

MacArthur, R. H., & Wilson, E. O. (1967). The theory of island biogeography. Princeton landmarks in biology. Princeton, N.J., Oxford: Princeton University Press.

Martin, L. J., Blossey, B., & Ellis, E. (2012). Mapping where ecologists work: biases in the global distribution of terrestrial ecological observations. Frontiers in Ecology and the Environment, 10(4), 195–201.

McRae, L., Deinet, S., & Freeman, R. (2017). The diversity-weighted living planet index: controlling for taxonomic bias in a global biodiversity indicator. PloS One, 12(1), e0169156.

MEA (2005). Ecosystems and human well-being: Biodiversity synthesis. Washington DC: World Resources Institute.

Morris, W. F., & Doak, D. F. (2002). Quantitative conservation biology: Theory and practice of population viability analysis. New York: W. H. Freeman; Basingstoke: Palgrave Macmillan.

Morris, W. F., Pfister, C. A., Tuljapurkar, S., Haridas, C. V., Boggs, C. L., Boyce, M. S., … Menges, E. S. (2008). Longevity can buffer plant and animal populations against changing climatic variability. Ecology, 89(1), 19–25.

Myers, N., Mittermeier, R. A., Mittermeier, C. G., da Fonseca, G. A., & Kent, J. (2000). Biodiversity hotspots for conservation priorities. Nature, 403(6772), 853–858.

NERC Centre for Population Biology (2010). The Global Population Dynamics Database Version 2. Retrieved from http://www.sw.ic.ac.uk/cpb/cpb/gpdd.html.

Norris, K. E. N. (2004). Managing threatened species: the ecological toolbox, evolutionary theory and declining-population paradigm. Journal of Applied Ecology, 41(3), 413–426.

Olson, D. M., Dinerstein, E., Wikramanayake, E. D., Burgess, N. D., Powell, G. V. N., Underwood, E. C., … Kassem, K. R. (2001). Terrestrial ecoregions of the world: a new map of life on earth. BioScience, 51(11), 933.

Pérez-Llorca, M., Fenollosa, E., Salguero-Gómez, R., & Munné-Bosch, S. (2018). What is the minimal optimal sample size for plant ecophysiological studies? Plant Physiology, 178(3), 953–955.

Pickett, S. T. A. (1989). Space-for-Time Substitution as an Alternative to Long-Term Studies: In: Gene E. Likens (Ed.), Long-Term Studies in Ecology: Approaches and Alternatives. New York NY: Springer New York.

Poore, D. (2013). No Timber Without Trees: Routledge.

Poorter, L., Zuidema, P. A., Peña-Claros, M., & Boot, R. G. A. (2005). A monocarpic tree species in a polycarpic world: how can *Tachigali vasquezii* maintain itself so successfully in a tropical rain forest community? Journal of Ecology, 93(2), 268–278.

R Core Team (2021). R: A language and environment for statistical computing. R Foundation for Statistical Computing, Vienna, Austria. URL https://www.R-project.org/

Ramage, M. H., Burridge, H., Busse-Wicher, M., Fereday, G., Reynolds, T., Shah, D. U., … Scherman, O. (2017). The wood from the trees: The use of timber in construction. Renewable and Sustainable Energy Reviews, 68, 333–359.

Raunkiær, C. (1907). Planterigets Livsformer og deres Betydning for Geografien.: Med 77 figurer i teksten. København and Kristiania: Gyldendalske Boghandel - Nordisk Forlag.

Raunkiær, C. (1934). The Life Forms of Plants and Statistical Plant Geography, being the collected papers of C. Raunkiær.: Translated by H. Gilbert-Carter, A. Fausbøll, and A. G. Tansley. Oxford University Press, Oxford. Reprinted 1978 (ed. by Frank N. Egerton). History of ecology. New York: Arno Press.

Rodrigues, A. S. L., Pilgrim, J. D., Lamoreux, J. F., Hoffmann, M., & Brooks, T. M. (2006). The value of the IUCN Red List for conservation. Trends in Ecology & Evolution, 21(2), 71–76.

Römer, G., Christiansen, D. M., de Buhr, H., Hylander, K., Jones, O. R., Merinero, S., Reitzel, K., Ehrlén, J., and Dahlgren, J. P. (2021). Drivers of large-scale spatial demographic variation in a perennial plant. Ecosphere 12(1). e03356.

Rueda-Cediel, P., Brain, R., Galic, N., & Forbes, V. (2019). Comparative analysis of plant demographic traits across species of different conservation concern: Implications for pesticide risk assessment. Environmental Toxicology and Chemistry, 38(9), 2043–2052.

Sagar, G. R., & Harper, J. L. (1964). Plantago major L., P. media L. and P. lanceolata L. Journal of Ecology, 52(1), 189.

Salguero-Gómez, R. (2017). Applications of the fast-slow continuum & reproductive strategy framework of plant life histories. New Phytologist, 4, 1618–1624

Salguero-Gómez, R., & Plotkin, J. B. (2010). Matrix dimensions bias demographic inferences: Implications for comparative plant demography. American Naturalist, 176(6), 710–722.

Salguero-Gómez, R., Jones, O. R., Archer, C. R., Buckley, Y. M., Che-Castaldo, J., Caswell, H., … Vaupel, J. W. (2015). The COMPADRE Plant Matrix Database: an open online repository for plant demography. Journal of Ecology, 103(1), 202–218.

Salguero-Gómez, R., Jones, O. R., Jongejans, E., Blomberg, S. P., Hodgson, D. J., Mbeau-Ache, C., … Buckley, Y. M. (2016). Fast-slow continuum and reproductive strategies structure plant life-history variation worldwide. Proceedings of the National Academy of Sciences, 113(1), 230–235.

Salguero-Gómez, R., & Kroon, H. de (2010). Matrix projection models meet variation in the real world. Journal of Ecology, 98(2), 250–254.

Silander, J. A. (1983). Demographic variation in the Australian desert cassia under grazing pressure. Oecologia, 60(2), 227–233.

Silva, J. F., Raventos, J., Caswell, H., & Trevisan, M. C. (1991). Population responses to fire in a tropical savanna grass, *Andropogon semiberbis*: A matrix model approach. Journal of Ecology, 79, 345–356.

Silvertown, J., Franco, M., & McConway, K. (1992). A demographic interpretation of Grime’s triangle. Functional Ecology, 6(2), 130.

Silvertown, J., Franco, M., Pisanty, I., & Mendoza, A. (1993). Comparative plant demography--relative importance of life-cycle components to the finite rate of increase in woody and herbaceous perennials. Journal of Ecology, 81(3), 465.

Staerk, J., Conde, D. A., Ronget, V., Lemaitre, J.-F., Gaillard, J.-M., & Colchero, F. (2019). Performance of generation time approximations for extinction risk assessments. Journal of Applied Ecology, 56(6), 1436–1446.

Stearns, S. C. (1992). The evolution of life histories. Oxford University Press, Oxford. UK.

Stott, I., Franco, M., Carslake, D., Townley, S., & Hodgson, D. J. (2010). Boom or bust? A comparative analysis of transient population dynamics in plants. Journal of Ecology, 98(2), 302–311.

Stott, I., Townley, S., Carslake, D., & Hodgson, D. J. (2010). On reducibility and ergodicity of population projection matrix models. Methods in Ecology and Evolution, 1: 242–252.

Stott, I., Townley, S., & Hodgson, D. J. (2011). A framework for studying transient dynamics of population projection matrix models. Ecology Letters, 14, 959–970.

Swinehart, A. L., & Jacobs, M. E. (1998). Rediscovery, status and preservation of the endangered Kankakee Globe Mallow (*Iliamna remota*) in Indiana. Rhodora, 100, 82–87.

Teller, B. J., Adler, P. B., Edwards, C. B., Hooker, G., & Ellner, S. P. (2016). Linking demography with drivers: climate and competition. Methods in Ecology and Evolution, 7(2), 171–183.

The Plant List (2010). The Plant List: A working list of all plant species. Version 1.1. Retrieved from http://www.theplantlist.org

Travis, J. M. J., Harris, C. M., Park, K. J., & Bullock, J. M. (2011). Improving prediction and management of range expansions by combining analytical and individual-based modelling approaches. Methods in Ecology and Evolution, 2(5), 477–488.

van Noorden, R., & Butler, D. (2019). Science in Europe: By the numbers. Nature, 569(7757), 470–471.

Verwer, C., Peña-Claros, M., van der Staak, D., Ohlson-Kiehn, K., & Sterck, F. J. (2008). Silviculture enhances the recovery of overexploited mahogany *Swietenia macrophylla*. Journal of Applied Ecology, 45(6), 1770–1779.

Wardle, G. M., Buckley, Y. M., & PlantPopNet steering group (2014). PlantPopNet: A Spatially Distributed Model System for Population Ecology (Ecology Society of Australia 2014 Annual Conference). Alice Springs NT, Australia.

Willis, K. J. (Ed.) (2017). State of the world’s plants, 2017. [Richmond, Surrey, UK]: Royal Botanic Gardens Kew.

World Bank (2018, October 18). World Development Indicators: GDP per capita (current US$). Retrieved from http://databank.worldbank.org/data/reports.aspx?source=2&type=metadata&series=NY.GDP.PCAP.CD#

World Bank (2019). Research and development expenditure (% of GDP). Retrieved from https://data.worldbank.org/indicator/gb.xpd.rsdv.gd.zs?view=map

Wright, I. J., Reich, P. B., Westoby, M., Ackerly, D. D., Baruch, Z., Bongers, F., … Villar, R. (2004). The worldwide leaf economics spectrum. Nature, 428(6985), 821–827.

Zotz, G. (2013). The systematic distribution of vascular epiphytes - a critical update. Botanical Journal of the Linnean Society, 171(3), 453–481.

Zuidema, P. A., & Boot, R. G. A. (2002). Demography of the Brazil nut tree (*Bertholletia excelsa*) in the Bolivian Amazon: impact of seed extraction on recruitment and population dynamics. Journal of Tropical Ecology, 18(1), 1–31.

Zuidema, P. A., Brienen, R. J. W., During, H. J., & Güneralp, B. (2009). Do persistently fast-growing juveniles contribute disproportionately to population growth? A new analysis tool for matrix models and its application to rainforest trees. American Naturalist, 174(5), 709–719.

Zuidema, P. A., Kroon, H. de, & Werger, M. J. A. (2007). Testing sustainability by prospective and retrospective demographic analyses: evaluation for palm leaf harvest. Ecological Applications, 17(1), 118–128

Zuidema, P. A. (2000). Demography of exploited tree species in the Bolivian Amazon. PROMAB scientific series: Vol. 2. Utrecht: PROMAB.

