## Appendix S1: Supplementary figures and tables for "Plant demographic knowledge is biased towards short-term studies of temperate-region herbaceous perennials"

Table S1. The terrestrial biomes identified by Olson *et al*., (2001) and the associated abbreviations used in the COMPADRE Plant Matrix Database (see digitization protocol, https://jonesor.github.io/CompadreGuides/). We examined the representativity of studies across different biomes by collapsing Olson's 14 categories into five ecoregions to ensure statistically adequate sample sizes.

| **Olson biome** | **COMPADRE abbreviation** | **Collapsed ecoregion name** |
|---|---|---|
| Tropical and subtropical moist broadleaf forests | TMB | Tropical |
| Tropical and subtropical dry broadleaf forests | TDB | |
| Tropical and subtropical coniferous forests | TSC | |
| Tropical and subtropical grasslands, savannas and shrublands | TGV | |
| Temperate broadleaf and mixed forests | TBM | Temperate |
| Temperate coniferous forests | TCF | |
| Temperate grasslands, savannas, and shrublands | TGS | |
| Mediterranean forests, woodlands and scrubs | MED | Mediterranean/desert |
| Deserts and xeric shrublands | DES | |
| Boreal forests/ taiga | BOR | Tundra/boreal |
| Tundra | TUN | |
| Montane grasslands and shrublands | MON | |
| Mangroves | MAN | Wetland |
| Flooded grasslands and savannas | FGS | |

Table S2. The distribution of studies in COMPADRE across continents. The columns show the overall distribution, the distribution pre-2000, and the distribution from 2000 onwards. The proportion of studies focussed on North America and Europe increased after 2000, while the proportion of studies focussed on other continents decreased.

| **Continent** | **Overall %** | **% <2000 studies** | **% 2000 +** |
|---|---|---|---|
| North America | 46.8 | 41.9 | 47.8 |
| Europe | 26.0 | 21.4 | 27.6 |
| South America | 9.8 | 10.0 | 9.8 |
| Asia | 8.0 | 11.9 | 6.9 |
| Oceania | 6.1 | 9.0 | 5.3 |
| Africa | 3.3 | 5.7 | 2.7 |

**A**

$$\begin{bmatrix} 0.4 & 0.0 & 0.0 & 0.0 & 0.0 & 0.0 \\ 0.3 & 0.2 & 0.0 & 0.0 & 0.0 & 0.0 \\ 0.1 & 0.3 & 0.2 & 0.0 & 0.0 & 0.0 \\ 0.0 & 0.2 & 0.1 & 0.1 & 0.0 & 0.0 \\ 0.0 & 0.0 & 0.5 & 0.2 & 0.1 & 0.0 \\ 0.0 & 0.0 & 0.0 & 0.6 & 0.7 & 0.7 \end{bmatrix}$$

0.8 0.7 0.8 0.9 0.8 0.7

**C**

$$\begin{bmatrix} 0.4 & 0.0 & 0.0 & 0.0 & 0.0 & 0.0 \\ 0.3 & 0.2 & 0.0 & 0.0 & 0.0 & 0.0 \\ 0.1 & 0.3 & 0.2 & 0.0 & 0.0 & 0.0 \\ 0.0 & 0.2 & 0.1 & 0.1 & 0.0 & 0.0 \\ 0.0 & 0.0 & 0.5 & 0.2 & 0.1 & 0.0 \\ 0.0 & 0.0 & 0.0 & 0.5 & 0.7 & 0.7 \end{bmatrix}$$

0.8 0.7 0.8 0.8 0.8 0.7

**B**

$$\begin{bmatrix} 0.4 & 0.0 & 0.0 & 0.0 & 0.0 & 0.0 \\ 0.3 & 0.2 & 0.0 & 0.0 & 0.0 & 0.0 \\ 0.1 & 0.3 & 0.2 & 0.0 & 0.0 & 0.0 \\ 0.0 & 0.2 & 0.2 & 0.1 & 0.0 & 0.0 \\ 0.0 & 0.0 & 0.5 & 0.2 & 0.1 & 0.0 \\ 0.0 & 0.0 & 0.0 & 0.6 & 0.7 & 0.7 \end{bmatrix}$$

0.8 0.7 0.9 0.9 0.8 0.7

**D**

$$\begin{bmatrix} 0.4 & 0.0 & 0.0 & 0.0 & 0.0 & 0.0 \\ 0.3 & 0.2 & 0.0 & 0.0 & 0.0 & 0.0 \\ 0.1 & 0.3 & 0.2 & 0.0 & 0.0 & 0.0 \\ 0.0 & 0.2 & 0.1 & 0.1 & 0.0 & 0.0 \\ 0.0 & 0.0 & 0.5 & 0.2 & 0.1 & 0.1 \\ 0.0 & 0.0 & 0.0 & 0.5 & 0.7 & 0.7 \end{bmatrix}$$

0.8 0.7 0.8 0.8 0.8 0.8

**E**

$$\begin{bmatrix} 0.4 & 0.0 & 0.0 & 0.0 & 0.0 & 0.0 \\ 0.3 & 0.2 & 0.0 & 0.0 & 0.0 & 0.0 \\ 0.1 & 0.3 & 0.2 & 0.0 & 0.0 & 0.0 \\ 0.0 & 0.3 & 0.1 & 0.1 & 0.0 & 0.0 \\ 0.0 & 0.0 & 0.5 & 0.2 & 0.1 & 0.1 \\ 0.0 & 0.0 & 0.0 & 0.6 & 0.7 & 0.7 \end{bmatrix}$$

0.8 0.8 0.8 0.9 0.8 0.8

Figure S1. Five examples to illustrate MPM element averaging survival over different stages. **(A)** All column sums are different, and therefore no averaging is assumed. **(B)** and **(C)** show matrices where up to 50% of the column sums are the same, therefore we assume that there has been some averaging. Matrix **(D)** shows assumed averaging in >50% of consecutive columns. In matrix **(E)** >50% of the column sums are the same, but since they are not in consecutive columns, this matrix falls in the category with up to 50% averaging.

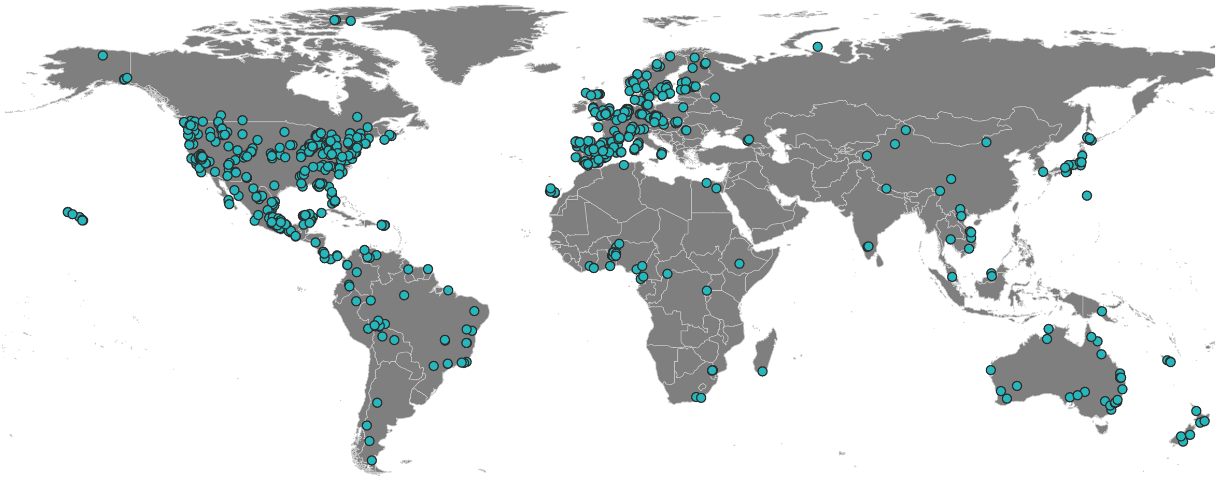

Figure S2. Spatial distribution of studies in COMPADRE. Each point represents a studied population. Most of the studies are conducted in North America and Europe. Sixteen studies are not represented here because their GPS coordinates are either not available or because they were carried out in laboratory conditions.

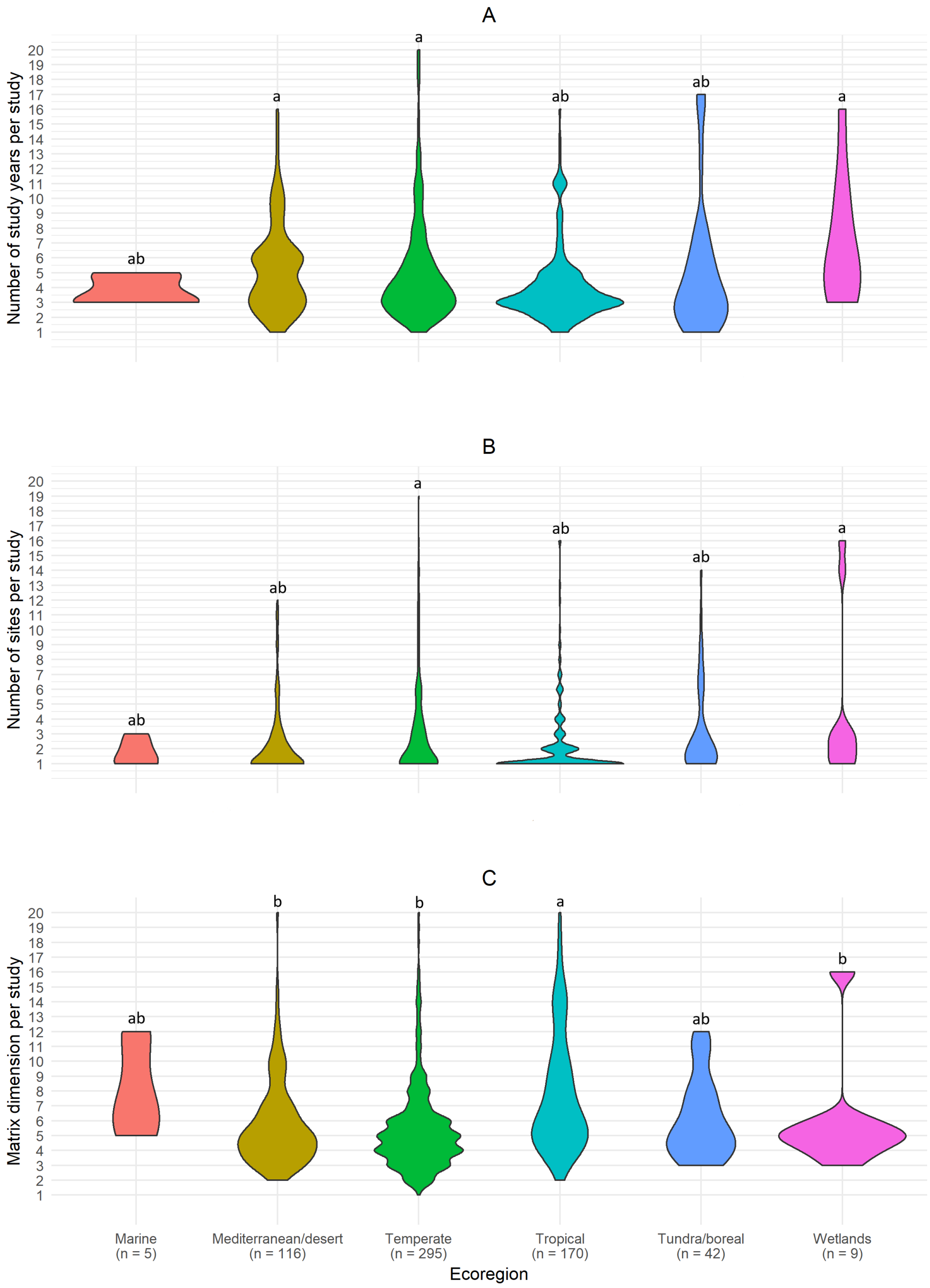


Figure S3. Variation in **(A)** number of study years, **(B)** number of populations and **(C)** matrix dimension across ecoregions per study. We used the number of studied years and populations and the matrix dimension for each studied species in each study. Small letters above the violins indicate groupings according to Tukey-test results. Ecoregions that share letters are not significantly different.

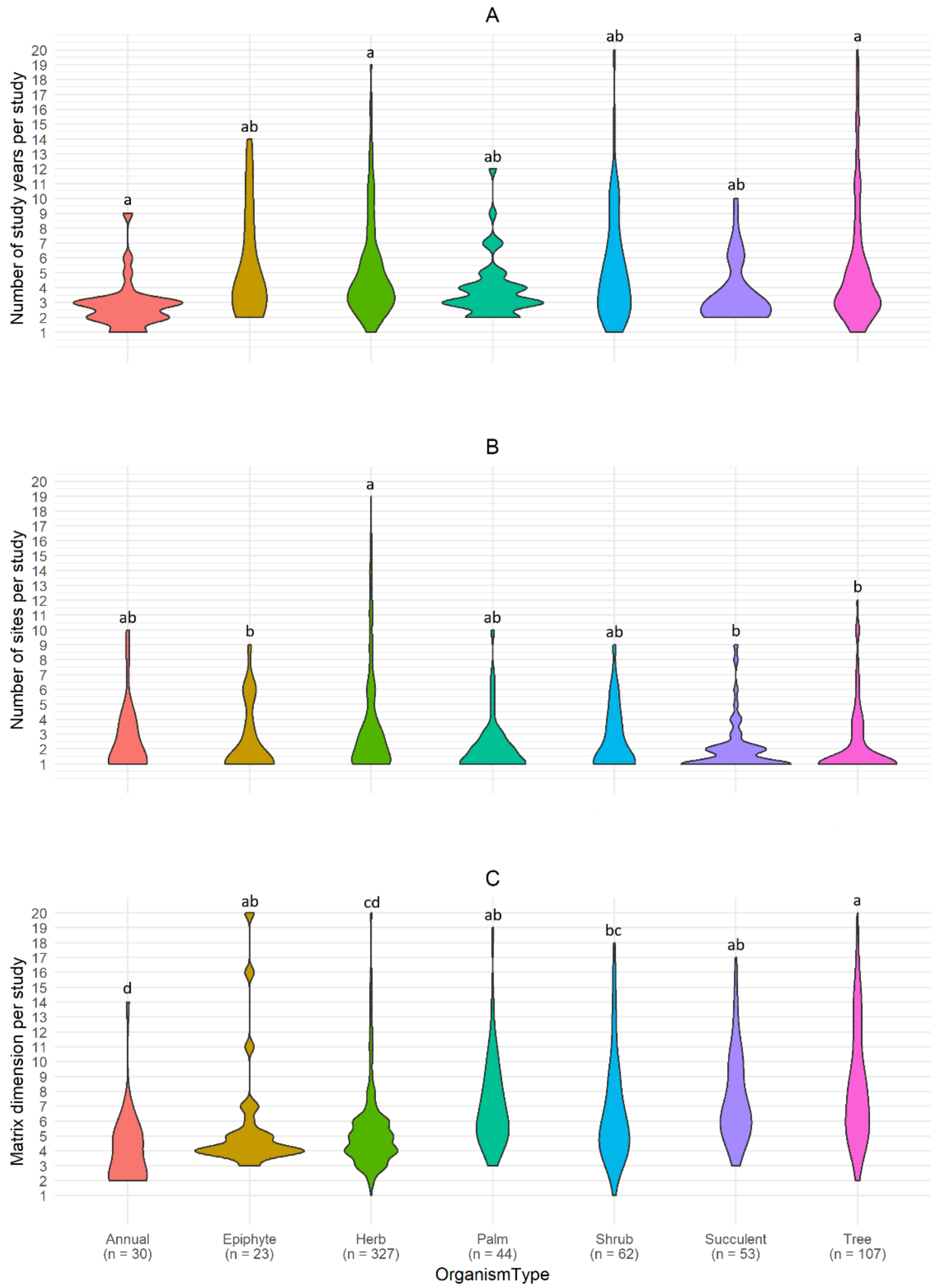


Figure S4. Variation in **(A)** number of study years, **(B)** number of populations and **(C)** matrix dimension across growth forms per study. We used the number of studied years and populations and the matrix dimension for each studied species in each study. Small letters above the violins indicate groupings according to Tukey-test results. Growth forms that share letters are not significantly different. Bryophyte, Liana and Fern were omitted due to small sample sizes.
