## Appendix S2: Analysis code for "Plant demographic knowledge is biased towards short-term studies of temperate-region herbaceous perennials"

### COMPADRE bias manuscript analysis code

#### Preamble

This document presents the code we used in our analysis for the manuscript, "*Plant demographic knowledge is biased towards short-term studies of temperate region herbaceous perennials*".

The analyses are performed in R v. 4.0.1, using RStudio. The document is split into six parts: (1) preparation of environment and data; (2) analysis of geographic bias; (3) analysis of temporal trends; (4) analysis of life history and taxonomic bias; (5) analysis of methodological bias; (6) material for the manuscript Appendix.

Note that we do not expand much on the rationale for the analyses, nor do we discuss the results in this document. For that we refer readers to the manuscript itself.

#### A) Preparation

At this stage we also make some corrections to various minor errors and inconsistencies in the data set (note that these errors have been corrected in later versions of COMPADRE). We also set up a plotting theme that we can use throughout.

##### Load libraries and set preferences

First we load the R packages required for the analyses.

```
library(tidyverse)
library(reshape)
library(conflicted)
library(directlabels)
library(agricolae)
library(Rcompadre)
library(popdemo)
library(mapproj)
library(patchwork)
```

Note that the `Rcompadre` package is available from GitHub at https://github.com/jonesor/Rcompadre.

Next we set conflict preferences.

```
conflict_prefer("filter", "dplyr")
conflict_prefer("rename", "dplyr")
conflict_prefer("map", "purrr")
```

#### Load and prepare COMPADRE data from file

We load the COMPADRE data file, which we have already downloaded from the COMPADRE database web site (htts://www.compadre-db.org).

New versions of the database are regularly made available at the COMPADRE database web site. Here we use version 5.0.0, which was released on June 6th, 2019, and is now available archived at https://github.com/jonesor/compadreDB/tree/master/OldDBVersions.

We establish a new data column, `Study`, which uniquely identifies individual scientific studies, and a column called `Row` as a convenient identifier.

We then make a copy of the compadre object called `db`, which we work from throughout.

```
database <- cdb_fetch("data/COMPADRE_v.5.0.0.RData")

## This is COMPADRE version 5.0.0 (release date Jun_06_2019)
## See user agreement at https://compadre-db.org/Help/UserAgreement
## See how to cite at https://compadre-db.org/Help/HowToCite

db <- cdb_metadata(database) %>%
  mutate(Study = as.integer(as.factor(paste(Authors, YearPublication,
DOI.ISBN)))) %>%
  mutate(Row = 1:n()) %>%
  as.data.frame()
```

#### Set up a plotting theme

We also establish a `ggplot` theme for our plots.

```
theme_set(theme_minimal() +
  theme(
    legend.title = element_blank(),
    legend.justification = c(1, 1), legend.position = c(1, 1),
    text = element_text(size = 12),
    axis.text.x = element_text(size = 12),
    axis.text.y = element_text(vjust = 0.5, size = 12),
    axis.title.y = element_text(margin = margin(t = 0, r = 10, b = 0, l =
0)),
    axis.title.x = element_text(margin = margin(t = 10, r = 0, b = 0, l = 0))
  ))
```

#### Correction of database errors

We identified a small number of data entry errors and inconsistencies in COMPADRE v.5.0.0. We have notified the database team and these errors will be corrected in later versions of the database. For the present analysis, however, we correct them here with the following code, which we divide into issues with *Biogeography*, *Taxonomy*, *OrganismType* and *Other errors*.

##### Biogeography

We correct the Ecoregion data for species where (i) the ecoregion is incorrect, or (ii) where a species has multiple ecoregion possibilities, but where we know that all populations are in the same ecoregion.

```
# change ecoregion variable names
db <- db %>%
  mutate(Ecoregion = ifelse(SpeciesAuthor == "Mircothlaspi_perfoliatum",
"TGS", Ecoregion)) %>%
  mutate(Ecoregion = ifelse(SpeciesAuthor == "Veronica_arvensis_2", "TGS",
Ecoregion)) %>%
  mutate(Ecoregion = ifelse(SpeciesAuthor == "Danthonia_sericea", "TCF",
Ecoregion)) %>%
  mutate(Ecoregion = ifelse(SpeciesAuthor == "Epilobium_latifolium", "TUN",
Ecoregion)) %>%
  mutate(Ecoregion = ifelse(SpeciesAuthor == "Eryngium_maritimum", "TBM",
Ecoregion)) %>%
  mutate(Ecoregion = ifelse(SpeciesAuthor == "Geum_reptans", "TCF",
Ecoregion)) %>%
  mutate(Ecoregion = ifelse(SpeciesAuthor == "Hudsonia_montana", "TBM",
Ecoregion)) %>%
  mutate(Ecoregion = ifelse(SpeciesAuthor == "Calamus_platyacanthus", "TMB",
Ecoregion)) %>%
  mutate(Ecoregion = ifelse(SpeciesAuthor == "Calamus_rhabdocladus", "TMB",
Ecoregion)) %>%
  mutate(Ecoregion = ifelse(SpeciesAuthor == "Chamaedorea_radicalis_2",
"TMB", Ecoregion)) %>%
  mutate(Ecoregion = ifelse(SpeciesAuthor == "Rubus_saxatilis", "TBM",
Ecoregion)) %>%
  mutate(Ecoregion = ifelse(SpeciesAuthor == "Primula_vulgaris_2", "TBM",
Ecoregion)) %>%
  mutate(Ecoregion = ifelse(SpeciesAuthor == "Rumex_rupestris", "TBM",
Ecoregion)) %>%
  mutate(Ecoregion = ifelse(SpeciesAuthor == "Saponaria_bellidifolia", "TCF",
Ecoregion)) %>%
  mutate(Ecoregion = ifelse(SpeciesAuthor == "Succisa_pratensis", "TBM",
Ecoregion)) %>%
  mutate(Ecoregion = ifelse(SpeciesAuthor == "Succisa_pratensis_2", "TBM",
Ecoregion)) %>%
  mutate(Ecoregion = ifelse(SpeciesAuthor == "Succisa_pratensis_3", "TBM",
Ecoregion)) %>%
  mutate(Ecoregion = ifelse(SpeciesAuthor == "Taraxacum_erythrospermum",
"TGS", Ecoregion)) %>%
  mutate(Ecoregion = ifelse(SpeciesAuthor == "Sabal_minor", "TGV",
Ecoregion)) %>%
  mutate(Ecoregion = ifelse(SpeciesAuthor == "Grias_peruviana", "TMB",
```

```
Ecoregion)) %>%
  mutate(Ecoregion = ifelse(SpeciesAuthor == "Eremophila_forrestii", "DES",
Ecoregion)) %>%
  mutate(Ecoregion = ifelse(SpeciesAuthor == "Eremophila_maitlandii", "DES",
Ecoregion)) %>%
  mutate(Ecoregion = ifelse(SpeciesAuthor == "Guettarda_viburnoides", "TGV",
Ecoregion)) %>%
  mutate(Ecoregion = ifelse(SpeciesAuthor == "Khaya_senegalensis_2", "TGV",
Ecoregion)) %>%
  mutate(Ecoregion = ifelse(SpeciesAuthor == "Prunus_africana", "TMB",
Ecoregion)) %>%
  mutate(Ecoregion = ifelse(SpeciesAuthor == "Daemonorops_poilanei", "TMB",
Ecoregion)) %>%
  mutate(Ecoregion = ifelse(SpeciesAuthor == "Dioon_caputoi", "TMB",
Ecoregion)) %>%
  mutate(Ecoregion = ifelse(SpeciesAuthor == "Microthlaspi_perfoliatum",
"TGS", Ecoregion)) %>%
  mutate(Ecoregion = ifelse(SpeciesAuthor == "Cirsium_perplexans", "TCF",
Ecoregion)) %>%
  mutate(Ecoregion = ifelse(SpeciesAuthor == "Cirsium_perplexans_2", "TCF",
Ecoregion)) %>%
  mutate(Ecoregion = ifelse(SpeciesAuthor == "Agave_augustifolia", "TDB",
Ecoregion)) %>%
  mutate(Ecoregion = ifelse(SpeciesAuthor == "Furcraea_parmentieri", "TSC",
Ecoregion)) %>%
  mutate(Ecoregion = ifelse(SpeciesAuthor == "Froelichia_floridana", "TBM",
Ecoregion)) %>%
  mutate(Ecoregion = ifelse(SpeciesAuthor == "Melocactus_ernestii", "DES",
Ecoregion)) %>%
  mutate(Ecoregion = ifelse(SpeciesAuthor == "Alyxia_stellata", "TMB",
Ecoregion)) %>%
  mutate(Ecoregion = ifelse(SpeciesAuthor == "Coccothrinax_readii", "TMB",
Ecoregion)) %>%
  mutate(Ecoregion = ifelse(SpeciesAuthor == "Choerospondias_axillaris",
"TDB", Ecoregion)) %>%
  # species with multiple populations in different ecoregions
  mutate(Ecoregion = ifelse(SpeciesAuthor == "Dactylorhiza_lapponica" &
    MatrixPopulation == "Solendet", "TUN", Ecoregion)) %>%
  mutate(Ecoregion = ifelse(SpeciesAuthor == "Dactylorhiza_lapponica" &
    MatrixPopulation == "Tagdalen", "BOR", Ecoregion)) %>%
  mutate(Ecoregion = ifelse(SpeciesAuthor == "Scaphium_macropodum" &
    MatrixPopulation == "Cattien", "TDB", Ecoregion)) %>%
  mutate(Ecoregion = ifelse(SpeciesAuthor == "Scaphium_macropodum" &
    MatrixPopulation == "Dakuy", "TDB", Ecoregion)) %>%
  mutate(Ecoregion = ifelse(SpeciesAuthor == "Scaphium_macropodum" &
    MatrixPopulation == "Bachma National Park", "TMB", Ecoregion)) %>%
  mutate(Ecoregion = ifelse(SpeciesAuthor == "Plantago_coronopus_2" &
    MatrixPopulation == "T", "MED", Ecoregion)) %>%
  mutate(Ecoregion = ifelse(SpeciesAuthor == "Plantago_coronopus_2" &
    MatrixPopulation == "CA", "MED", Ecoregion)) %>%
```

```
  mutate(Ecoregion = ifelse(SpeciesAuthor == "Plantago_coronopus_2" &
    MatrixPopulation == "C", "TBM", Ecoregion)) %>%
  mutate(Ecoregion = ifelse(SpeciesAuthor == "Plantago_coronopus_2" &
    MatrixPopulation == "TB", "TBM", Ecoregion)) %>%
  mutate(Ecoregion = ifelse(SpeciesAuthor == "Plantago_coronopus_2" &
    MatrixPopulation == "DH", "TBM", Ecoregion)) %>%
  mutate(Ecoregion = ifelse(SpeciesAuthor == "Plantago_coronopus_2" &
    MatrixPopulation == "DS", "TBM", Ecoregion)) %>%
  mutate(Ecoregion = ifelse(SpeciesAuthor == "Plantago_coronopus_2" &
    MatrixPopulation == "SG", "TBM", Ecoregion)) %>%
  mutate(Ecoregion = ifelse(SpeciesAuthor == "Plantago_coronopus_2" &
    MatrixPopulation == "ST", "TBM", Ecoregion)) %>%
  mutate(Ecoregion = ifelse(SpeciesAuthor == "Plantago_coronopus_2" &
    MatrixPopulation == "EA", "TBM", Ecoregion)) %>%
  mutate(Ecoregion = ifelse(SpeciesAuthor == "Plantago_coronopus_2" &
    MatrixPopulation == "ES", "TBM", Ecoregion)) %>%
  mutate(Ecoregion = ifelse(SpeciesAuthor == "Plantago_coronopus_2" &
    MatrixPopulation == "F", "TBM", Ecoregion)) %>%
  mutate(Ecoregion = ifelse(SpeciesAuthor == "Ligularia_sibirica" &
    MatrixPopulation == "CZ N/1", "TBM", Ecoregion)) %>%
  mutate(Ecoregion = ifelse(SpeciesAuthor == "Ligularia_sibirica" &
    MatrixPopulation == "CZ N/2", "TBM", Ecoregion)) %>%
  mutate(Ecoregion = ifelse(SpeciesAuthor == "Ligularia_sibirica" &
    MatrixPopulation == "CZ N/3", "TBM", Ecoregion)) %>%
  mutate(Ecoregion = ifelse(SpeciesAuthor == "Ligularia_sibirica" &
    MatrixPopulation == "CZ N/4", "TBM", Ecoregion)) %>%
  mutate(Ecoregion = ifelse(SpeciesAuthor == "Ligularia_sibirica" &
    MatrixPopulation == "CZ N/5", "TBM", Ecoregion)) %>%
  mutate(Ecoregion = ifelse(SpeciesAuthor == "Ligularia_sibirica" &
    MatrixPopulation == "CZ C/1", "TBM", Ecoregion)) %>%
  mutate(Ecoregion = ifelse(SpeciesAuthor == "Ligularia_sibirica" &
    MatrixPopulation == "CZ C/2", "TBM", Ecoregion)) %>%
  mutate(Ecoregion = ifelse(SpeciesAuthor == "Ligularia_sibirica" &
    MatrixPopulation == "CZ C/3", "TBM", Ecoregion)) %>%
  mutate(Ecoregion = ifelse(SpeciesAuthor == "Ligularia_sibirica" &
    MatrixPopulation == "CZ S/1", "TBM", Ecoregion)) %>%
  mutate(Ecoregion = ifelse(SpeciesAuthor == "Ligularia_sibirica" &
    MatrixPopulation == "SK SR", "TCF", Ecoregion)) %>%
  mutate(Ecoregion = ifelse(SpeciesAuthor == "Ligularia_sibirica" &
    MatrixPopulation == "SK P/1", "TBM", Ecoregion)) %>%
  # add labels (ecoregionLabel) for terrestrial categories
  mutate(ecoregionLabel = NA) %>%
  mutate(ecoregionLabel = ifelse(Ecoregion == "TMB", "Tropical",
ecoregionLabel)) %>%
  mutate(ecoregionLabel = ifelse(Ecoregion == "TDB", "Tropical",
ecoregionLabel)) %>%
  mutate(ecoregionLabel = ifelse(Ecoregion == "TSC", "Tropical",
ecoregionLabel)) %>%
  mutate(ecoregionLabel = ifelse(Ecoregion == "TGV", "Tropical",
ecoregionLabel)) %>%
```

```
  mutate(ecoregionLabel = ifelse(Ecoregion == "TBM", "Temperate",
ecoregionLabel)) %>%
  mutate(ecoregionLabel = ifelse(Ecoregion == "TCF", "Temperate",
ecoregionLabel)) %>%
  mutate(ecoregionLabel = ifelse(Ecoregion == "TGS", "Temperate",
ecoregionLabel)) %>%
  mutate(ecoregionLabel = ifelse(Ecoregion == "MED", "Mediterranean/desert",
ecoregionLabel)) %>%
  mutate(ecoregionLabel = ifelse(Ecoregion == "DES", "Mediterranean/desert",
ecoregionLabel)) %>%
  mutate(ecoregionLabel = ifelse(Ecoregion == "BOR", "Tundra/boreal",
ecoregionLabel)) %>%
  mutate(ecoregionLabel = ifelse(Ecoregion == "TUN", "Tundra/boreal",
ecoregionLabel)) %>%
  mutate(ecoregionLabel = ifelse(Ecoregion == "MON", "Tundra/boreal",
ecoregionLabel)) %>%
  mutate(ecoregionLabel = ifelse(Ecoregion == "MAN", "Wetlands",
ecoregionLabel)) %>%
  mutate(ecoregionLabel = ifelse(Ecoregion == "FGS", "Wetlands",
ecoregionLabel)) %>%
  # add label for marine categories
  mutate(ecoregionLabel = ifelse(Ecoregion == "POE", "Marine",
ecoregionLabel)) %>%
  mutate(ecoregionLabel = ifelse(Ecoregion == "TSS", "Marine",
ecoregionLabel)) %>%
  mutate(ecoregionLabel = ifelse(Ecoregion == "TEU", "Marine",
ecoregionLabel)) %>%
  mutate(ecoregionLabel = ifelse(Ecoregion == "TRU", "Marine",
ecoregionLabel)) %>%
  mutate(ecoregionLabel = ifelse(Ecoregion == "TRC", "Marine",
ecoregionLabel)) %>%
  # change label for species assigned to two ecoregions that fall into the
same categories in our system
  mutate(ecoregionLabel = ifelse(Ecoregion == "TMB; TDB", "Tropical",
ecoregionLabel)) %>%
  mutate(ecoregionLabel = ifelse(Ecoregion == "BOR; TUN", "Tundra/boreal",
ecoregionLabel)) %>%
  # correct continent where wrong or missing
  mutate(Continent = ifelse(SpeciesAuthor == "Alyxia_stellata", "N America",
Continent)) %>%
  unique()
```

##### Taxonomy

Next, we correct some inconsistencies with some recorded taxonomic information.

```
db <- db %>%
  mutate(AngioGymno = as.character(AngioGymno)) %>%
  mutate(DicotMonoc = as.character(DicotMonoc)) %>%
  # Change AngioGymno to "other" when Algae, Bryophyte or Fern
```

```
  mutate(AngioGymno = ifelse(OrganismType == "Algae", "other", AngioGymno))
%>%
  mutate(AngioGymno = ifelse(OrganismType == "Bryophyte", "other",
AngioGymno)) %>%
  mutate(AngioGymno = ifelse(OrganismType == "Fern", "other", AngioGymno))
%>%
  # correct typos in database for AngioGymno
  mutate(AngioGymno = ifelse(AngioGymno == "Gymno", "Gymnosperm",
AngioGymno)) %>%
  # add information for species with missing information
  mutate(AngioGymno = ifelse(SpeciesAuthor == "Alliaria_petiolata",
"Angiosperm", AngioGymno)) %>%
  mutate(DicotMonoc = ifelse(SpeciesAuthor == "Alliaria_petiolata",
"Eudicot", DicotMonoc)) %>%
  mutate(OrganismType = ifelse(SpeciesAuthor == "Alliaria_petiolata",
"Herbaceous perennial", OrganismType)) %>%
  mutate(AngioGymno = ifelse(SpeciesAuthor == "Anthyllis_vulneraria",
"Angiosperm", AngioGymno)) %>%
  mutate(DicotMonoc = ifelse(SpeciesAuthor == "Anthyllis_vulneraria",
"Eudicot", DicotMonoc)) %>%
  mutate(OrganismType = ifelse(SpeciesAuthor == "Anthyllis_vulneraria",
"Herbaceous perennial", OrganismType)) %>%
  mutate(AngioGymno = ifelse(SpeciesAuthor == "Cirsium_vulgare",
"Angiosperm", AngioGymno)) %>%
  mutate(DicotMonoc = ifelse(SpeciesAuthor == "Cirsium_vulgare", "Eudicot",
DicotMonoc)) %>%
  mutate(OrganismType = ifelse(SpeciesAuthor == "Cirsium_vulgare",
"Herbaceous perennial", OrganismType)) %>%
  mutate(AngioGymno = ifelse(SpeciesAuthor == "Scorzonera_humilis",
"Angiosperm", AngioGymno)) %>%
  mutate(DicotMonoc = ifelse(SpeciesAuthor == "Scorzonera_humilis",
"Eudicot", DicotMonoc)) %>%
  mutate(OrganismType = ifelse(SpeciesAuthor == "Scorzonera_humilis",
"Herbaceous perennial", OrganismType)) %>%
  mutate(AngioGymno = ifelse(SpeciesAuthor == "Ulex_minor", "Angiosperm",
AngioGymno)) %>%
  mutate(DicotMonoc = ifelse(SpeciesAuthor == "Ulex_minor", "Eudicot",
DicotMonoc)) %>%
  mutate(OrganismType = ifelse(SpeciesAuthor == "Ulex_minor", "Herbaceous
perennial", OrganismType)) %>%
  mutate(AngioGymno = ifelse(SpeciesAuthor == "Eritrichium_caucasicum",
"Angiosperm", AngioGymno)) %>%
  mutate(DicotMonoc = ifelse(SpeciesAuthor == "Eritrichium_caucasicum",
"Eudicot", DicotMonoc)) %>%
  mutate(OrganismType = ifelse(SpeciesAuthor == "Eritrichium_caucasicum",
"Herbaceous perennial", OrganismType)) %>%
  mutate(AngioGymno = ifelse(SpeciesAuthor == "Geum_reptans", "Angiosperm",
AngioGymno)) %>%
  mutate(DicotMonoc = ifelse(SpeciesAuthor == "Geum_reptans", "Eudicot",
DicotMonoc)) %>%
```

```
  mutate(OrganismType = ifelse(SpeciesAuthor == "Geum_reptans", "Herbaceous
perennial", OrganismType)) %>%
  mutate(AngioGymno = ifelse(SpeciesAuthor == "Silene_spaldingii_2",
"Angiosperm", AngioGymno)) %>%
  mutate(DicotMonoc = ifelse(SpeciesAuthor == "Silene_spaldingii_2",
"Eudicot", DicotMonoc)) %>%
  mutate(OrganismType = ifelse(SpeciesAuthor == "Silene_spaldingii_2",
"Herbaceous perennial", OrganismType)) %>%
  mutate(AngioGymno = ifelse(SpeciesAuthor == "Hybrid: Raphanus_raphanistrum
x R. sativus", "Angiosperm", AngioGymno)) %>%
  mutate(DicotMonoc = ifelse(SpeciesAuthor == "Hybrid: Raphanus_raphanistrum
x R. sativus", "Eudicot", DicotMonoc)) %>%
  mutate(OrganismType = ifelse(SpeciesAuthor == "Hybrid:
Raphanus_raphanistrum x R. sativus", "Herbaceous perennial", OrganismType))
```

##### OrganismType

Next, we add `OrganismType` information for species where it is missing in the database. We also correct some database inconsistencies/typos.

```
db <- db %>%
  mutate(OrganismType = ifelse(SpeciesAuthor == "Alliaria_petiolata",
"Herbaceous perennial", OrganismType)) %>%
  mutate(OrganismType = ifelse(SpeciesAuthor == "Anthyllis_vulneraria",
"Herbaceous perennial", OrganismType)) %>%
  mutate(OrganismType = ifelse(SpeciesAuthor == "Cirsium_vulgare",
"Herbaceous perennial", OrganismType)) %>%
  mutate(OrganismType = ifelse(SpeciesAuthor == "Scorzonera_humilis",
"Herbaceous perennial", OrganismType)) %>%
  mutate(OrganismType = ifelse(SpeciesAuthor == "Eritrichium_caucasicum",
"Herbaceous perennial", OrganismType)) %>%
  mutate(OrganismType = ifelse(SpeciesAuthor == "Geum_reptans", "Herbaceous
perennial", OrganismType)) %>%
  mutate(OrganismType = ifelse(SpeciesAuthor == "Silene_spaldingii_2",
"Herbaceous perennial", OrganismType)) %>%
  mutate(OrganismType = ifelse(SpeciesAuthor == "Ulex_minor", "Herbaceous
perennial", OrganismType)) %>%
  mutate(OrganismType = ifelse(SpeciesAuthor == "Hybrid:
Raphanus_raphanistrum x R. sativus",
    "Herbaceous perennial", OrganismType
  )) %>%
  # correct typo
  mutate(OrganismType = ifelse(OrganismType == "Herbaceous perennials",
"Herbaceous perennial", OrganismType)) %>%
  # change Herbaceous perennial to Herb
  mutate(OrganismType = ifelse(OrganismType == "Herbaceous perennial",
"Herb", OrganismType))
```

##### Other errors

We noticed that the data for `Lepanthes_rubripetala_2` had the incorrect number of populations (`NumberPopulations`) listed. We correct that to `6` here after consulting the original publication.

```
db <- db %>%
  # correct number of populations
  mutate(NumberPopulations = ifelse(SpeciesAuthor ==
"Lepanthes_rubripetala_2", 6, NumberPopulations))
```

### B) WHEN HAVE THE STUDIES BEEN PUBLISHED?

In this section, we analyse patterns of WHEN studies have been published. This includes (i) a comparison of the total number of MPM papers published and MPM papers released in the COMPADRE database, through time. (ii) an estimation of the proportion of plant ecology papers that use MPMs through time.

#### Digitisation progress in COMPADRE

##### Published papers

The COMPADRE Plant Matrix Database digitisation team maintains a list of all known publications containing MPMs and works steadily to digitise this growing list. The COMPADRE team provided us with a list of literature (`Full_List accepted literature.csv`) which we use to produce a picture of MPM publication and digitisation efforts.

```
MPMLiterature <- read.csv("data/Full_List accepted literature.csv") %>%
  mutate(YearPublication = as.integer(as.character(YearPublication))) %>%
  mutate(digitized = as.factor(paste(Authors, YearPublication))) %>%
  distinct(digitized, .keep_all = TRUE)
```

##### Cumulative publication and digitisation through time

To look at publication and digitization of MPM papers through time, we create a data frame including information on (known) published papers and papers included in the database for the period 1966-2019. We do this by comparing data available in COMPADRE to the list of all papers containing MPMs.

```
digitisationData <- left_join(MPMLiterature,
  db %>%
    mutate(digitized = as.factor(paste(Authors, YearPublication))) %>%
    select(Authors, SpeciesAuthor, YearPublication, Journal, OrganismType,
digitized) %>%
    unique() %>%
    distinct(digitized, .keep_all = TRUE) %>%
    mutate(included = "YES") %>%
    select(SpeciesAuthor, included),
```

```
  by = "SpeciesAuthor"
) %>%
  mutate(included = ifelse(is.na(included), "NO", included))
```

We now create a data frame including information on (known) published papers and papers included in the database for the period 1966-2019.

```
# change NAs to zeros and calculate cumulative sums
data_years <- left_join(left_join(data.frame(YearPublication = 1966:2019),
  digitisationData %>%
    mutate(YearPublication = as.integer(YearPublication)) %>%
    group_by(YearPublication) %>%
    summarise(published = length(unique(digitized))) %>%
    filter(YearPublication != "NDY"),
  by = "YearPublication"
), digitisationData %>%
  filter(included == "YES") %>%
  mutate(YearPublication = as.numeric(YearPublication)) %>%
  group_by(YearPublication) %>%
  summarise(COMPADRE_digi = length(unique(digitized))) %>%
  filter(!is.na(YearPublication)), by = "YearPublication") %>%
  mutate(published = ifelse(is.na(published), 0, published)) %>%
  mutate(COMPADRE_digi = ifelse(is.na(COMPADRE_digi), 0, COMPADRE_digi)) %>%
  mutate(CumSum_p = cumsum(published)) %>%
  mutate(CumSum_d = cumsum(COMPADRE_digi))
```

##### Plot digitisation progress

We then plot the data frame to create Fig. 1A.

```
Fig1A <- ggplot() +
  # plot as lines
  geom_line(data = data_years, aes(x = YearPublication, y = (CumSum_d - 1),
color = "COMPADRE"), size = 1) +
  geom_line(data = data_years, aes(x = YearPublication, y = CumSum_p, color =
"published"), size = 1) +

  # adjust x and y axis
  scale_x_continuous(
    breaks = seq(0, 2020, 10),
    limits = c(1970, 2020)
  ) +
  xlab("Year of publication") +
  ylab("Number of studies") +
  scale_color_manual(
    values = c("#25b7bc", "black"),
    breaks = c("published", "COMPADRE"),
    labels = c(
      " MPM papers published",
      " MPM papers included in COMPADRE"
    )
```

```
  ) +

  # adjust legend
  theme(
    legend.justification = c(0.1, 1),
    legend.position = c(0.1, 0.91),
    panel.grid.minor.x = element_blank(),
    panel.grid.major.x = element_blank()
  ) +
  NULL

Fig1A
```

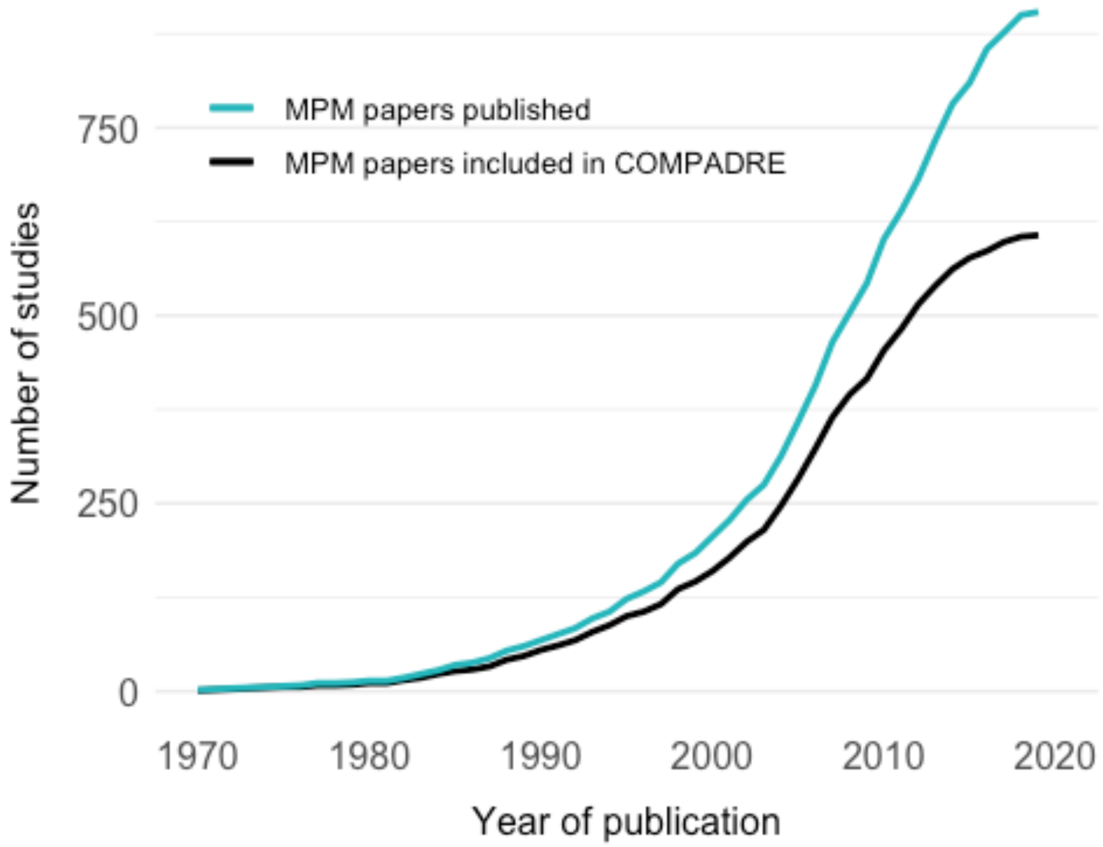


#### The relative importance of MPMs in the literature

Is demographic research using MPMs becoming increasingly important within ecology? We investigate this by examining how the proportion of articles that use MPMs changes through time, from a sample of literature published in Journal of Ecology.

##### Plot proportion through time

We have downloaded Journal of Ecology bibliographic information to the folder `WoS_search` and we scan the text (abstracts and keywords) to create a data frame summarising the use of MPMs through time.

```
# Load and combine data, to build the dataset
abstractData <- list.files(path = "data/WoS_search/", full.names = TRUE) %>%
  purrr::map(read_delim, delim = "\t", quote = "") %>%
  # read in all the files individually
  reduce(rbind) %>%
  # reduce into one dataframe
  # Find mentions of MPMs in abstract and keywords/topic areas.
  mutate(matrixInAbstract = str_detect(AB, "projection model|matrix
model|Matrix model|MPM")) %>%
  mutate(matrixInKeywords1 = str_detect(DE, "projection model|matrix
model|Matrix model|MPM")) %>%
  mutate(matrixInKeywords2 = str_detect(ID, "projection model|matrix
model|Matrix model|MPM")) %>%
  mutate(matrixModel = matrixInAbstract + matrixInKeywords1 +
matrixInKeywords2)

# prepare dataframe
papersPerYear <- abstractData %>%
  filter(!is.na(AB)) %>%
  group_by(PY) %>%
  summarise(N = length(PY))

MPMsInEcology <- abstractData %>%
  filter(matrixModel > 0) %>%
  group_by(PY) %>%
  summarise(NMatrix = length(PY)) %>%
  left_join(papersPerYear) %>%
  mutate(Proportion = NMatrix / N) %>%
  rename(N_withMPM = NMatrix, N_total = N) %>%
  rename(PublicationYear = "PY") %>%
  mutate(Percentage = Proportion * 100)
```

We now model this relationship and see that there is no significant relationship between year and percentage of *J. Ecol.* papers containing MPMs.

```
mod_publication_trend <- lm(Percentage ~ PublicationYear, data =
MPMsInEcology)
summary(mod_publication_trend)

##
## Call:
## lm(formula = Percentage ~ PublicationYear, data = MPMsInEcology)
##
## Residuals:
```

```
##      Min       1Q  Median      3Q     Max 
## -2.1680 -0.8732 -0.5439  0.6875  4.5328 
## 
## Coefficients:
##                  Estimate Std. Error t value Pr(>|t|)
## (Intercept)     -65.65877   87.84357  -0.747    0.463
## PublicationYear   0.03394    0.04378   0.775    0.447
## 
## Residual standard error: 1.629 on 21 degrees of freedom
## Multiple R-squared:  0.02783,    Adjusted R-squared:  -0.01847 
## F-statistic: 0.6011 on 1 and 21 DF,  p-value: 0.4468
```

##### plot relative importance of MPMs in the sample

We remove data before 1993, because no abstracts are available for that period, then plot the data create Fig. 1B:

```
Fig1B <- ggplot(MPMsInEcology %>% filter(PublicationYear > 1992), aes(x = 
PublicationYear, y = Percentage)) +
  geom_line(linetype = "dashed", color = "#25b7bc", size = 0.5) +
  geom_point(color = "#25b7bc", size = 2) +

  # adjust x and y axis
  scale_x_continuous(
    breaks = seq(0, 2020, 10),
    limits = c(1970, 2020)
  ) +
  xlab("Year of publication") +
  ylab("Percentage of studies") +

  # adjust legend
  theme(
    legend.justification = c(0.1, 1),
    legend.position = c(0.1, 0.91),
    panel.grid.minor.x = element_blank(),
    panel.grid.major.x = element_blank()
  ) +
  NULL

Fig1B
```

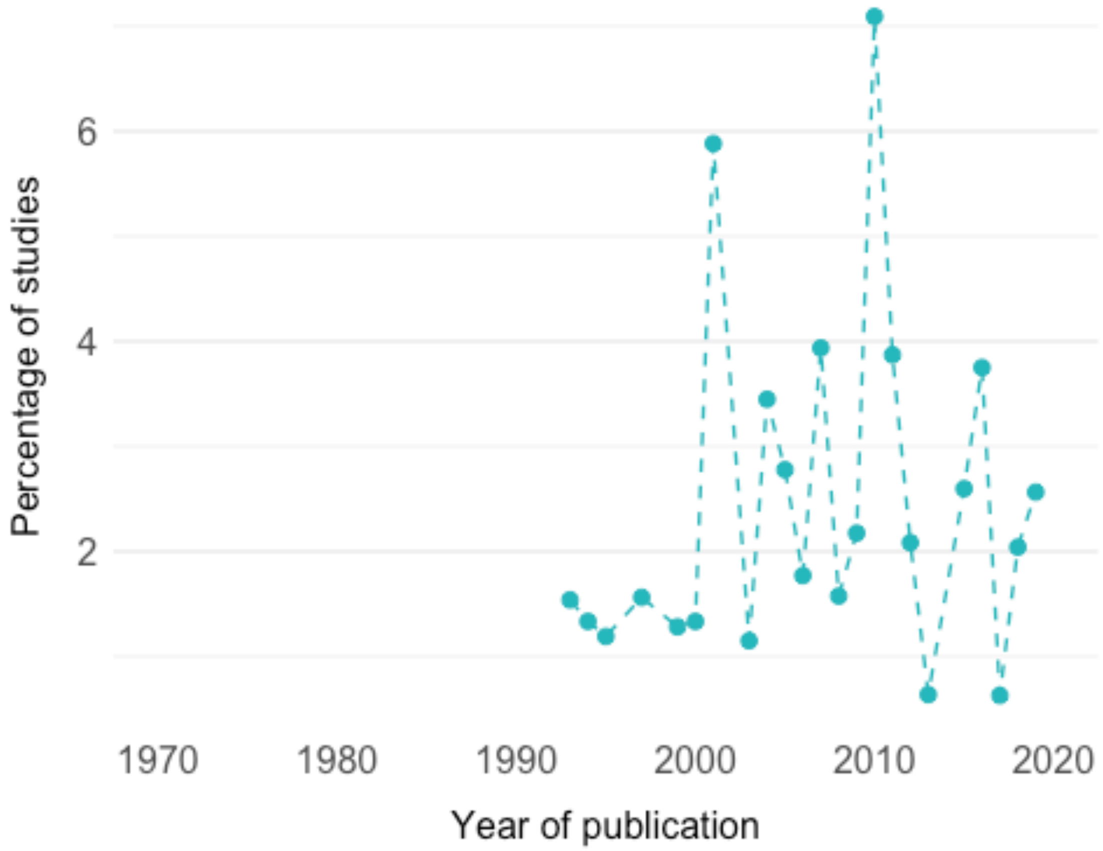


We combine Fig. 1A and Fig. 1B:

```
(Fig1A + ggtitle("A")) / (Fig1B + ggtitle("B"))
```

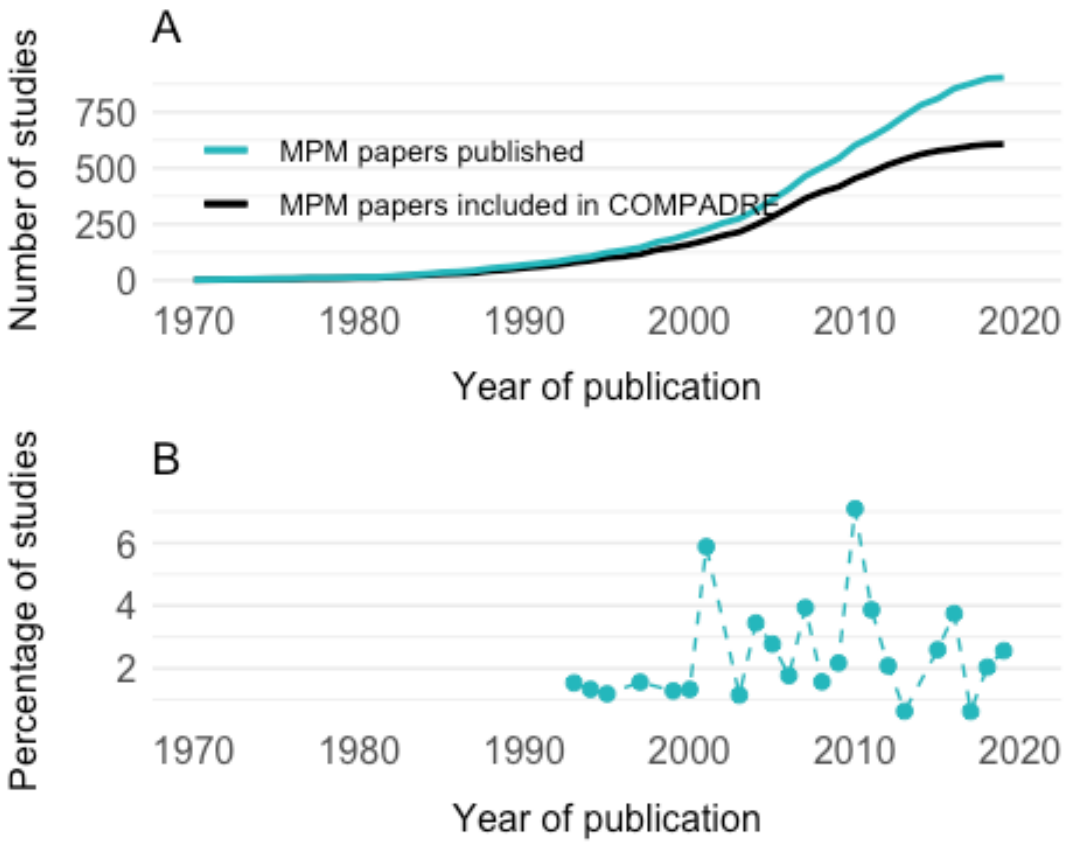


#### C) WHERE IS THE RESEARCH DONE?

This section contains the analyses on WHERE studies have been carried out, focussing on geographic biases. This includes:

- The distribution of species among ecoregions in COMPADRE compared to the natural distribution
- The distribution of plant demography study density across continents
- The relationship between country per-capita GDP and number of plant demography studies

##### Distribution among ecoregions

We here examine how well the data in COMPADRE represent true plant distributions across ecoregions.

Because epiphytes include both vascular and non-vascular plants, and our analysis is solely focussed on vascular plants, we check which epiphytes in COMPADRE are vascular plants.

```r
table(db %>%
  filter(OrganismType == "Epiphyte") %>%
  pull(Class))
```

```
##
## Liliopsida
##        327
```

We see that all of COMPADRE's epiphytes are in Liliopsida and are thus all vascular.

Next, we remove algae, bryophytes and non-vascular epiphytes from the subset, and summarise the resulting data by ecoregion:

```r
compadreEcoregionSummary <- db %>%
  select(Study, SpeciesAuthor, Ecoregion, ecoregionLabel, Continent, Country,
OrganismType) %>%
  filter(!OrganismType %in% c("Algae", "Bryophyte")) %>%
  select(SpeciesAuthor, ecoregionLabel) %>%
  filter(ecoregionLabel != is.na(ecoregionLabel)) %>%
  unique() %>%
  group_by(ecoregionLabel) %>%
  summarise(CompadreSpeciesCount = n()) %>%
  mutate(ecoregionLabel = as.factor(ecoregionLabel))

compadreEcoregionSummary
```

```
## # A tibble: 5 x 2
##   ecoregionLabel       CompadreSpeciesCount
## * <fct>                               <int>
## 1 Mediterranean/desert                  155
## 2 Temperate                             416
## 3 Tropical                              230
## 4 Tundra/boreal                          48
## 5 Wetlands                               10
```

We use the dataset from Kier et al. (2005) [1] and summarise it to provide an estimate of plant species distribution among global ecoregions.

```r
# load data
kierEcoregionSummary <- read.csv("data/kier_ecoegions.csv") %>%
  mutate(ecoregionLabel = Biome) %>%
  mutate(ecoregionLabel = ifelse(ecoregionLabel %in% c("BOR", "TUN", "MON"),
"Tundra/boreal", ecoregionLabel)) %>%
  mutate(ecoregionLabel = ifelse(ecoregionLabel %in% c("TMB", "TDB", "TSC",
"TGV"), "Tropical", ecoregionLabel)) %>%
```

[1] Kier, G., Mutke, J., Dinerstein, E., Ricketts, T. H., Küper, W., Kreft, H., & Barthlott, W. (2005). Global patterns of plant diversity and floristic knowledge. Journal of Biogeography, 32(7), 1107–1116.

```
  mutate(ecoregionLabel = ifelse(ecoregionLabel %in% c("TBM", "TCF", "TGS"),
"Temperate", ecoregionLabel)) %>%
  mutate(ecoregionLabel = ifelse(ecoregionLabel %in% c("MED", "DES"),
"Mediterranean/desert", ecoregionLabel)) %>%
  mutate(ecoregionLabel = ifelse(ecoregionLabel %in% c("MAN", "FGS"),
"Wetlands", ecoregionLabel)) %>%
  group_by(ecoregionLabel) %>%
  summarise(KierSpeciesCount = sum(PlantNumber))
```

We then process the COMPADRE data and combine it with the Kier data so that they can be compared. Specifically, we make a data frame of species counts in each ecoregion, and also express those counts as percentages.

```
ecoregionData <- left_join(kierEcoregionSummary, compadreEcoregionSummary)
%>%
  left_join(data.frame(
    ecoregionAbb = c("Trop", "Temp", "Med/Des", "Tund/Bor", "Wetl"),
    ecoregionLabel = c(
      "Tropical", "Temperate", "Mediterranean/desert",
      "Tundra/boreal", "Wetlands"
    )
  )) %>%
  mutate(KierPercentage = KierSpeciesCount / sum(KierSpeciesCount)) %>%
  mutate(CompadrePercentage = CompadreSpeciesCount /
sum(CompadreSpeciesCount))

## Joining, by = "ecoregionLabel"
## Joining, by = "ecoregionLabel"

ecoregionData

## # A tibble: 5 x 6
##   ecoregionLabel   KierSpeciesCount CompadreSpecies… ecoregionAbb
KierPercentage
##   <chr>                       <int>            <int> <chr>
<dbl>
## 1 Mediterranean/d…             3372              155 Med/Des
0.165
## 2 Temperate                    4851              416 Temp
0.238
## 3 Tropical                     8557              230 Trop
0.419
## 4 Tundra/boreal                2657               48 Tund/Bor
0.130
## 5 Wetlands                      972               10 Wetl
0.0476
## # … with 1 more variable: CompadrePercentage <dbl>
```

We rearrange this data for convenience when plotting.

```r
ecoregionKierCompadre <- ecoregionData %>%
  select(ecoregionAbb, ends_with("Percentage")) %>%
  pivot_longer(cols = ends_with("Percentage"), names_to = "DataSource") %>%
  mutate(DataSource = gsub("Percentage", "", DataSource))
```

We test the statistical significance of these differences with `chisq.test` i.e., we ask if the distribution of species across ecoregions in COMPADRE is significantly different than the global distributions summarised in the Kier data. We find that the distribution in COMPADRE is indeed significantly different than the true distribution.

```r
chisq.test(x = ecoregionData$CompadreSpeciesCount, p =
ecoregionData$KierPercentage) # all

##
##  Chi-squared test for given probabilities
##
## data:  ecoregionData$CompadreSpeciesCount
## X-squared = 327.79, df = 4, p-value < 2.2e-16
```

We test the pairwise differences using `prop.test`. The total number of species in Kier is 20409 and the total number is COMPADRE is 859.

```r
#Tropical
prop.test(x = c(8557,230), n = c(20409,859))

##
##  2-sample test for equality of proportions with continuity correction
##
## data:  c(8557, 230) out of c(20409, 859)
## X-squared = 77.432, df = 1, p-value < 2.2e-16
## alternative hypothesis: two.sided
## 95 percent confidence interval:
##  0.1205414 0.1825038
## sample estimates:
##    prop 1    prop 2
## 0.4192758 0.2677532

#Temperate
prop.test(x = c(4851,416), n = c(20409,859))

##
##  2-sample test for equality of proportions with continuity correction
##
## data:  c(4851, 416) out of c(20409, 859)
## X-squared = 267.71, df = 1, p-value < 2.2e-16
## alternative hypothesis: two.sided
## 95 percent confidence interval:
##  -0.2811278 -0.2120618
## sample estimates:
##    prop 1    prop 2
## 0.2376893 0.4842841
```

```
#Med/Des
prop.test(x = c(3372, 155), n = c(20409,859))

##
##  2-sample test for equality of proportions with continuity correction
##
## data:  c(3372, 155) out of c(20409, 859)
## X-squared = 1.2727, df = 1, p-value = 0.2593
## alternative hypothesis: two.sided
## 95 percent confidence interval:
##  -0.04204403  0.01160173
## sample estimates:
##    prop 1    prop 2
## 0.1652212 0.1804424

#Tundra/boreal
prop.test(x = c(2657,48), n = c(20409,859))

##
##  2-sample test for equality of proportions with continuity correction
##
## data:  c(2657, 48) out of c(20409, 859)
## X-squared = 40.335, df = 1, p-value = 2.139e-10
## alternative hypothesis: two.sided
## 95 percent confidence interval:
##  0.05766340 0.09095407
## sample estimates:
##     prop 1     prop 2
## 0.13018766 0.05587893

#Wetlands
prop.test(x = c(972,10), n = c(20409,859))

##
##  2-sample test for equality of proportions with continuity correction
##
## data:  c(972, 10) out of c(20409, 859)
## X-squared = 23.426, df = 1, p-value = 1.298e-06
## alternative hypothesis: two.sided
## 95 percent confidence interval:
##  0.02763258 0.04433663
## sample estimates:
##     prop 1     prop 2
## 0.04762605 0.01164144
```

Next we plot the ecoregion data to create Fig. 2A:

```
Fig2A <- ggplot(ecoregionKierCompadre %>% arrange(ecoregionAbb,DataSource),
aes(x = ecoregionAbb, y = value, fill = DataSource)) +
  geom_bar(stat = "identity", size = 2, position = position_dodge()) +
  scale_x_discrete(limits = c("Trop", "Temp", "Med/Des", "Tund/Bor", "Wetl"))
```

```
+
  scale_fill_manual(
    values = c("black", "#25b7bc"),
    labels = c(" Nature", " COMPADRE")
  ) +
  theme(
    panel.grid.minor.x = element_blank(),
    panel.grid.major.x = element_blank()
  ) +
  xlab("Ecoregion") +
  ylab("Percentage of species") +
  NULL

Fig2A
```

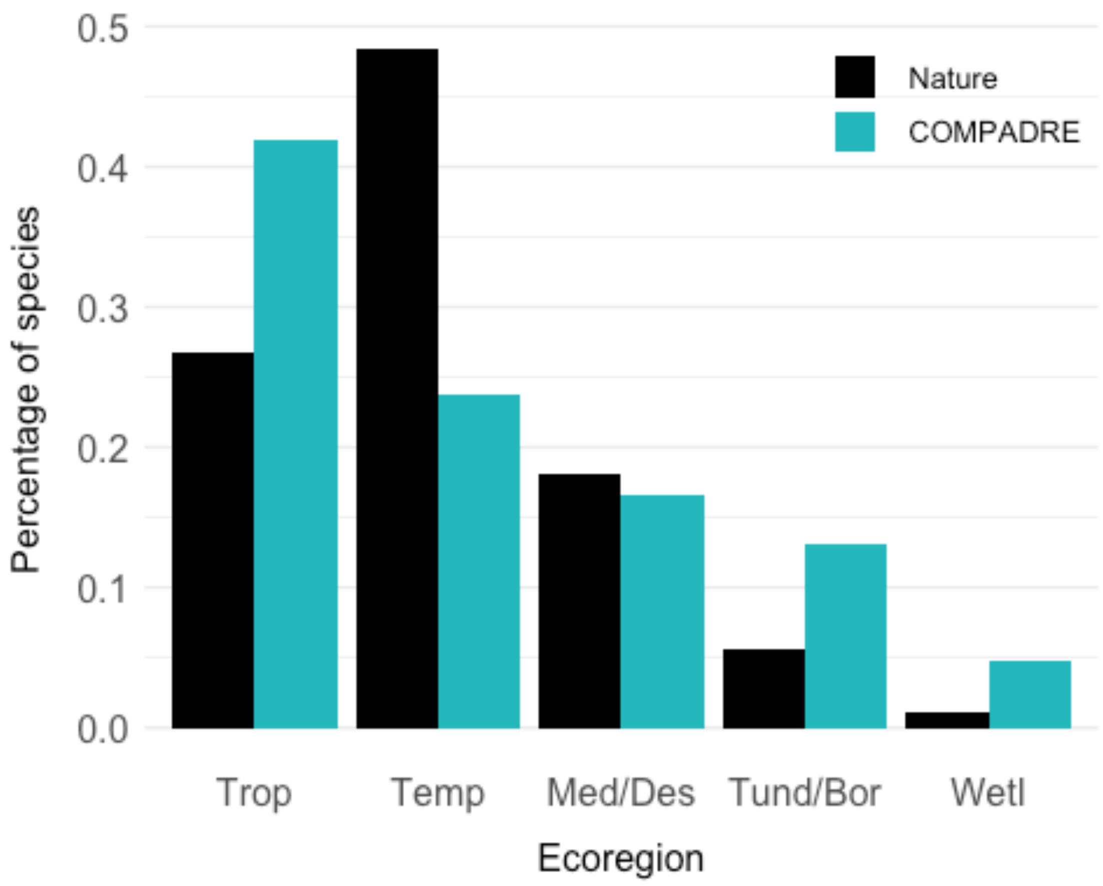


#### Length of study vs. ecoregion and organism type

Here we test whether the length of a study is associated with the ecoregion (i.e., do some studies in some regions tend to be longer/shorter than others?) and with organism type (i.e., do some types of organism tend to be studied for longer than others?)

```
tempDB <- db %>%
  select(Study, ecoregionLabel, OrganismType, StudyStart, StudyEnd) %>%
```

```
  unique() %>%
  na.omit() %>%
  mutate(nyrs = as.numeric(StudyEnd) - as.numeric(StudyStart))

mod1 <- glm(nyrs ~ ecoregionLabel, data = tempDB, family = "poisson")
anova(mod1, test = "Chi")

## Analysis of Deviance Table
##
## Model: poisson, link: log
##
## Response: nyrs
##
## Terms added sequentially (first to last)
##
##
##                Df Deviance Resid. Df Resid. Dev  Pr(>Chi)
## NULL                             664     2782.1
## ecoregionLabel  5   159.55       659     2622.6 < 2.2e-16 ***
## ---
## Signif. codes:  0 '***' 0.001 '**' 0.01 '*' 0.05 '.' 0.1 ' ' 1

mod1 <- glm(nyrs ~ OrganismType, data = tempDB, family = "poisson")
anova(mod1, test = "Chi")

## Analysis of Deviance Table
##
## Model: poisson, link: log
##
## Response: nyrs
##
## Terms added sequentially (first to last)
##
##
##              Df Deviance Resid. Df Resid. Dev  Pr(>Chi)
## NULL                           664     2782.1
## OrganismType 11   165.42       653     2616.7 < 2.2e-16 ***
## ---
## Signif. codes:  0 '***' 0.001 '**' 0.01 '*' 0.05 '.' 0.1 ' ' 1
```

##### Distribution across continents

We first check whether we find species in the database associated with several continents, or with other issues.

```
table(db$Continent,useNA = 'always')

##
##     Africa       Asia     Europe      FRANK        LAB  N America         NC 
Oceania
##        192        248       2996          4         12       4864        120 
```

```
258
## S America      <NA>
##       419          8
```

We can filter the database to exclude studies with unknown continent (NA and NC), laboratory studies (LAB), and studies with "Frankenstein" matrices (FRANK) (these are matrices that have been parameterised from disparate populations) and rename the continent variables:

```
# remove rows with continent = LAB, FRANK or NC
Continent <- db %>%
  select(Study, SpeciesAccepted, SpeciesAuthor, Continent, YearPublication)
%>%
  filter(!(Continent %in% c("LAB", "FRANK","NC"))) %>%
  filter(!is.na(Continent)) %>%
  group_by(Continent) %>%
  summarize(n = length(unique(SpeciesAuthor))) %>%
  mutate(continentLabel = Continent) %>%
  mutate(continentLabel = ifelse(continentLabel ==
"Europe","Euro",continentLabel)) %>%
  mutate(continentLabel = ifelse(continentLabel == "N America","N
Am",continentLabel)) %>%
  mutate(continentLabel = ifelse(continentLabel ==
"Oceania","Ocea",continentLabel)) %>%
  mutate(continentLabel = ifelse(continentLabel == "S America","S
Am",continentLabel))
```

Now, we normalise the data by size of the continent (surface area in km^2) and process the dataset to obtain studies per million square km for plotting:

```
continent_size <- data.frame(
  continentLabel = c("Africa", "Asia", "Euro", "N Am", "Ocea", "S Am"),
  size_km2 = c(30370000, 44579000, 10180000, 24709000, 8525989, 17850000)
)

studyDensityByContinent <- left_join(Continent, continent_size) %>%
  mutate(n_norm = ((n / size_km2))) %>%
  mutate(size_millionkm2 = size_km2 / 1000000) %>%
  mutate(studiesPerMillionkm2 = n / size_millionkm2) %>%
  arrange(-studiesPerMillionkm2) %>%
  select(Continent,continentLabel, studiesPerMillionkm2)

## Joining, by = "continentLabel"
```

We plot the data to create Fig. 2B:

```
Fig2B <- ggplot(data = studyDensityByContinent, aes(x =
reorder(continentLabel,-studiesPerMillionkm2), y = studiesPerMillionkm2)) +
  geom_bar(stat = "identity", size = 2, fill = "#25b7bc") +
  theme(
    panel.grid.minor.x = element_blank(),
```

```
    panel.grid.major.x = element_blank()
  ) +
  xlab("Continent") +
  ylab("Number of studies per 1 M " ~ km^2)
```

```
Fig2B
```

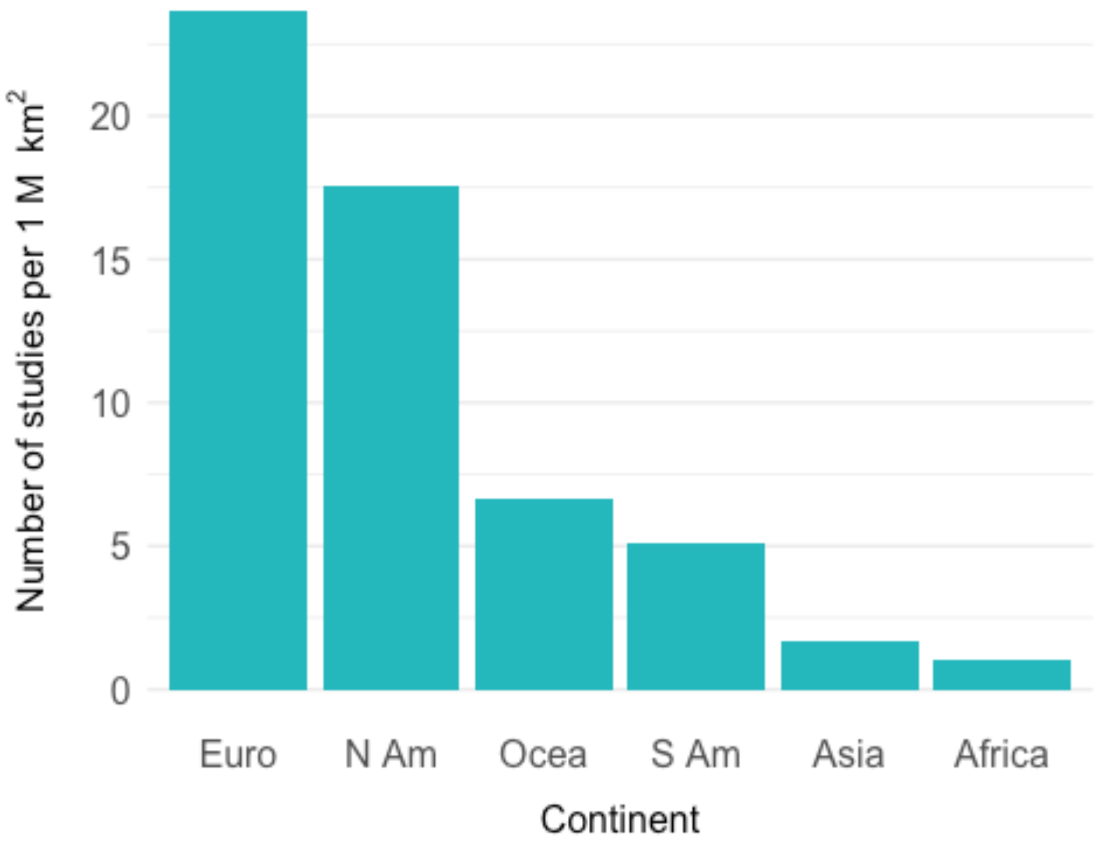


##### Continent data in light of different time frames

We checked if the dominance of Europe and North America changed over time (before 2000 vs. 2000 and onwards):

```
Continent_Pre2000Studies <- db %>%
  filter(YearPublication < 2000) %>%
  select(Study, SpeciesAccepted, SpeciesAuthor, Continent, YearPublication)
%>%
  filter(!(Continent %in% c("LAB", "FRANK","NC"))) %>%
  filter(!is.na(Continent)) %>%
  group_by(Continent) %>%
  summarize(n = length(unique(SpeciesAuthor))) %>%
  mutate(continentLabel = Continent) %>%
  mutate(continentLabel = ifelse(continentLabel ==
"Europe","Euro",continentLabel)) %>%
```

```
  mutate(continentLabel = ifelse(continentLabel == "N America","N
Am",continentLabel)) %>%
  mutate(continentLabel = ifelse(continentLabel ==
"Oceania","Ocea",continentLabel)) %>%
  mutate(continentLabel = ifelse(continentLabel == "S America","S
Am",continentLabel))


Continent_from2000Studies <- db %>%
  filter(YearPublication >= 2000) %>%
  select(Study, SpeciesAccepted, SpeciesAuthor, Continent, YearPublication)
%>%
  filter(!(Continent %in% c("LAB", "FRANK","NC"))) %>%
  filter(!is.na(Continent)) %>%
  group_by(Continent) %>%
  summarize(n = length(unique(SpeciesAuthor))) %>%
  mutate(continentLabel = Continent) %>%
  mutate(continentLabel = ifelse(continentLabel ==
"Europe","Euro",continentLabel)) %>%
  mutate(continentLabel = ifelse(continentLabel == "N America","N
Am",continentLabel)) %>%
  mutate(continentLabel = ifelse(continentLabel ==
"Oceania","Ocea",continentLabel)) %>%
  mutate(continentLabel = ifelse(continentLabel == "S America","S
Am",continentLabel))
```

We then compare all three time frames:

```
# transform both version of continent to percentages for better comparison.
Continent_all <- Continent %>%
  mutate(percent_all = (n / sum(n)) * 100) %>%
  select(continentLabel,percent_all)

Continent_Pre2000Studies <- Continent_Pre2000Studies %>%
  mutate(percent_before2000 = (n / sum(n)) * 100) %>%
  select(continentLabel,percent_before2000)

Continent_from2000Studies <- Continent_from2000Studies %>%
  mutate(percent_2000_onwards = (n / sum(n)) * 100)%>%
  select(continentLabel,percent_2000_onwards)

continentTimeFrameComparison <- left_join(left_join(Continent_all,
Continent_Pre2000Studies),
                  Continent_from2000Studies)

## Joining, by = "continentLabel"
## Joining, by = "continentLabel"
```

We compare these distributions visually.

```
ggplot(continentTimeFrameComparison %>%
  pivot_longer(values_to = "percentage",cols =
starts_with("percent"),names_to = "Timeframe") %>%
  mutate(Timeframe = gsub("percent_","",Timeframe)),
  aes(x = continentLabel,y = percentage,fill = Timeframe)) +
  geom_col(position = "dodge") +
  xlab("Continent")
```

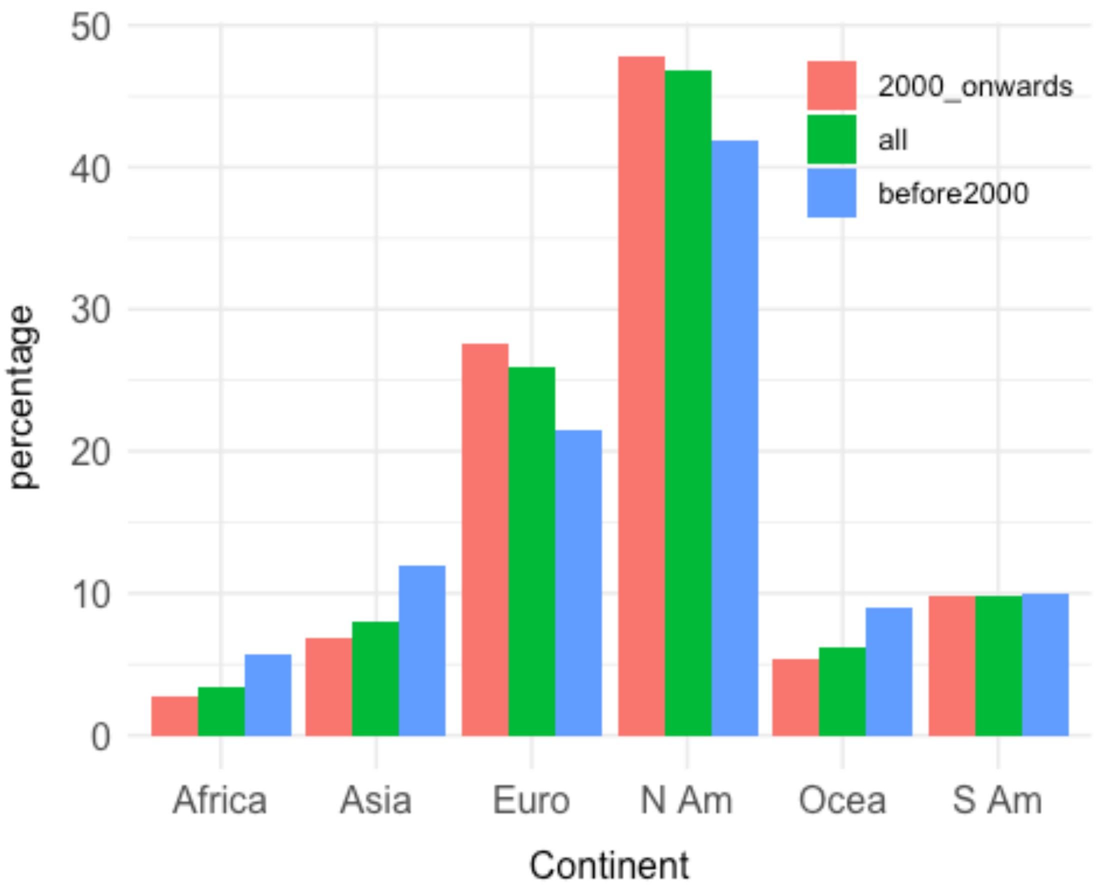


#### Relationship between GDP and number of plant demography studies

To examine the relationship between economy size and the amount of plant demographic studies we downloaded GDP data from the World Bank [2].

We first load these data and process the COMPADRE data as follows:

---

[2] World Bank (2018, October 18). World Development Indicators: GDP per capita (current US$). Retrieved from http://databank.worldbank.org/data/reports.aspx?source=2&type=metadata&series=NY.GDP.PCAP.CD#

```
GDP_worldbank <- read.csv("data/GDPcapita_worldbank_04-11-2018.csv") %>%
  select(Country.Name, Country = Country.Code, GDP2017 = YR2017)
```

We calculate the number of studies per country like this.

```
compadre_studiesPerCountry <- db %>%
  select(Country, Study, SpeciesAccepted, SpeciesAuthor) %>%
  unique() %>%
  group_by(Country) %>%
  summarize(nStudies = length(unique(Study)))
```

We merge the two data frames to get a table including `Country`, `GDP2017` and `n` (number of studies).

```
GDP_and_MPMstudies <- left_join(GDP_worldbank, compadre_studiesPerCountry)
%>%
  mutate(logGDP = log(GDP2017)) %>%
  select(Country, logGDP, nStudies) %>%
  na.omit()

## Joining, by = "Country"
```

We then fit a Poisson generalised linear model (GLM) (log-link) with log-transformed GDP as the explanatory variable and number of demographic studies as the response variable. We also filter outliers

```
mod_gdp <- glm(nStudies ~ logGDP, data = GDP_and_MPMstudies, family =
"poisson")
summary(mod_gdp)

##
## Call:
## glm(formula = nStudies ~ logGDP, family = "poisson", data =
GDP_and_MPMstudies)
##
## Deviance Residuals:
##     Min       1Q   Median       3Q      Max
## -7.2803  -3.3726  -1.0081   0.0432  22.2924
##
## Coefficients:
##             Estimate Std. Error z value Pr(>|z|)
## (Intercept) -3.33166    0.37665  -8.846   <2e-16 ***
## logGDP       0.59406    0.03673  16.175   <2e-16 ***
## ---
## Signif. codes:  0 '***' 0.001 '**' 0.01 '*' 0.05 '.' 0.1 ' ' 1
##
## (Dispersion parameter for poisson family taken to be 1)
##
##     Null deviance: 1494.7  on 55  degrees of freedom
## Residual deviance: 1144.3  on 54  degrees of freedom
## AIC: 1337.7
```

```
##
## Number of Fisher Scoring iterations: 6
```

```
anova(mod_gdp, test = "Chi")
```

```
## Analysis of Deviance Table
##
## Model: poisson, link: log
##
## Response: nStudies
##
## Terms added sequentially (first to last)
##
##
##        Df Deviance Resid. Df Resid. Dev  Pr(>Chi)
## NULL                      55     1494.7
## logGDP  1   350.36        54     1144.3 < 2.2e-16 ***
## ---
## Signif. codes:  0 '***' 0.001 '**' 0.01 '*' 0.05 '.' 0.1 ' ' 1
```

```
# Pseudo-R2
mod_gdp$deviance / mod_gdp$null.deviance
```

```
## [1] 0.765599
```

We plot the obtained data to create Fig. 2C. We first create a data frame to predict from (newDF).

```
newDF <- data.frame(logGDP = seq(min(GDP_and_MPMstudies$logGDP),
max(GDP_and_MPMstudies$logGDP),length.out = 100))

pv <- predict(mod_gdp, newDF, se.fit = TRUE)
newDF$fitted_link <- pv$fit
newDF$se_link <- pv$se.fit

# Get inverse link from the model
invLink <- family(mod_gdp)$linkinv

newDF <- mutate(newDF,
  fit_response = invLink(fitted_link),
  fit_upper = invLink(fitted_link + (2 * se_link)),
  fit_lower = invLink(fitted_link - (2 * se_link))
)

# Get outliers
outliers_studies <- GDP_and_MPMstudies %>%
  filter(nStudies > 50)

outliers_gdp <- GDP_and_MPMstudies %>%
  filter(logGDP == max(logGDP))
```

Now we plot the data and model predictions

```
Fig2C <- ggplot(data = GDP_and_MPMstudies, aes(x = logGDP, y = nStudies)) +
  # add regression
  geom_ribbon(data = newDF, aes(x = logGDP, ymin = fit_lower, ymax =
fit_upper), fill = "grey80", inherit.aes = FALSE) +
  geom_line(data = newDF, aes(x = logGDP, y = fit_response), colour =
"#25b7bc", inherit.aes = FALSE) +
  # add points
  geom_point() +
  # Modify to put outliers (Mexico and USA) as arrows.
  ylim(0, 45) +
  geom_segment(
    data = outliers_studies, aes(x = logGDP, y = 40, xend = logGDP, yend =
45),
    arrow = arrow(end = "last", length = unit(0.3, "cm"), type = "closed")
  ) +
  geom_text(data = outliers_studies, aes(x = logGDP, y = 37, label =
paste0(Country, " (", nStudies, " studies)")), colour = "#25b7bc") +
  # Modify add GDP outlier (LUX)
  geom_text(data = outliers_gdp, aes(x = logGDP, y = 0, label = Country),
colour = "#25b7bc") +
  xlab("log(GDP per capita)") +
  ylab("Number of studies") +
  NULL

Fig2C
```

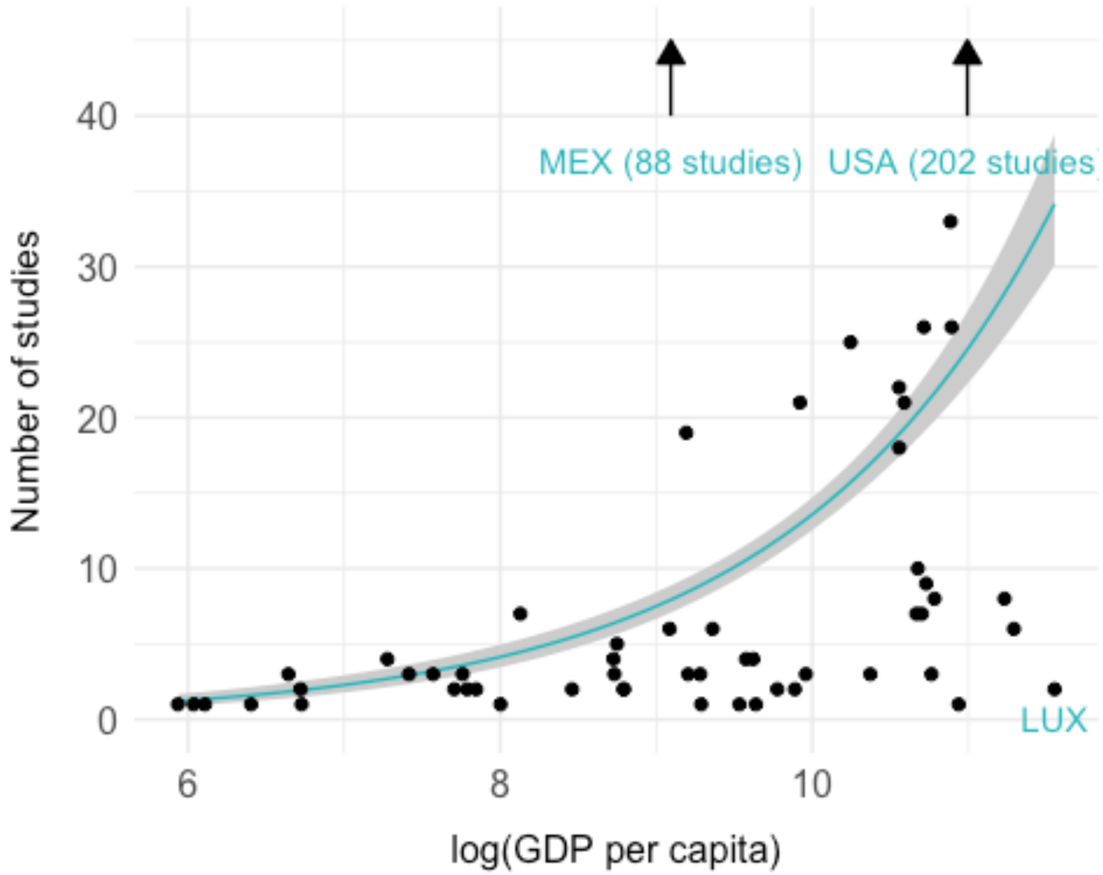


We double check if the above obtained outliers influence the results of our analyses:

```
# re-run regression without outliers
mod_gdp_noOutliers <- glm(nStudies ~ logGDP, data = GDP_and_MPMstudies %>%
                filter(!Country %in% c("USA","MEX","LUX")), family =
"poisson")
summary(mod_gdp_noOutliers)

##
## Call:
## glm(formula = nStudies ~ logGDP, family = "poisson", data =
GDP_and_MPMstudies %>%
##     filter(!Country %in% c("USA", "MEX", "LUX")))
##
## Deviance Residuals:
##     Min       1Q   Median       3Q      Max
## -4.5012  -1.4118  -0.4568   0.3909   4.5095
##
## Coefficients:
##             Estimate Std. Error z value Pr(>|z|)
## (Intercept) -3.08487    0.47736  -6.462 1.03e-10 ***
## logGDP       0.52164    0.04736  11.016  < 2e-16 ***
## ---
```

```
## Signif. codes:  0 '***' 0.001 '**' 0.01 '*' 0.05 '.' 0.1 ' ' 1
##
## (Dispersion parameter for poisson family taken to be 1)
##
##     Null deviance: 385.19  on 52  degrees of freedom
## Residual deviance: 228.79  on 51  degrees of freedom
## AIC: 406.11
##
## Number of Fisher Scoring iterations: 5
```

```
# Pseudo-R2
mod_gdp_noOutliers$deviance / mod_gdp_noOutliers$null.deviance
```

```
## [1] 0.5939757
```

```
newDF <- data.frame(logGDP = seq(
  min(GDP_and_MPMstudies %>%
        filter(!Country %in% c("USA","MEX","LUX")) %>%
        pull(logGDP)),
  max(GDP_and_MPMstudies %>%
        filter(!Country %in% c("USA","MEX","LUX")) %>%
        pull(logGDP)),length.out = 100))

pv <- predict(mod_gdp_noOutliers, newDF, se.fit = TRUE)
newDF$fitted_link <- pv$fit
newDF$se_link <- pv$se.fit

# Get inverse link from the model
invLink <- family(mod_gdp_noOutliers)$linkinv

newDF <- mutate(newDF,
  fit_response = invLink(fitted_link),
  fit_upper = invLink(fitted_link + (2 * se_link)),
  fit_lower = invLink(fitted_link - (2 * se_link))
)

# Plot
Fig2C_noOutliers <- ggplot(GDP_and_MPMstudies %>%
                filter(!Country %in% c("USA","MEX","LUX")), aes(x = logGDP, y
= nStudies)) +
  geom_point() +
  # add regression
  geom_ribbon(data = newDF, aes(x = logGDP, ymin = fit_lower, ymax =
fit_upper), inherit.aes = FALSE, fill = "#D8D8D880") +
  geom_line(data = newDF, aes(x = logGDP, y = fit_response), colour =
"#25b7bc") +
  xlab("log transformed GDP (in 2017)") +
  ylab("Number of studies") +
  NULL
```

```
Fig2C_noOutliers
```

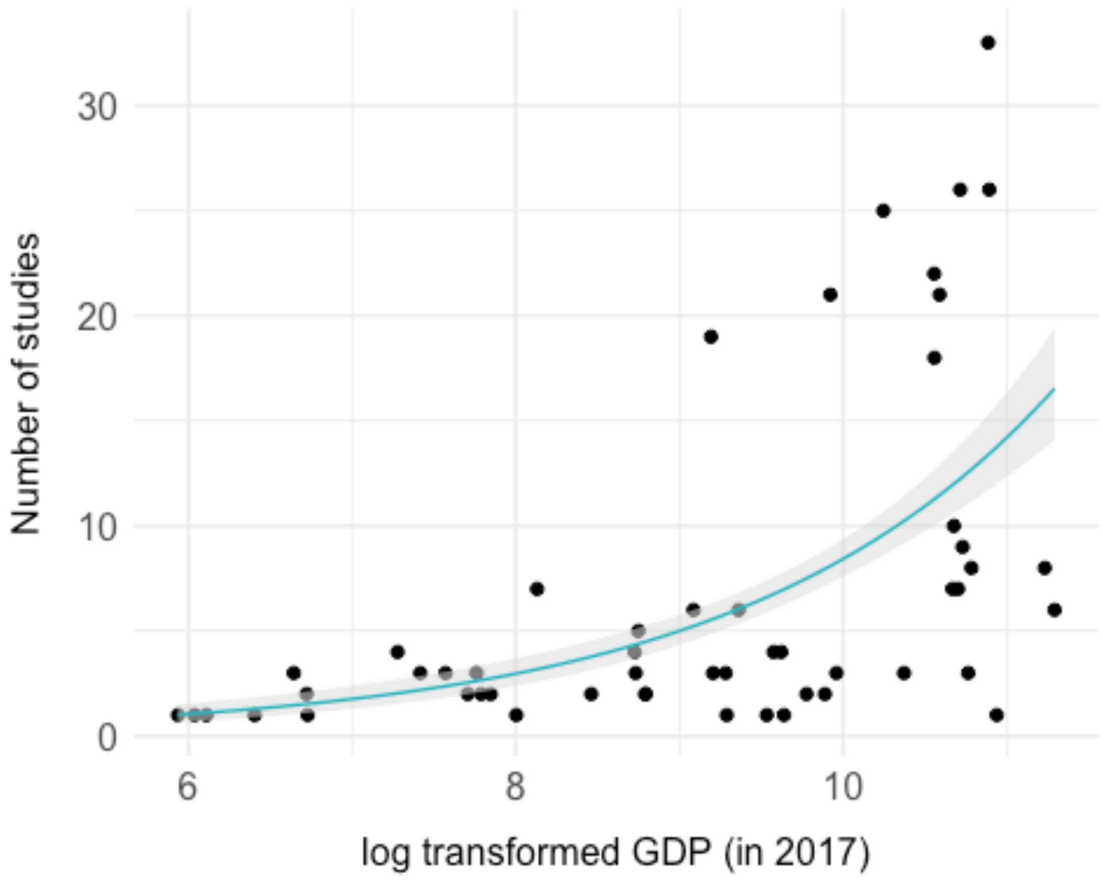


The outliers do not influence the result qualitatively.

#### D) FOR WHICH SPECIES AND POPULATIONS DO WE HAVE DEMOGRAPHIC DATA?

This section focusses on WHAT species and populations have been studied.

This includes (i) Several aspects of the taxonomy of the plants; (ii) a comparison of IUCN Red List conservation status; and (iii), an analysis of population growth rates (lambda distribution and lambda regression towards the mean)

##### Taxonomy

We analyse several aspects of the taxonomy of the species stored in COMPADRE (growth form, Angiosperms vs Gymnosperms, monocots vs. euicots, distribution among the five largest dicot families, distribution among the growth forms according to Raunkiær).

##### Angioperm vs. Gymnosperm representation

We obtained data about the distribution of angiosperms, gymnosperms and non-seed plants in nature from Campbell et al. (2018) [3] which we then compared to the data in COMPADRE.

We first process the data as follows.

```
AngioGymno <- db %>%
  select(Study, SpeciesAccepted, SpeciesAuthor, AngioGymno) %>%
  filter(!is.na(AngioGymno)) %>%
  unique() %>%
  group_by(AngioGymno) %>%
  summarise(compadre_nSpecies = length(unique(SpeciesAuthor))) %>%
  mutate(AngioGymno = ifelse(AngioGymno == "other", "Non-seed", AngioGymno))
%>%
  mutate(AngioGymno = ifelse(AngioGymno == "Angiosperm", "Angio",
AngioGymno)) %>%
  mutate(AngioGymno = ifelse(AngioGymno == "Gymnosperm", "Gymno",
AngioGymno)) %>%
  mutate(nature_nSpecies = c(200000 + 3000 + 90000, 807, 37300)) %>%
  mutate(percent_nature_nSpecies = (nature_nSpecies / sum(nature_nSpecies)) *
100) %>%
  mutate(percent_compadre_nSpecies = (compadre_nSpecies /
sum(compadre_nSpecies)) * 100)

AngioGymno

## # A tibble: 3 x 5
##   AngioGymno compadre_nSpecies nature_nSpecies percent_nature_…
percent_compadr…
## * <chr>                  <int>           <dbl>            <dbl>
<dbl>
## 1 Angio                    850          293000           88.5
91.2
## 2 Gymno                     55             807            0.244
5.90
## 3 Non-seed                  27           37300           11.3
2.90
```

We then test statistically whether the distribution in COMPADRE is different from in nature.

```
chisq.test(x = AngioGymno$compadre_nSpecies, p =
AngioGymno$percent_nature_nSpecies / 100) # all
```

---

[3] Campbell, N. A., Urry, L. A., Cain, M. L., Wasserman, S. A., Minorsky, P. V., & Reece, J. B. (2018). Biology: A global approach (Eleventh edition, global edition). Boston: Pearson

```
##
##  Chi-squared test for given probabilities
##
## data:  AngioGymno$compadre_nSpecies
## X-squared = 1282.7, df = 2, p-value < 2.2e-16
```

The distribution of species among Angiosperm, Gymnosperm and non-seed plants is significantly different in COMPADRE compared to distribution in nature.

Now we test where those differences lie. The number of species in COMPADRE (with this metadata) is 932 and the number of species (estimated) in nature is 3.31107^{5}

```
# Angio
prop.test(x = c(850, 293000), n = c(932, 331107))

##
##  2-sample test for equality of proportions with continuity correction
##
## data:  c(850, 293000) out of c(932, 331107)
## X-squared = 6.4455, df = 1, p-value = 0.01112
## alternative hypothesis: two.sided
## 95 percent confidence interval:
##  0.00835026 0.04586344
## sample estimates:
##    prop 1    prop 2
## 0.9120172 0.8849103

# Gymno
prop.test(x = c(55,807), n = c(932, 331107))

##
##  2-sample test for equality of proportions with continuity correction
##
## data:  c(55, 807) out of c(932, 331107)
## X-squared = 1127.1, df = 1, p-value < 2.2e-16
## alternative hypothesis: two.sided
## 95 percent confidence interval:
##  0.04090784 0.07224335
## sample estimates:
##      prop 1      prop 2
## 0.059012876 0.002437279

# Non-seed
prop.test(x = c(27,37300), n = c(932, 331107))

##
##  2-sample test for equality of proportions with continuity correction
##
## data:  c(27, 37300) out of c(932, 331107)
## X-squared = 64.39, df = 1, p-value = 1.021e-15
## alternative hypothesis: two.sided
```

```
## 95 percent confidence interval:
##  -0.09504204 -0.07232285
## sample estimates:
##     prop 1     prop 2
## 0.02896996 0.11265241
```

Next, we plot the data to obtain Fig. 2D:

```
(AngioGymnoPlotData <- AngioGymno %>%
  select(AngioGymno, starts_with("percent")) %>%
  pivot_longer(cols = ends_with("nSpecies"), names_to = "database", values_to
= "sp_perc") %>%
  mutate(database = ifelse(database == "percent_compadre_nSpecies",
"COMPADRE", "Nature")) %>%
  arrange(database))

## # A tibble: 6 x 3
##   AngioGymno database sp_perc
##   <chr>      <chr>      <dbl>
## 1 Angio      COMPADRE  91.2
## 2 Gymno      COMPADRE   5.90
## 3 Non-seed   COMPADRE   2.90
## 4 Angio      Nature    88.5
## 5 Gymno      Nature     0.244
## 6 Non-seed   Nature    11.3

Fig2D <- ggplot(AngioGymnoPlotData, aes(
  x = fct_rev(AngioGymno), y = sp_perc,
  fill = fct_rev(database)
)) +
  geom_col(position = "dodge") +
  scale_fill_manual(
    values = c("black", "#25b7bc"),
    labels = c(" Nature", " COMPADRE")
  ) +
  scale_y_continuous(limits = c(0, 100)) +
  theme(
    panel.grid.minor.x = element_blank(),
    panel.grid.major.x = element_blank(),
    legend.justification = c(0.1, 1), legend.position = c(0.1, 1)
  ) +
  # ggtitle("plant divisions")+
  xlab("") +
  ylab("Percentage of species")

Fig2D
```

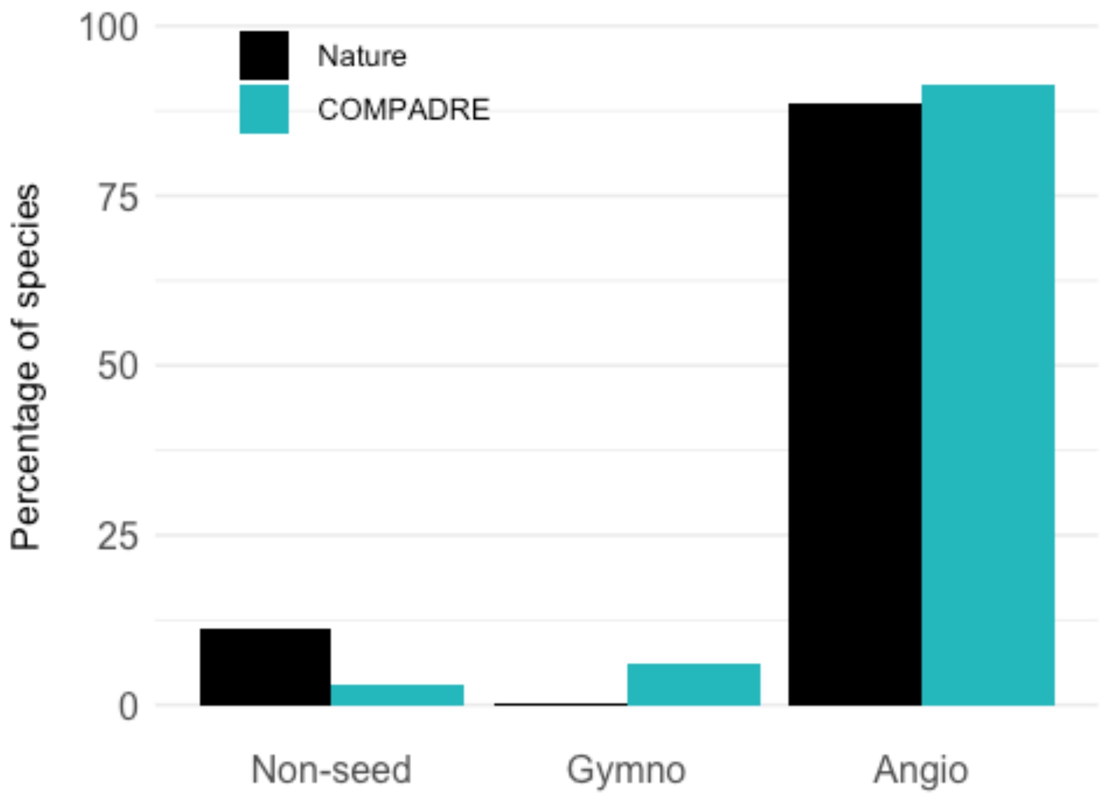


##### Monocot vs. Dicot

We obtained data about the distribution of eudicots and monocots in nature from Evert et al. (2013) [4] which we then compared to the data in COMPADRE.

We then process the data in a similar way as above:

```
DicotMonoc <- db %>%
  select(Study, SpeciesAccepted, SpeciesAuthor, DicotMonoc, AngioGymno) %>%
  filter(AngioGymno == "Angiosperm") %>%
  group_by(DicotMonoc) %>%
  summarise(CompadreData = length(unique(SpeciesAuthor))) %>%
  mutate(nature =  c(200000 + 3000, 90000)) %>%
  mutate(percent_COMPADRE = (CompadreData/sum(CompadreData))*100) %>%
  mutate(percent_nature = (nature/sum(nature))*100)
DicotMonoc

## # A tibble: 2 x 5
##   DicotMonoc CompadreData nature percent_COMPADRE percent_nature
```

[4] Evert, R. F. & Eichhorn, S. E. (2013). Raven biology of plants (Eighth edition). New York: W.H. Freeman and Company Publishers.

```
## * <chr>              <int>  <dbl>             <dbl>            <dbl>
## 1 Eudicot              631 203000              74.2             69.3
## 2 Monocot              219  90000              25.8             30.7
```

We then run statistical tests on the data. The number of species with this metadata in COMPADRE is 850, and the number (estimated) in nature is 2.93^{5}

```
# all
chisq.test(x = DicotMonoc$CompadreData, p = DicotMonoc$percent_nature/100)

##
##  Chi-squared test for given probabilities
##
## data:  DicotMonoc$CompadreData
## X-squared = 9.7944, df = 1, p-value = 0.00175
```

Next, we plot the data to obtain Fig. 2E:

```
DicotMonocPlotData <- DicotMonoc %>%
  select(DicotMonoc,starts_with("percent")) %>%
  pivot_longer(cols = starts_with("percent"),values_to =
"percent_species",names_to = "database") %>%
  mutate(database = gsub("percent_","",database)) %>%
  mutate(database = gsub("nature","Nature",database))



Fig2E <- ggplot(DicotMonocPlotData,aes(x = fct_rev(DicotMonoc), y =
percent_species, fill = fct_rev(database))) +
  geom_col(position = "dodge") +
  scale_fill_manual(
    values = c("black", "#25b7bc"),
    labels = c(" Nature", " COMPADRE")
  ) +
  scale_y_continuous(limits = c(0, 100)) +
  theme(
    panel.grid.minor.x = element_blank(),
    panel.grid.major.x = element_blank(),
    legend.justification = c(0.1, 1), legend.position = c(0.1, 1)
  ) +
  xlab("") +
  ylab("Percentage of species")

Fig2E
```

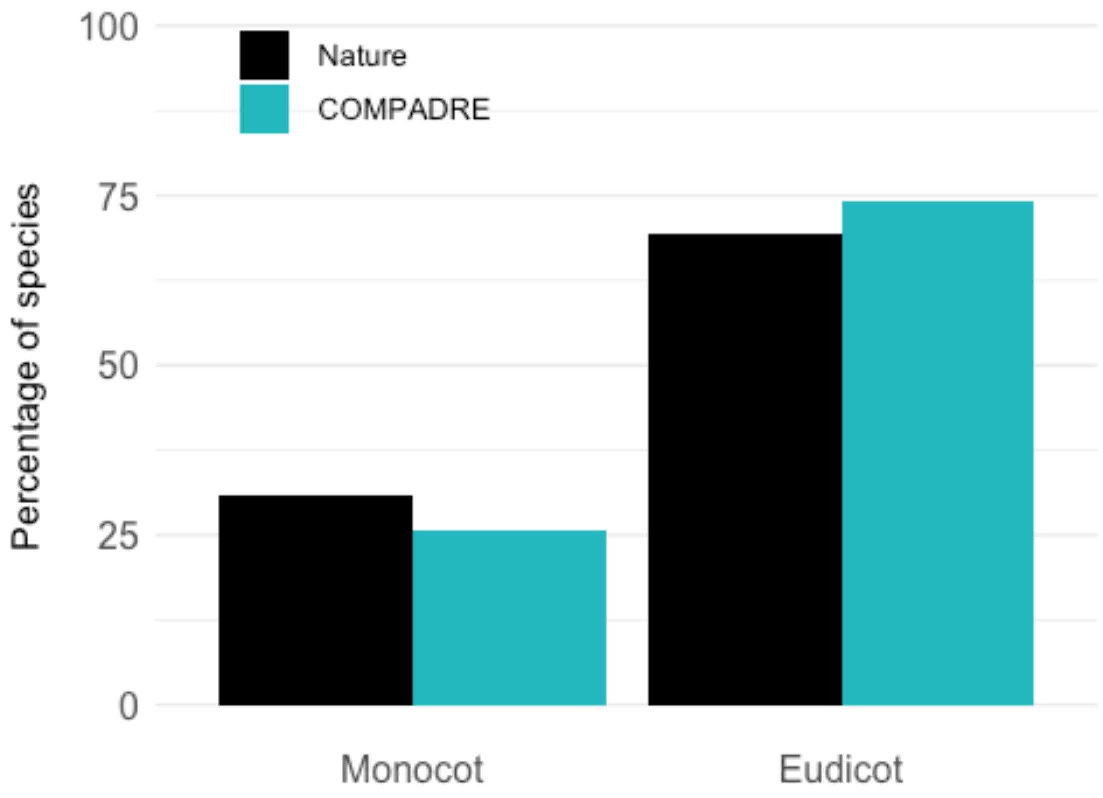


##### Top 5 taxonomic families

We downloaded a list of species and according families from The Plant List (2010) [5] and merged it with the data in COMPADRE.

```
familiesWORLD <- read.csv("data/TotalSpeciesList.csv") %>%
  select(Family, Species) %>%
  group_by(Family) %>%
  summarise(n = length(Species))

familiesCOMPADRE <- db %>%
  select(Family, SpeciesAuthor) %>%
  unique() %>%
  group_by(Family) %>%
  summarise(n = length(SpeciesAuthor))

familiesBOTH <- left_join(familiesWORLD, familiesCOMPADRE, by = "Family") %>%
  rename(PlantList = n.x) %>%
  rename(CompadreData = n.y) %>%
```

---

[5] The Plant List (2010). The Plant List: A working list of all known plant species. Version 1.1. Retrieved from http://www.theplantlist.org

```
  mutate(CompadreData = ifelse(is.na(CompadreData),0,CompadreData)) %>%
  arrange(-PlantList) %>%
  mutate(percPlantList = PlantList/sum(PlantList)) %>%
  mutate(percCompadreData = CompadreData/sum(CompadreData))

head(familiesBOTH)
```

```
## # A tibble: 6 x 5
##   Family      PlantList CompadreData percPlantList percCompadreData
##   <chr>           <int>        <dbl>         <dbl>            <dbl>
## 1 Compositae      62199           85        0.105           0.0958
## 2 Leguminosae     42836           64        0.0725          0.0722
## 3 Orchidaceae     31800           58        0.0538          0.0654
## 4 Rosaceae        21382           18        0.0362          0.0203
## 5 Rubiaceae       15486            4        0.0262          0.00451
## 6 Poaceae         14599           38        0.0247          0.0428
```

```
Fig2F <- ggplot(familiesBOTH %>% slice(1:5) %>%
  select(Family, starts_with("perc")) %>%
  pivot_longer(cols = -Family,values_to = "perc",names_to = "database") %>%
  mutate(source = gsub("perc","",database)),
  aes(x = Family, y = perc, fill = fct_rev(database))) +
  geom_col(position = "dodge") +
  scale_fill_manual(
    values = c("black", "#25b7bc"),
    labels = c(" Nature", " COMPADRE")
  ) +
  scale_x_discrete(labels = c("Comp", "Legum", "Orchid", "Rosa", "Rubia")) +
  theme(
    panel.grid.minor.x = element_blank(),
    panel.grid.major.x = element_blank()
  ) +
  xlab("") +
  ylab("Percentage of species")

Fig2F
```

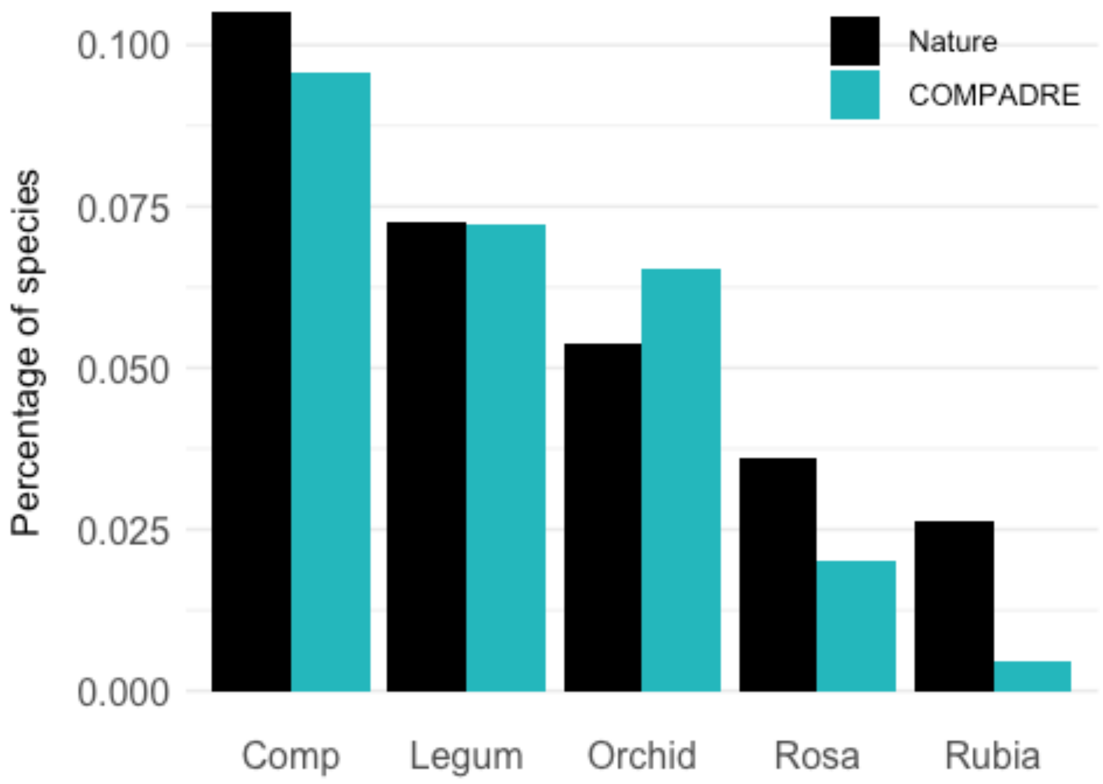


We then subset the data to only include the five biggest famlies (by number of species):

```
# only use Top5 families from PlantList
(top5 <- familiesBOTH %>% slice(1:5))

## # A tibble: 5 x 5
##   Family      PlantList CompadreData percPlantList percCompadreData
##   <chr>           <int>        <dbl>         <dbl>            <dbl>
## 1 Compositae      62199           85        0.105           0.0958
## 2 Leguminosae     42836           64        0.0725          0.0722
## 3 Orchidaceae     31800           58        0.0538          0.0654
## 4 Rosaceae        21382           18        0.0362          0.0203
## 5 Rubiaceae       15486            4        0.0262          0.00451
```

We then run statistical tests to compare the distribution in COMPADRE with the distribution in nature (from The Plant List). The number of species in the Top 5 list in COMPADRE is 229 and the equivalent number in the Plant List is 173703.

```
# all
chisq.test(x = top5$CompadreData, p = (top5$PlantList / sum(top5$PlantList)))

##
##  Chi-squared test for given probabilities
##
```

```
## data:  top5$CompadreData
## X-squared = 24.161, df = 4, p-value = 7.417e-05

#Compositae
prop.test(x = c(85, 62199), n = c(229, 173703))

##
##  2-sample test for equality of proportions with continuity correction
##
## data:  c(85, 62199) out of c(229, 173703)
## X-squared = 0.11856, df = 1, p-value = 0.7306
## alternative hypothesis: two.sided
## 95 percent confidence interval:
##  -0.05169737  0.07790201
## sample estimates:
##    prop 1    prop 2
## 0.3711790 0.3580767

#Leguminosae
prop.test(x = c(64, 42836), n = c(229, 173703))

##
##  2-sample test for equality of proportions with continuity correction
##
## data:  c(64, 42836) out of c(229, 173703)
## X-squared = 1.1589, df = 1, p-value = 0.2817
## alternative hypothesis: two.sided
## 95 percent confidence interval:
##  -0.02747064  0.09321293
## sample estimates:
##    prop 1    prop 2
## 0.2794760 0.2466048

# Orchidaceae
prop.test(x = c(58, 31800), n = c(229, 173703))

##
##  2-sample test for equality of proportions with continuity correction
##
## data:  c(58, 31800) out of c(229, 173703)
## X-squared = 7.0719, df = 1, p-value = 0.00783
## alternative hypothesis: two.sided
## 95 percent confidence interval:
##  0.01166263 0.12874538
## sample estimates:
##    prop 1    prop 2
## 0.2532751 0.1830711

# Rosaceae
prop.test(x = c(18, 21382), n = c(229, 173703))
```

```
##
##  2-sample test for equality of proportions with continuity correction
##
## data:  c(18, 21382) out of c(229, 173703)
## X-squared = 3.7936, df = 1, p-value = 0.05145
## alternative hypothesis: two.sided
## 95 percent confidence interval:
##  -0.081568642 -0.007416454
## sample estimates:
##     prop 1     prop 2
## 0.07860262 0.12309517
```

```
# Rubiaceae
prop.test(x = c(4, 15486), n = c(229, 173703))
```

```
##
##  2-sample test for equality of proportions with continuity correction
##
## data:  c(4, 15486) out of c(229, 173703)
## X-squared = 13.616, df = 1, p-value = 0.0002243
## alternative hypothesis: two.sided
## 95 percent confidence interval:
##  -0.09089148 -0.05247837
## sample estimates:
##     prop 1     prop 2
## 0.01746725 0.08915217
```

##### OrganismType

We check the distribution of species across different growth forms. We first process the data as follows.

```
# Lump Algae, Fern, Liana and Bryophyte into one category
GrowthUnique <- db %>%
  select(SpeciesAccepted, OrganismType) %>%
  mutate(OrganismType = ifelse(is.na(OrganismType), "others", OrganismType))
%>%
  mutate(OrganismType = fct_lump(OrganismType, n = 7)) %>%
  group_by(OrganismType) %>%
  summarize(n = length(unique(SpeciesAccepted))) %>%
  mutate(OrganismTypeLabel =  c("An", "Epi", "Herb", "Palm", "Shrub", "Suc",
"Tree", "Other")) %>%
  arrange(desc(n)) %>%
  mutate(perc = 100*n/sum(n))

GrowthUnique
```

```
## # A tibble: 8 x 4
##   OrganismType     n OrganismTypeLabel  perc
##   <fct>        <int> <chr>             <dbl>
## 1 Herb           370 Herb               48.6
```

```
## 2 Tree           125 Tree           16.4
## 3 Shrub           74 Shrub           9.72
## 4 Succulent       49 Suc             6.44
## 5 Palm            43 Palm            5.65
## 6 Epiphyte        39 Epi             5.12
## 7 Annual          37 An              4.86
## 8 Other           24 Other           3.15
```

We plot the data to obtain Fig. 2G.

```
Fig2G <- ggplot(data = GrowthUnique, aes(x = reorder(OrganismTypeLabel, -n),
y = n)) +
  geom_bar(stat = "identity", size = 2, fill = "#25b7bc") +
  theme(axis.text.x = element_text(size = 10)) +
  theme(
    panel.grid.minor.x = element_blank(),
    panel.grid.major.x = element_blank()
  ) +
  xlab("") +
  ylab("Number of species")

Fig2G
```

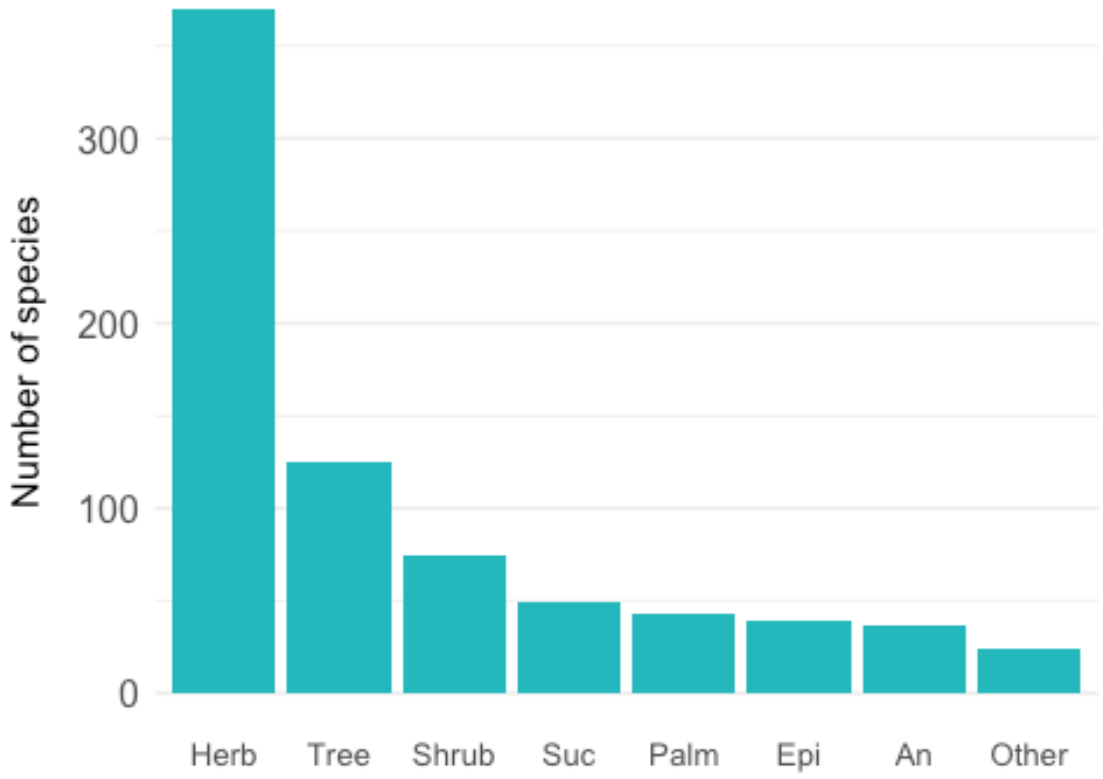

#### Conservation status

We downloaded a list of species and according conservation status from The IUCN Red List (2019) [6] and merged it with the data in COMPADRE.

We recode the deprecated classifications "LR/cd" and "LR/nt" as NT and "LR/lc" to LC.

```
IUCNstatusData <- read.csv("data/IUCNdataRAW.csv", header = TRUE) %>%
  filter(IUCNstatus != "result.scientific_name") %>%
  mutate(IUCNstatus = ifelse(IUCNstatus %in%
c("LR/cd","LR/nt"),"NT",IUCNstatus)) %>%
  mutate(IUCNstatus = ifelse(IUCNstatus == "LR/lc","LC",IUCNstatus)) %>%
  select(-X) %>%
  mutate(IUCNstatus = as.factor(IUCNstatus)) %>%
  mutate(IUCNstatus  =  fct_relevel(IUCNstatus, c("DD", "LC", "NT", "VU",
"EN", "CR", "EW", "EX"))) %>%
  filter(IUCNstatus!="DD")

IUCNDataSummary <- IUCNstatusData %>%
  group_by(IUCNstatus) %>%
  summarise(nRedList = n()) %>%
  mutate(percRedList = nRedList/sum(nRedList))

IUCNDataSummary

## # A tibble: 7 x 3
##   IUCNstatus nRedList percRedList
## * <fct>         <int>       <dbl>
## 1 LC            51772    0.621
## 2 NT             5944    0.0713
## 3 VU            11039    0.132
## 4 EN             8783    0.105
## 5 CR             5347    0.0641
## 6 EW               41    0.000492
## 7 EX              434    0.00521
```

We get our COMPADRE species list and add the IUCN Red List categories.

```
compadreRedListStatus <- left_join(db %>%
  select(SpeciesAccepted) %>%
  unique(),
  IUCNstatusData) %>%
  mutate(assessed = ifelse(is.na(IUCNstatus),"FALSE","TRUE"))

## Joining, by = "SpeciesAccepted"
```

[6] IUCN 2019. The IUCN Red List of Threatened Species. Version 2019-2. Retrieved from http://www.iucnredlist.org

```
#How many of COMPADRE's species have been assessed?
table(compadreRedListStatus$assessed)

##
## FALSE  TRUE
##   542   218

compadreRedListStatusSummary <- compadreRedListStatus %>%
  group_by(IUCNstatus) %>%
  summarise(nCompadre = n()) %>%
  filter(!is.na(IUCNstatus)) %>%
  mutate(percCompadre = nCompadre/sum(nCompadre))
```

We join the two databases (COMPADER and NATURE) for comparison and plotting.

```
bothDB_RedListSummary <-
left_join(compadreRedListStatusSummary,IUCNDataSummary)

## Joining, by = "IUCNstatus"

bothDB_RedListSummary

## # A tibble: 5 x 5
##   IUCNstatus nCompadre percCompadre nRedList percRedList
##   <fct>          <int>        <dbl>    <int>       <dbl>
## 1 LC               135       0.619     51772      0.621
## 2 NT                16       0.0734     5944      0.0713
## 3 VU                23       0.106     11039      0.132
## 4 EN                31       0.142      8783      0.105
## 5 CR                13       0.0596     5347      0.0641

Fig2H <- ggplot(bothDB_RedListSummary %>%
         select(IUCNstatus,starts_with("perc")) %>%
         pivot_longer(cols = starts_with("perc"),names_to =
"database",values_to = "percSpecies"),
       aes(x = IUCNstatus,y = percSpecies, fill = fct_rev(database))) +
  geom_col(position = "dodge") +
  scale_fill_manual(
    values = c("black", "#25b7bc"),
    labels = c(" Nature", " COMPADRE")
  ) +
  theme(
    panel.grid.minor.x = element_blank(),
    panel.grid.major.x = element_blank()
  ) +
  xlab("IUCN Red List status") +
  ylab("Percentage of species")

Fig2H
```

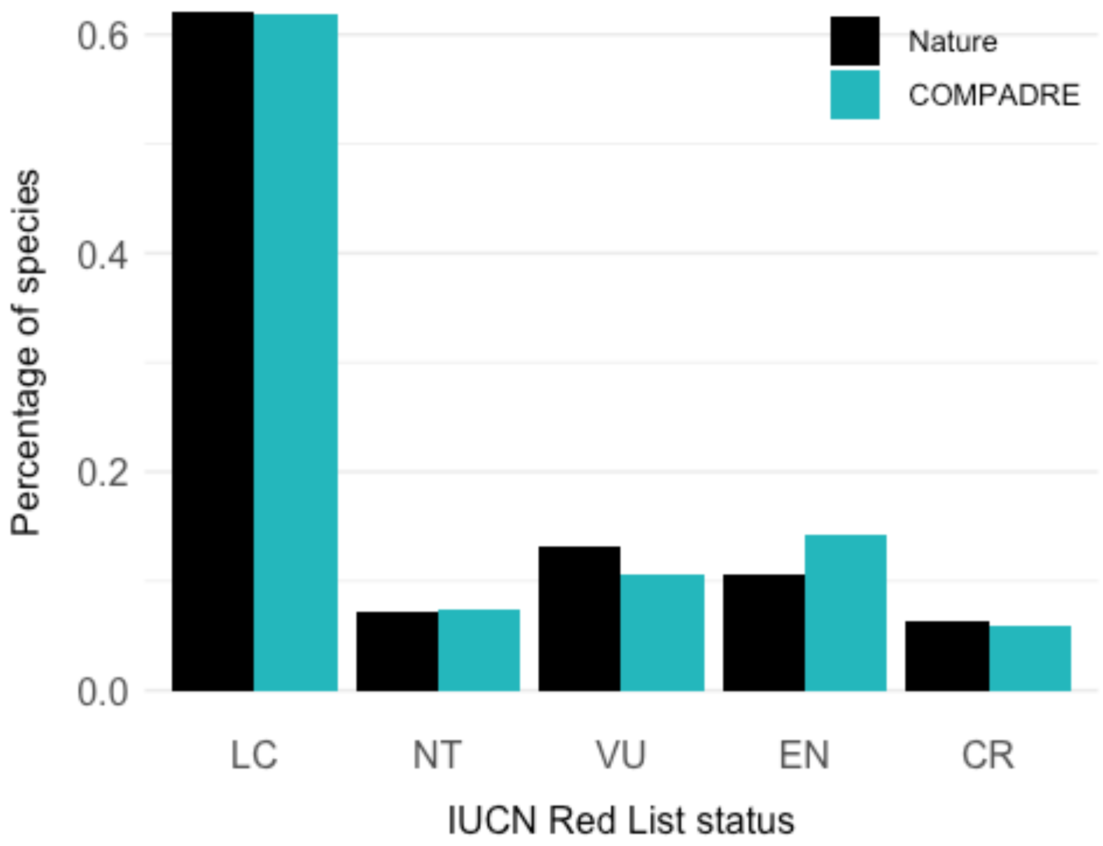


Next, we run statistical tests on the data from the `bothDB_RedListSummary` table above. The total in COMPDADRE is 218 and the total in the IUCN Red List is 82885.

```
#ALL
chisq.test(x = bothDB_RedListSummary$nCompadre,
           p =
bothDB_RedListSummary$nRedList/sum(bothDB_RedListSummary$nRedList) )

##
##  Chi-squared test for given probabilities
##
## data:  bothDB_RedListSummary$nCompadre
## X-squared = 4.0544, df = 4, p-value = 0.3987

totalCompadre <- sum(bothDB_RedListSummary$nCompadre)
totalRedList <- sum(bothDB_RedListSummary$nRedList)

#CR
prop.test(x = c(13, 5347), n = c(totalCompadre, totalRedList))

##
##  2-sample test for equality of proportions with continuity correction
##
## data:  c(13, 5347) out of c(totalCompadre, totalRedList)
```

```
## X-squared = 0.023957, df = 1, p-value = 0.877
## alternative hypothesis: two.sided
## 95 percent confidence interval:
##  -0.03865700  0.02890092
## sample estimates:
##     prop 1     prop 2
## 0.05963303 0.06451107

#EN
prop.test(x = c(31, 8783), n = c(totalCompadre, totalRedList))

##
##  2-sample test for equality of proportions with continuity correction
##
## data:  c(31, 8783) out of c(totalCompadre, totalRedList)
## X-squared = 2.641, df = 1, p-value = 0.1041
## alternative hypothesis: two.sided
## 95 percent confidence interval:
##  -0.01247350  0.08494498
## sample estimates:
##    prop 1    prop 2
## 0.1422018 0.1059661

#VU
prop.test(x = c(23, 11039), n = c(totalCompadre, totalRedList))

##
##  2-sample test for equality of proportions with continuity correction
##
## data:  c(23, 11039) out of c(totalCompadre, totalRedList)
## X-squared = 1.2138, df = 1, p-value = 0.2706
## alternative hypothesis: two.sided
## 95 percent confidence interval:
##  -0.07082486  0.01546497
## sample estimates:
##    prop 1    prop 2
## 0.1055046 0.1331845

#NT
prop.test(x = c(16, 5944), n = c(totalCompadre, totalRedList))

##
##  2-sample test for equality of proportions with continuity correction
##
## data:  c(16, 5944) out of c(totalCompadre, totalRedList)
## X-squared = 1.0871e-27, df = 1, p-value = 1
## alternative hypothesis: two.sided
## 95 percent confidence interval:
##  -0.03466230  0.03802365
## sample estimates:
```

```
##     prop 1     prop 2
## 0.07339450 0.07171382

#LC
prop.test(x = c(135, 51772), n = c(totalCompadre, totalRedList))

##
##  2-sample test for equality of proportions with continuity correction
##
## data:  c(135, 51772) out of c(totalCompadre, totalRedList)
## X-squared = 0.0086762, df = 1, p-value = 0.9258
## alternative hypothesis: two.sided
## 95 percent confidence interval:
##  -0.07219916  0.06148231
## sample estimates:
##    prop 1    prop 2
## 0.6192661 0.6246245

#DD
prop.test(x = c(2, 14686), n = c(totalCompadre, totalRedList))

##
##  2-sample test for equality of proportions with continuity correction
##
## data:  c(2, 14686) out of c(totalCompadre, totalRedList)
## X-squared = 41.034, df = 1, p-value = 1.496e-10
## alternative hypothesis: two.sided
## 95 percent confidence interval:
##  -0.1832310 -0.1527909
## sample estimates:
##      prop 1      prop 2
## 0.009174312 0.177185257
```

#### Population growth rate (Lambda)

We examine the asymptotic population growth rates (λ) to assess whether researchers have a tendency to collect demographic data on growing or declining populations, and whether researchers tend to study populations that are in a "boom phase".

##### Lambda distribution

We first subset the data to those studies for which λ could be calculated (i.e. the MPMs contained no missing values and did not violate ergodicity and irreducibility assumptions). We use the mean matrix for each population.

```
database <- cdb_flag(database)

compadre_sub <- database %>%
  filter(MatrixTreatment == "Unmanipulated") %>%
  filter(MatrixComposite == "Mean") %>%
```

```
  filter(check_NA_A == FALSE) %>%
  filter(check_ergodic == TRUE)
```

We then calculate λ for each matrix in the COMPADRE subset and place these in a data frame (`tibble`) called `allLambdas`.

```
compadre_sub$lambda <- sapply(matA(compadre_sub), popdemo::eigs, what =
"lambda")

allLambdas <- compadre_sub %>%
  cdb_metadata() %>%
  select(SpeciesAccepted, lambda)
```

We calculate some summary statistics…

```
mean(allLambdas$lambda)

## [1] 1.291234

median(allLambdas$lambda)

## [1] 1.010773
```

Then, we plot the λ data to create Fig. 3B:

```
Fig3B <- ggplot(data = allLambdas, aes(x = lambda)) +
  geom_histogram(binwidth = 0.05, fill = "#25b7bc", boundary = 1) +
  xlab(expression(italic(lambda))) +
  xlim(0, 4) +
  geom_vline(xintercept = 1) +
  theme(
    panel.grid.minor.x = element_blank(),
    panel.grid.major.x = element_blank()
  ) +
  ylab("Number of matrices")

Fig3B
```

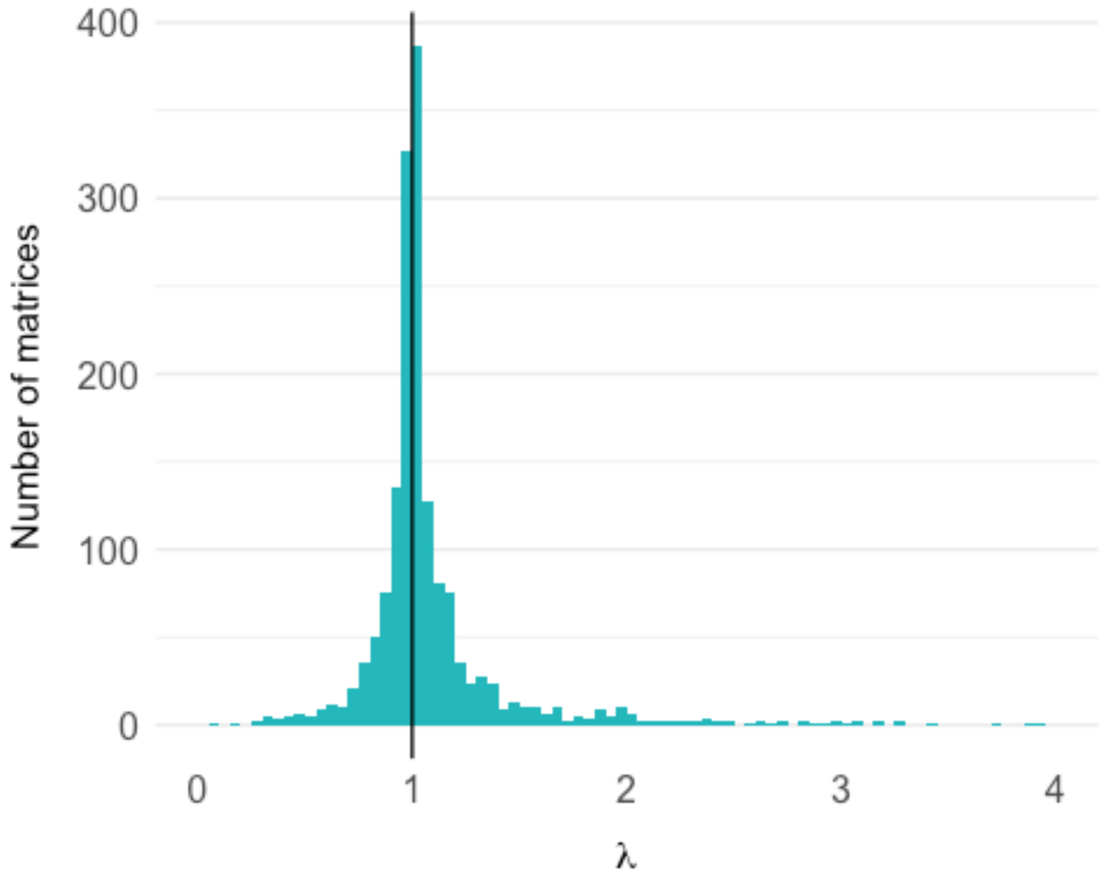


We use a two-sided t-test to check whether λ is significantly different from 1. The results show that the average population is slightly greater than one.

```
t.test(allLambdas$lambda - 1, alternative = "two.sided")

##
##  One Sample t-test
##
## data:  allLambdas$lambda - 1
## t = 2.0245, df = 1658, p-value = 0.04308
## alternative hypothesis: true mean is not equal to 0
## 95 percent confidence interval:
##  0.009079085 0.573389541
## sample estimates:
## mean of x
## 0.2912343
```

##### Regression to the mean

Here, we use the individual matrices for each study year and subset it to those studies that were experimentally unmanipulated, do not lack information on the start and end year of the study and that have a projection interval of 1 (were visited once a year).

```
regToMeanData <- database %>%
  filter(MatrixComposite == "Individual")  %>%
  filter(check_NA_A == FALSE) %>%
  filter(ProjectionInterval == 1) %>%
  filter(MatrixTreatment == "Unmanipulated") %>%
  filter(!is.na(MatrixStartYear)) %>%
  mutate(StudyPopID = as.numeric(as.factor(paste(SpeciesAuthor,
MatrixPopulation)))) %>%
  select(mat, StudyPopID, MatrixPopulation, SpeciesAuthor, SpeciesAccepted,
MatrixStartYear)
```

We again calculate lambda for each matrix.

```
regToMeanData$lambda <- unlist(lapply(matA(regToMeanData), popdemo::eigs,
what = "lambda"))
```

Some lambda calculations result in errors "`More than one eigenvalues have equal absolute magnitude`". In these cases the values calculated are 0. We replace these with NA.

```
regToMeanData <- regToMeanData %>%
  mutate(lambda = ifelse(lambda == 0,NA,lambda))
```

We then calculate a slope for each regression i.e. population. We do this by looping through every unique study-population and fitting an OLS regression. We extract the slope of the regression relationship, and the number of years of data. We filter to only get the slope if there are 5+ years of data.

```
nstudy <- length(unique(regToMeanData$StudyPopID))
regToMean_output <- data.frame(slope = rep(NA, nstudy), nyr = NA)

for (i in 1:nstudy) {
  tempData <- regToMeanData %>% filter(StudyPopID == i)
  if (nrow(tempData) > 4) {
    regToMean_output$slope[i] <- coef(lm(lambda ~ MatrixStartYear, data =
tempData))[2]
  } else {
    regToMean_output$slope[i] <- NA
  }
  regToMean_output$nyr[i] <- nrow(tempData)
}

regToMean_output <- regToMean_output %>%
  filter(nyr >=5)
```

We then use a t-test to ask whether the slope is significantly different to 0. It is not.

```
# T-test on whether th slope is different from 0
t.test(regToMean_output %>%
  pull(slope),alternative = "two.sided")
```

```
##
##  One Sample t-test
##
## data:  regToMean_output %>% pull(slope)
## t = -0.019584, df = 192, p-value = 0.9844
## alternative hypothesis: true mean is not equal to 0
## 95 percent confidence interval:
##  -0.07810938  0.07657351
## sample estimates:
##     mean of x
## -0.0007679328
```

We plot the data to create Fig. 3C:

```
Fig3C <- ggplot(regToMean_output, aes(x = slope)) +
  geom_density(fill = "#D3F0F1", colour = "#25b7bc") +
  geom_vline(xintercept = 0) +
  theme(
    panel.grid.minor.x = element_blank(),
    panel.grid.major.x = element_blank()
  ) +
  ylab("Density") +
  xlab("Slope") +
  NULL

Fig3C
```

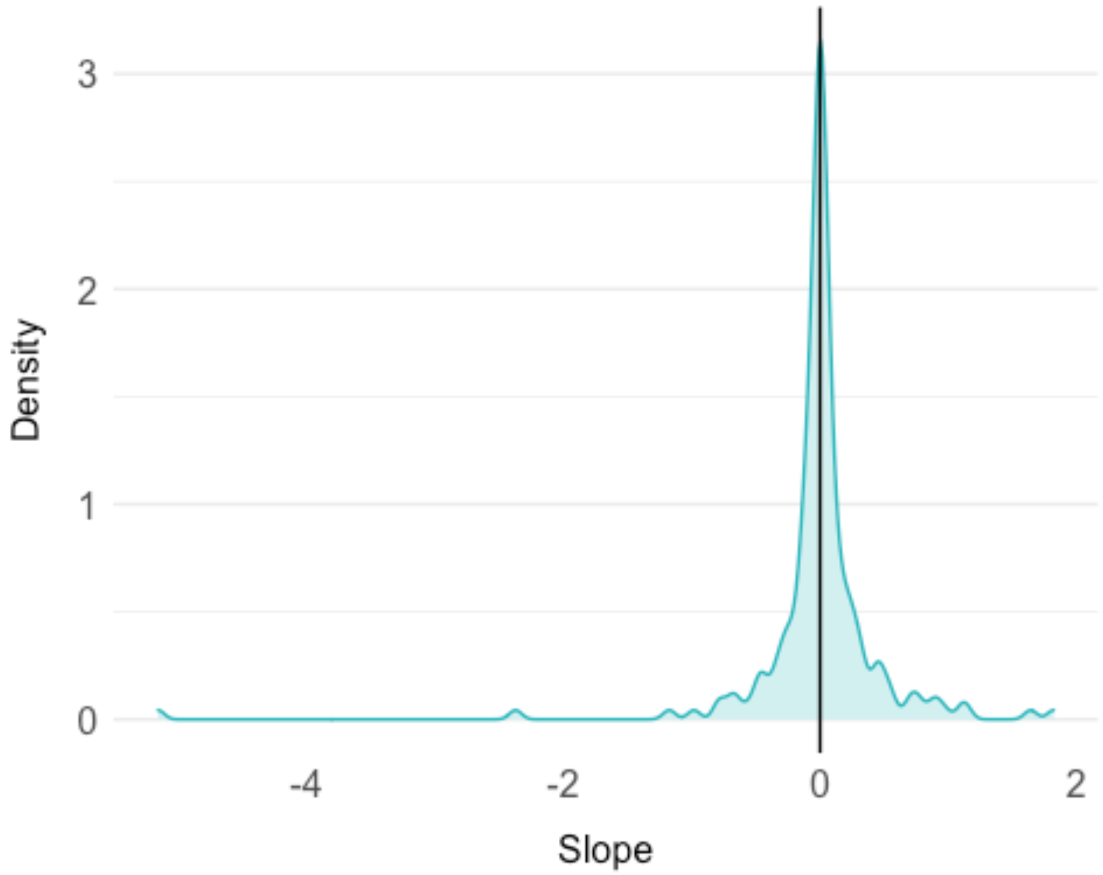


#### E) HOW ARE THE MPMS CONSTRUCTED?

This section focuses on biases in HOW we study populations. This includes biases in:

- the temporal replication for species
- the spatial replication for species
- the matrix dimension used for each species
- the prevalence of averaging over MPM elements

##### Temporal replication

```
temporalRepData <- db %>%
  select(SpeciesAuthor, Study, StudyDuration) %>%
  mutate(StudyDuration = as.numeric(StudyDuration)) %>%
  unique()

(m <- mean(temporalRepData$StudyDuration, na.rm = TRUE))

## [1] 5.48441

(n <- median(temporalRepData$StudyDuration, na.rm = TRUE))

## [1] 4
```

```
summary(temporalRepData$StudyDuration)

##    Min. 1st Qu.  Median    Mean 3rd Qu.    Max.    NA's 
##   1.000   3.000   4.000   5.484   6.000  51.000      47
```

We summarise the data to make a histogram.

```
temporalRepData_summary <- temporalRepData %>%
  group_by(StudyDuration) %>%
  summarise(n = length(SpeciesAuthor))

plot_studyDurationDistrib <- ggplot(data = temporalRepData_summary, aes(x = 
StudyDuration, y = n)) +
  geom_bar(stat = "identity", size = 2, fill = "#25b7bc") +
  geom_vline(xintercept = m, colour = "#bc25b7", size = 1) +
  geom_vline(xintercept = n, colour = "#b7bc25", size = 1) +
  xlab("Number of study years") +
  ylab("Number of studies")

plot_studyDurationDistrib
```

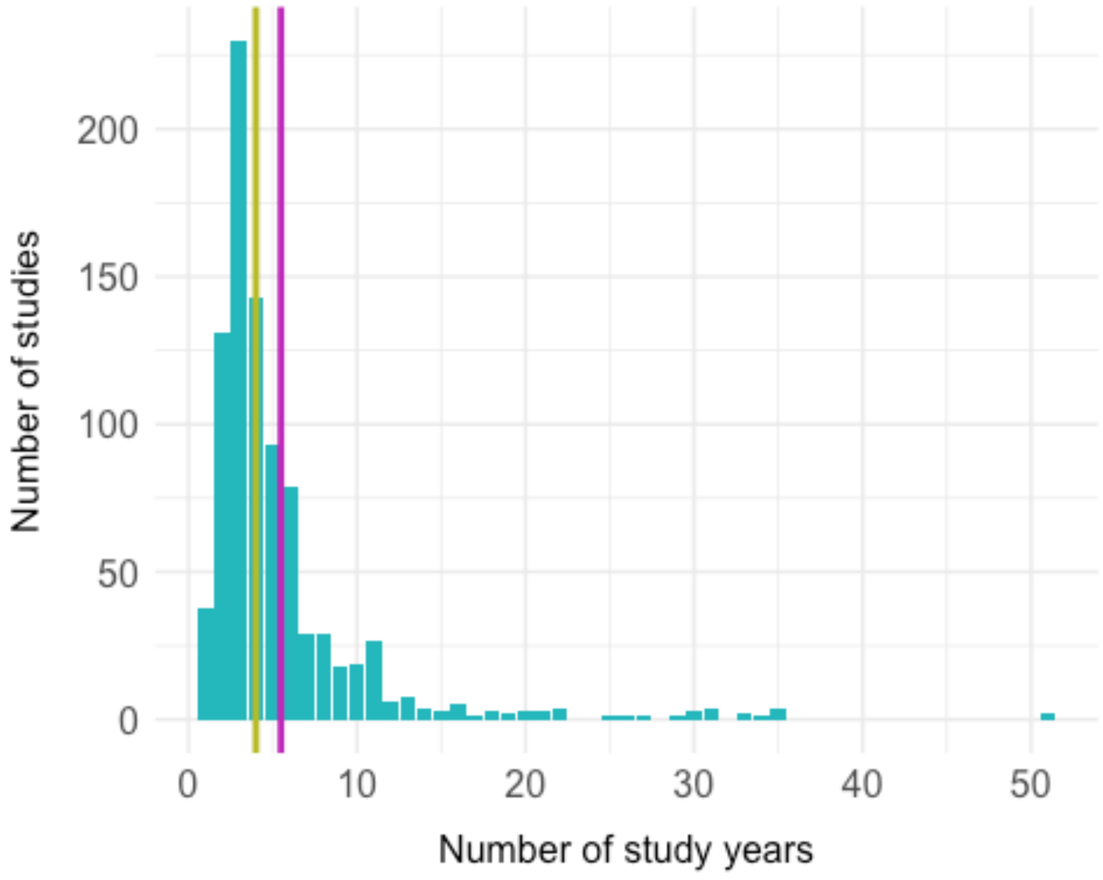


#### Spatial replication

```
spatialRepData <- db %>%
  select(SpeciesAuthor, Study, NumberPopulations) %>%
```

```r
  mutate(NumberPopulations = as.numeric(NumberPopulations)) %>%
  unique()

(m <- mean(spatialRepData$NumberPopulations, na.rm = TRUE))

## [1] 2.818589

(n <- median(spatialRepData$NumberPopulations, na.rm = TRUE))

## [1] 2

summary(spatialRepData$NumberPopulations)

##    Min. 1st Qu.  Median    Mean 3rd Qu.    Max.    NA's
##   1.000   1.000   2.000   2.819   3.000  60.000      51
```

We summarise the data to plot in a histogram.

```r
spatialRepData_summary <- spatialRepData %>%
  group_by(NumberPopulations) %>%
  summarise(n = length(SpeciesAuthor))

plot_spatialRepDistrib <- ggplot(data = spatialRepData_summary, aes(x =
NumberPopulations, y = n)) +
  geom_bar(stat = "identity", size = 2, fill = "#25b7bc") +
  geom_vline(xintercept = m, colour = "#bc25b7", size = 1) +
  geom_vline(xintercept = n, colour = "#b7bc25", size = 1) +
  scale_x_continuous(breaks = c(seq(0, 55,5), 60, 70, 80)) +
  xlab("Number of sites") +
  ylab("Number of studies")

plot_spatialRepDistrib
```

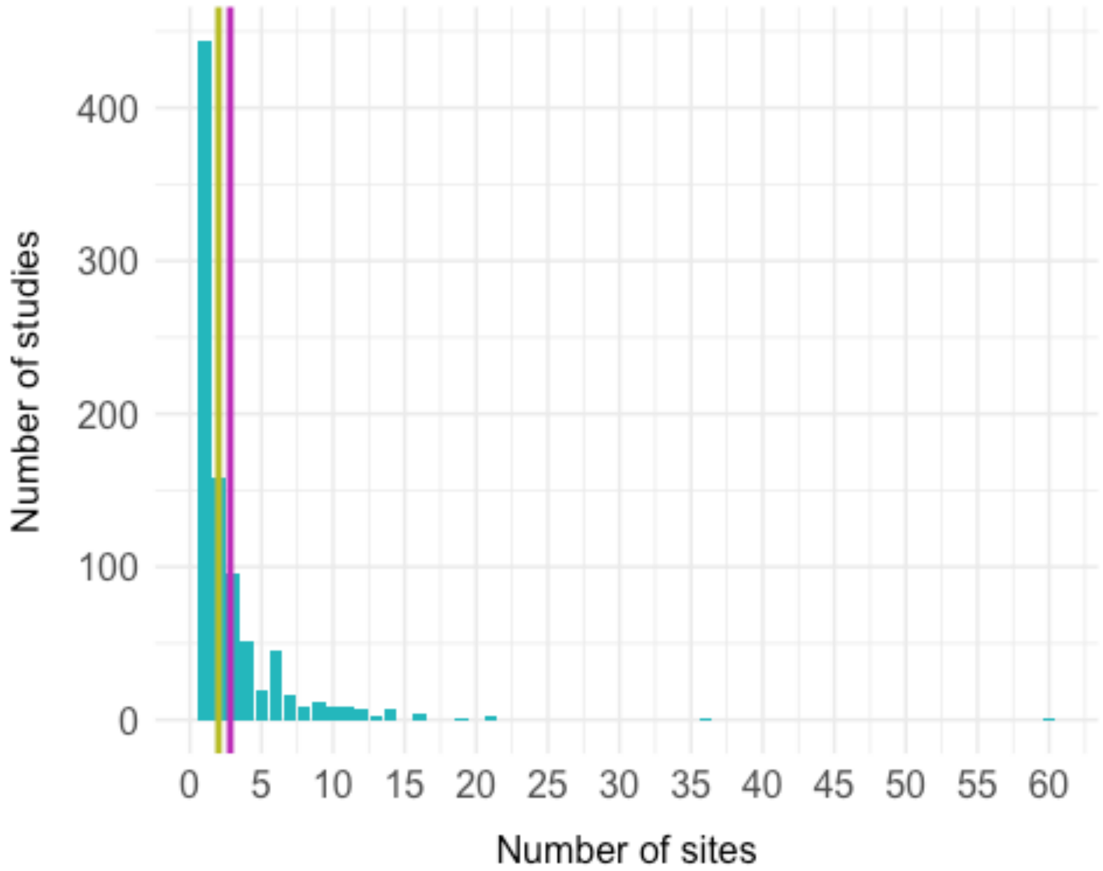


#### Matrix dimension

```
# matDim per individual Study
matrixDimData <- db %>%
  select(SpeciesAuthor, Study, MatrixDimension) %>%
  unique()

(m <- mean(matrixDimData$MatrixDimension, na.rm = TRUE))

## [1] 6.507903

(n <- median(matrixDimData$MatrixDimension, na.rm = TRUE))

## [1] 5

summary(matrixDimData$MatrixDimension)

##    Min. 1st Qu.  Median    Mean 3rd Qu.    Max.
##   1.000   4.000   5.000   6.508   7.000  60.000

matrixDimData_summary <- matrixDimData %>%
  group_by(MatrixDimension) %>%
  summarize(n = length(unique(SpeciesAuthor)))
```

```
plot_matrixDimDistrib <- ggplot(data = matrixDimData_summary, aes(x =
MatrixDimension, y = n)) +
  geom_bar(stat = "identity", size = 2, fill = "#25b7bc") +
  geom_vline(xintercept = m, colour = "#bc25b7", size = 1) +
  geom_vline(xintercept = n, colour = "#b7bc25", size = 1) +
  scale_x_continuous(breaks = c(seq(0, 55,5), 60, 70, 80)) +
  xlab("Matrix dimension") +
  ylab("Number of matrices")

plot_matrixDimDistrib
```

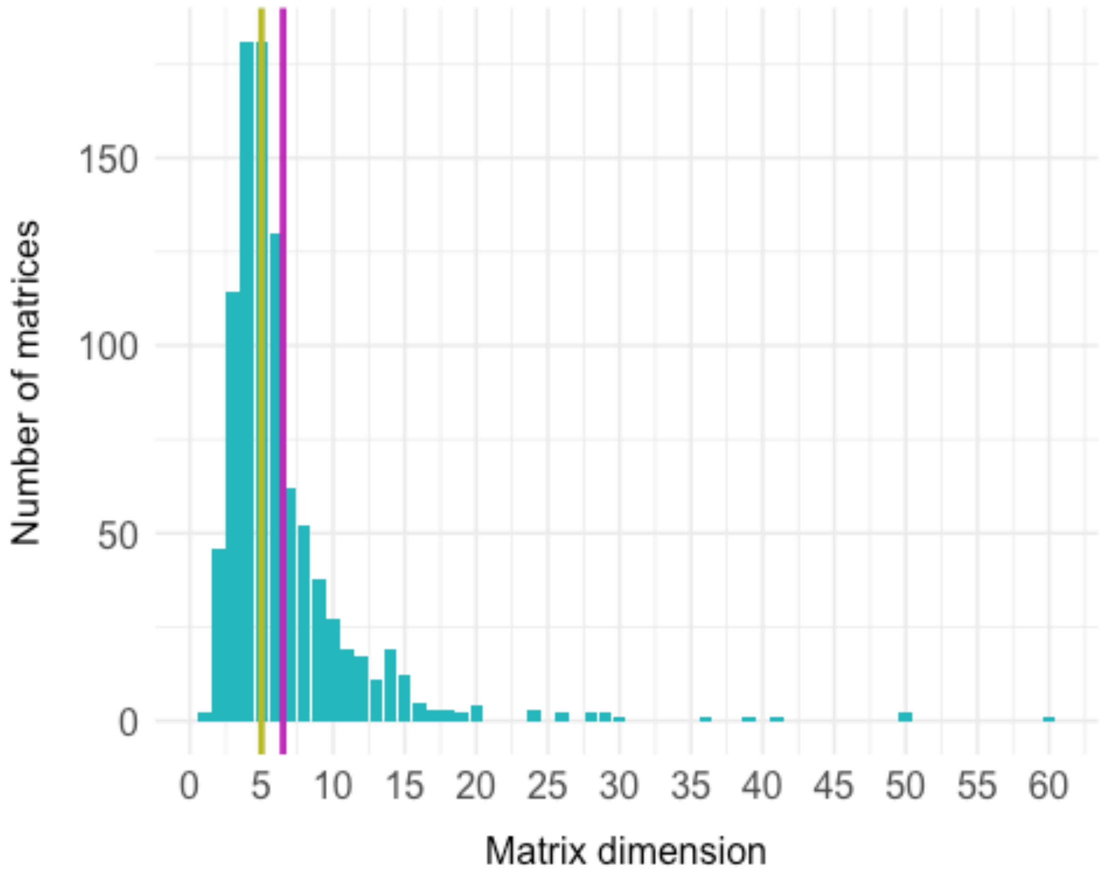


#### Averaging across MPM elements

```
DB <- database
```

##### Survival

Here we work out whether there appears to be averaging of survival elements in the matrices. We want to work with U matrices that don’t contain NAs and do contain nonzero entries we can calculate stuff from these. `DB$check_nonzero_U` is a variable that is `TRUE` if `matU` contains at least one nonzero number and no `NA`s, `FALSE` otherwise.

```
DB$check_nonzero_U <- !DB$check_NA_U & !DB$check_zero_U
```

Note that these variables are connected to the "`MatrixSplit`" variable, as a U matrix that indivisible contains all `NA`s. However a matrix can be '`Divided`' and contain either `NA`s, or no nonzero entries:

We can check this here as follows (results not shown).

```
which(DB$check_NA_U + (DB$MatrixSplit == "Divided") == 2)
which(DB$check_zero_U + (DB$MatrixSplit == "Divided") == 2)
```

A matrix can also be recorded as `indivisible` but have a nonzero U matrix. We can check that here (results not shown).

```
which(DB$check_nonzero_U + (DB$MatrixSplit == "Indivisible") == 2)
```

We calculate survival per stage (column sums of U matrix). `stageSurv` is a list containing the column sums of `matU` (survival per stage).

```
stageSurv <- lapply(matU(DB), colSums)
```

We then find the groups of consecutive average survival using run length encoding (`rle`).

```
survGroups_rle <- lapply(
  stageSurv,
  function(sS) {
    if (!is.null(sS)) rle(sS)
  }
)
```

Lengths only, without corresponding values.

```
survGroupsSummary <- lapply(survGroups_rle, function(G) {
  grpSum <- G$lengths
  names(grpSum) <- NULL
  grpSum
})
```

Next we find the different 'groups' of averages the stages belong to. `survGroups` assigns groups to `stageSurv`. For example, a matrix with `stageSurv` of `c(0.1, 0.12, 0.5, 0.5, 0.5, 0.9)` would have `survGroups` of `c(1, 2, 3, 3, 3, 4)`.

```
survGroups <- lapply(
  survGroupsSummary,
  function(S) {
    unlist(lapply(seq_along(S), function(G) {
      rep(G, S[[G]])
    }))
  }
)
```

Next we identify stages with zero fecundity (rather than being "averages" of survival, these have merely not been observed), and flag in the `survGroups` variable. For example, a matrix with stageSurv of `c(0.1, 0.12, 0.5, 0.5, 0.5, 0, 0, 0.9)` would have survGroupsZero of `c(1, 2, 3, 3, 3, 0, 0, 5)`.

```
survGroupsZero <- mapply(function(sG, sS, mUz) {
  if (is.null(sG) | mUz) sGZ <- NULL
  if (!(is.null(sG)) & !mUz) {
    sGZ <- sG
    sGZ[sS %in% 0] <- 0
  }
  sGZ
},
sG = survGroups, sS = stageSurv,
mUz = DB$check_zero_U
)
```

Nest we collapse these so that zero survival groups aren't included. For example, a matrix with stageSurv of `c(0.1, 0.12, 0.5, 0.5, 0.5, 0, 0, 0.9)` would have survGroupsNoZero of `c(1, 2, 3, 3, 3, 5)`.

```
survGroupsNoZero <- lapply(survGroupsZero, function(sGZ) sGZ[!(sGZ %in% 0)])
```

Run length encoding on new no-zero groups

```
survGroupsNoZero_rle <- lapply(
  survGroupsNoZero,
  function(sGNZ) {
    if (!is.null(sGNZ)) rle(sGNZ)
  }
)
```

Lengths only, without corresponding values

```
survGroupsNoZeroSummary <- lapply(survGroupsNoZero_rle, function(G) {
  grpSum <- G$lengths
  names(grpSum) <- NULL
  grpSum
})
```

Find the maximum number of consecutive averaged stages in each matrix including zeroes:

```
maxConsecSurv <- sapply(
  survGroupsSummary,
  function(sGS) {
    if (!is.null(sGS)) mCS <- max(sGS)
    if (is.null(sGS)) mCS <- NA
    mCS
  }
)
```

Excluding zeroes:

```r
maxConsecSurvNoZero <- sapply(
  survGroupsNoZeroSummary,
  function(sGNZS) {
    if (!is.null(sGNZS)) mCSNZ <- max(sGNZS)
    if (is.null(sGNZS)) mCSNZ <- NA
    mCSNZ
  }
)
```

Extract number of stages.

```r
nStages <- DB$MatrixDimension
```

Now we take the lists of consecutive survival and work out a "traffic light" system to categorise them. Find the length of the consecutive averages for each matrix, then find the maximum for each matrix.

- G If a all stages have different survival, the matrix is rated "G" (green).
- Y If there is apparent averaging (2 or more consecutive survival values the same), but the number of stages with the same value of survival doesn't exceed half the number of stages, then the matrix is rated "Y" (yellow).
- R If half or more of the fecund stages have the same average value, then the matrix is rated "R" (red), unless there is only one stage.

Including Zeroes:

```r
survRYG <- character(length(maxConsecSurv))
survRYG[maxConsecSurv %in% 1] <- "G"
survRYG[maxConsecSurv >= 2 & maxConsecSurv <= nStages / 2] <- "Y"
survRYG[maxConsecSurv >= 2 & maxConsecSurv > nStages / 2] <- "R"
# excluding zeroes:
survRYGNoZero <- character(length(maxConsecSurvNoZero))
survRYGNoZero[maxConsecSurvNoZero %in% 1] <- "G"
survRYGNoZero[maxConsecSurvNoZero >= 2 & maxConsecSurvNoZero <= nStages / 2]
<- "Y"
survRYGNoZero[maxConsecSurvNoZero >= 2 & maxConsecSurvNoZero > nStages / 2]
<- "R"
```

Make very sure `NA`s in the correct places

```r
stageSurv[!DB$check_nonzero_U] <- NA
survGroups_rle[!DB$check_nonzero_U] <- NA
survGroupsSummary[!DB$check_nonzero_U] <- NA
survGroups[!DB$check_nonzero_U] <- NA
survGroupsZero[!DB$check_nonzero_U] <- NA
survGroupsNoZero[!DB$check_nonzero_U] <- NA
survGroupsNoZero_rle[!DB$check_nonzero_U] <- NA
survGroupsNoZeroSummary[!DB$check_nonzero_U] <- NA
maxConsecSurv[!DB$check_nonzero_U] <- NA
maxConsecSurvNoZero[!DB$check_nonzero_U] <- NA
```

```
survRYG[!DB$check_nonzero_U] <- NA
survRYGNoZero[!DB$check_nonzero_U] <- NA
```

Add to metadata

```
DB$UcolSum <- stageSurv
DB$AvSurvGrp <- survGroupsZero
DB$AvSurvMax <- maxConsecSurv
DB$AvSurvMaxNoZero <- maxConsecSurvNoZero
DB$AvSurvRYG <- survRYG
DB$AvSurvRYGNoZero <- survRYGNoZero

AvSurv <- fct_count(DB$AvSurvRYGNoZero)

AvSurv <- AvSurv %>%
  rename("category" = "f") %>%
  as.data.frame()


AvSurv$f <- c("none", ">50%", "\U2265 50%", NA)

AvSurv$f <- factor(AvSurv$f, c("none", "\U2265 50%", ">50%", NA))

AvSurv["Percentage"] <- round((AvSurv$n / (sum(AvSurv$n)) * 100), 1)

PlotN <- ggplot(data = AvSurv, aes(x = f, y = n)) +
  geom_bar(stat = "identity", size = 2, fill = c("#25b7bc", "#b7bc25",
"#bc25b7", "black")) +
  theme(
    panel.grid.minor.x = element_blank(),
    panel.grid.major.x = element_blank(),
    axis.text.x = element_text(size = 10)
  ) +
  scale_y_continuous(limits = c(0, 8500)) +
  xlab("") +
  ylab("Number of matrices")

PlotN
```

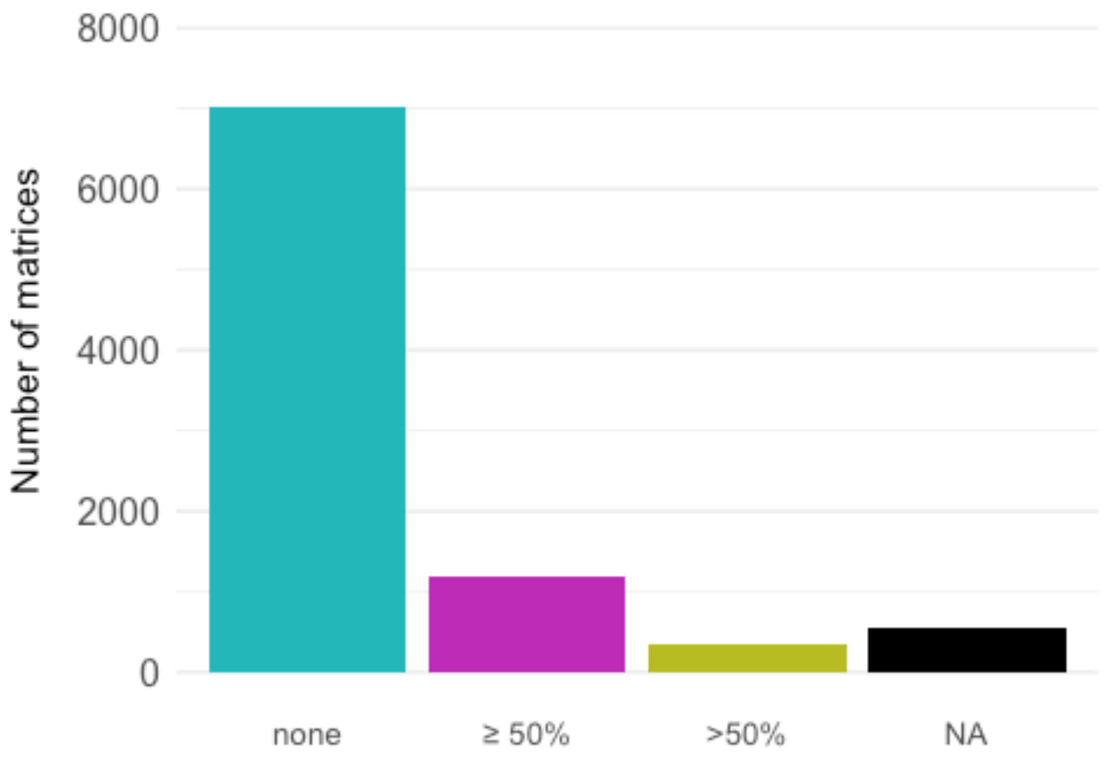


##### Fecundity

Work out whether there appears to be averaging of fecundity We want to work with `F` matrices that don't contain `NA`s and do contain nonzero entries:

we can calculate stuff from these. `DB$check_nonzero_F` is a variable that is `TRUE` if `matU` contains at least one nonzero number and no `NA`s, `FALSE` otherwise.

```
DB$check_zero_F <- sapply(matF(DB), function(M) {
  all(M %in% 0)
})
DB$check_nonzero_F <- !DB$check_NA_F & !DB$check_zero_F
```

Note that these variables are connected to the "`MatrixSplit`" variable, because an `F` matrix that indivisible contains all `NA`s. However a matrix can be '`Divided`' and contain either `NA`s, or no nonzero entries.

We can check these issues (results not shown).

```
which(DB$check_NA_F + (DB$MatrixSplit == "Divided") == 2)
which(DB$check_zero_F + (DB$MatrixSplit == "Divided") == 2)
```

A matrix can also be recorded as indivisible but have a nonzero `F` matrix (results not shown).

```
which(DB$check_nonzero_F + (DB$MatrixSplit == "Indivisible") == 2)
```

Fecundity per stage (column sums of F matrix). `stageFec` is a list containing the column sums of `matF` (the total sexual reproduction per stage)

```
stageFec <- lapply(matF(DB), colSums)
```

First fecund stage. `firstFec` is a variable with the number of the first stage where `stageFec` is greater than zero.

```
firstFec <- mapply(function(sF, Fna, Fz, Fnz) {
  if (Fna | Fz) {
    return(NA)
  }
  if (Fnz) {
    return(min(which(!(sF %in% 0))))
  }
},
sF = stageFec, Fna = DB$check_NA_F,
Fz = DB$check_zero_F, Fnz = DB$check_nonzero_F
)
```

Extract only fecund stages (those to RHS of first fecund stage). `stageFecMature` is a list containing `stageFec` for only those stages after and including `firstFec`.

```
stageFecMature <- mapply(function(sF, fF) {
  if (!is.na(fF)) sF[fF:length(sF)]
},
sF = stageFec, fF = firstFec
)
```

Find the groups of consecutive average fecundity using run length encoding (`rle`).

```
fecGroups_rle <- lapply(
  stageFecMature,
  function(sF) {
    if (!is.null(sF)) rle(sF)
  }
)
```

Lengths only, without corresponding values.

```
fecGroupsSummary <- lapply(fecGroups_rle, function(G) {
  grpSum <- G$lengths
  names(grpSum) <- NULL
  grpSum
})
```

Find the different 'groups' of averages the stages belong to. `fecGroups` assigns groups to `stageFecMature`. For example, a matrix with `stageFecMature` of `c(0.1, 0.12, 0.5, 0.5, 0.5, 0, 0, 0.9)` would have `fecGroups` of `c(1, 2, 3, 3, 3, 4, 4, 5)`.

```
fecGroups <- lapply(
  fecGroupsSummary,
  function(S) {
    unlist(lapply(seq_along(S), function(G) {
      rep(G, S[[G]])
    }))
  }
)
```

Identify stages with zero fecundity (rather than being "averages" of fecundity, these have merely not been observed), and flag in the `fecGroups` variable. For example, a matrix with `stageFecMature` of `c(0.1, 0.12, 0.5, 0.5, 0.5, 0, 0, 0.9)` would have `fecGroupsZero` of `c(1, 2, 3, 3, 3, 0, 0, 5)`.

```
fecGroupsZero <- mapply(function(fG, sFM, mFz) {
  if (is.null(fG) | mFz) fGZ <- NULL
  if (!(is.null(fG)) & !mFz) {
    fGZ <- fG
    fGZ[sFM %in% 0] <- 0
  }
  fGZ
},
fG = fecGroups, sFM = stageFecMature,
mFz = DB$check_zero_F
)
```

Collapse these so that zero fecundity groups aren't included. For example, a matrix with `stageFecMature` of `c(0.1, 0.12, 0.5, 0.5, 0.5, 0, 0, 0.9)` would have `fecGroupsNoZero` of `c(1, 2, 3, 3, 3, 5)`.

```
fecGroupsNoZero <- lapply(fecGroupsZero, function(fGZ) fGZ[!(fGZ %in% 0)])
# run length encoding on new no-zero groups
fecGroupsNoZero_rle <- lapply(
  fecGroupsNoZero,
  function(fGNZ) {
    if (!is.null(fGNZ)) rle(fGNZ)
  }
)
```

Lengths only, without corresponding values.

```
fecGroupsNoZeroSummary <- lapply(fecGroupsNoZero_rle, function(G) {
  grpSum <- G$lengths
  names(grpSum) <- NULL
  grpSum
})
```

Find the maximum number of consecutive averaged stages in each matrix including zeroes:

```
maxConsecFec <- sapply(
  fecGroupsSummary,
```

```
  function(fGS) {
    if (!is.null(fGS)) mCF <- max(fGS)
    if (is.null(fGS)) mCF <- NA
    mCF
  }
)
```

Excluding zeroes:

```
maxConsecFecNoZero <- sapply(
  fecGroupsNoZeroSummary,
  function(fGNZS) {
    if (!is.null(fGNZS)) mCFNZ <- max(fGNZS)
    if (is.null(fGNZS)) mCFNZ <- NA
    mCFNZ
  }
)
```

Extract number of fertile stages

```
nFstages <- DB$MatrixDimension - firstFec + 1
```

Extract number of reproducing stages

```
nRstages <- sapply(stageFec, function(sF) length(which(sF != 0)))
```

Take the lists of consecutive fecundity and work out a “traffic light” system to categorise them. Find the length of the consecutive averages for each matrix, then find the maximum for each matrix.

- G If a all stages have different fertility, the matrix is rated “G” (green).
- Y If there is apparent averaging (2 or more consecutive fertility values the same), but the number of stages with the same value of fertility doesn’t exceed half the number of fecund stages, then the matrix is rated “Y” (yellow).
- R If half or more of the fecund stages have the same average value, then the matrix is rated “R” (red), unless there is only one fecund stage.

(NOTE: these could be changed to use `nRstages` instead of `nFstages`)

Including zeroes:

```
fecRYG <- character(length(maxConsecFec))
fecRYG[maxConsecFec %in% 1] <- "G"
fecRYG[maxConsecFec >= 2 & maxConsecFec <= nFstages / 2] <- "Y"
fecRYG[maxConsecFec >= 2 & maxConsecFec > nFstages / 2] <- "R"
# excluding zeroes:
fecRYGNoZero <- character(length(maxConsecFecNoZero))
fecRYGNoZero[maxConsecFecNoZero %in% 1] <- "G"
fecRYGNoZero[maxConsecFecNoZero >= 2 & maxConsecFecNoZero <= nFstages / 2] <-
"Y"
```

```
fecRYGNoZero[maxConsecFecNoZero >= 2 & maxConsecFecNoZero > nFstages / 2] <-
"R"
```

Make very sure NAs in the correct places

```
stageFec[!DB$check_nonzero_F] <- NA
firstFec[!DB$check_nonzero_F] <- NA
stageFecMature[!DB$check_nonzero_F] <- NA
fecGroups_rle[!DB$check_nonzero_F] <- NA
fecGroupsSummary[!DB$check_nonzero_F] <- NA
fecGroups[!DB$check_nonzero_F] <- NA
fecGroupsZero[!DB$check_nonzero_F] <- NA
fecGroupsNoZero[!DB$check_nonzero_F] <- NA
fecGroupsNoZero_rle[!DB$check_nonzero_F] <- NA
fecGroupsNoZeroSummary[!DB$check_nonzero_F] <- NA
maxConsecFec[!DB$check_nonzero_F] <- NA
maxConsecFecNoZero[!DB$check_nonzero_F] <- NA
nFstages[!DB$check_nonzero_F] <- NA
fecRYG[!DB$check_nonzero_F] <- NA
fecRYGNoZero[!DB$check_nonzero_F] <- NA
```

Add to metadata.

```
DB$FcolSum <- stageFec
DB$FirstF <- firstFec
DB$FnStages <- nFstages
DB$AvFecGrp <- fecGroupsZero
DB$AvFecMaxNoZero <- maxConsecFecNoZero
DB$AvFecRYG <- fecRYG
DB$AvFecRYGNoZero <- fecRYGNoZero

FecSurv <- fct_count(DB$AvFecRYGNoZero)

FecSurv <- FecSurv %>%
  rename("category" = "f") %>%
  as.data.frame()


FecSurv$f <- c("none", ">50%", "\U2265 50%", NA)

FecSurv$f <- factor(FecSurv$f, c("none", "\U2265 50%", ">50%", NA))

FecSurv["Percentage"] <- round((FecSurv$n / (sum(FecSurv$n)) * 100), 1)

PlotO <- ggplot(data = FecSurv, aes(x = f, y = n)) +
    geom_bar(stat = "identity", size = 2, fill = c("#25b7bc", "#b7bc25",
"#bc25b7", "black")) +
  theme(
    panel.grid.minor.x = element_blank(),
    panel.grid.major.x = element_blank(),
    axis.text.x = element_text(size = 10)
  ) +
  scale_y_continuous(limits = c(0, 8500)) +
```

```
  xlab("") +
  ylab("")

PlotO
```

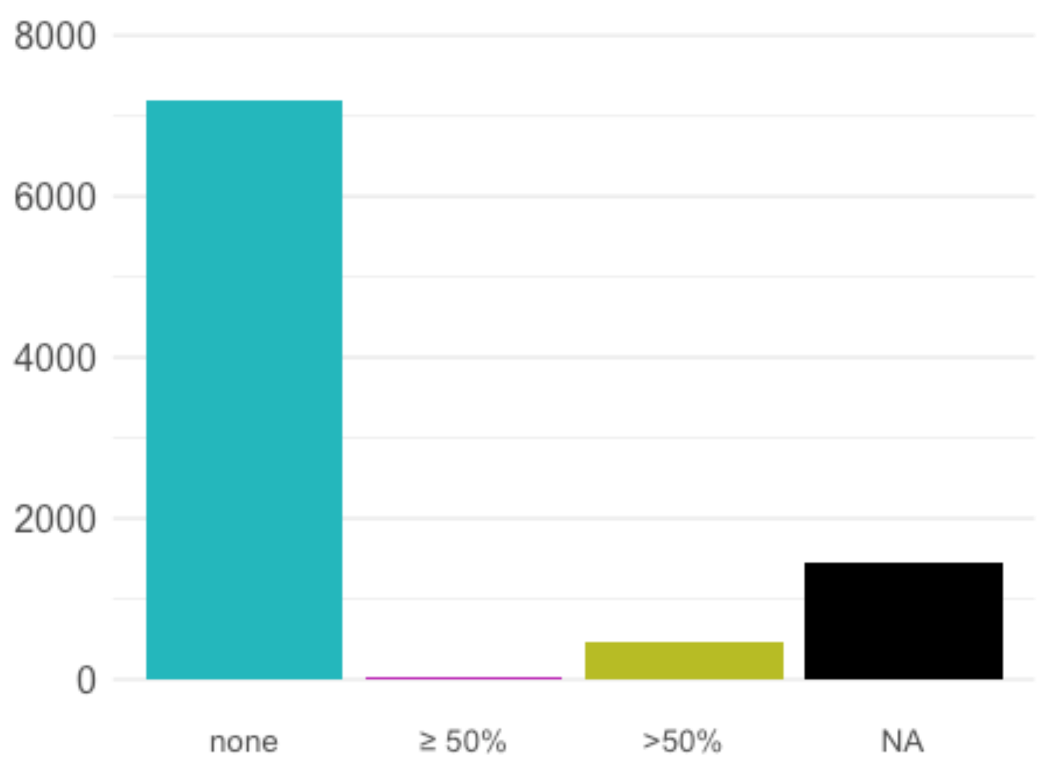


##### Clonality

Work out whether there appears to be averaging of clonality. We want to work with C matrices that don't contain NAs and do contain nonzero entries: we can calculate stuff from these. `DB$check_nonzero_C` is a variable that is `TRUE` if `matC` contains at least one nonzero number and no `NA`s, `FALSE` otherwise.

```
DB$check_zero_C <- sapply(matC(DB), function(M) {
  all(M %in% 0)
})
DB$check_nonzero_C <- !DB$check_NA_C & !DB$check_zero_C
```

Note that these variables are connected to the "`MatrixSplit`" variable, as an `C` matrix that indivisible contains all `NA`s. However a matrix can be '`Divided`' and contain either `NA`s, or no nonzero entries.

Results not shown.

```
which(DB$check_NA_C + (DB$MatrixSplit == "Divided") == 2)
which(DB$check_zero_C + (DB$MatrixSplit == "Divided") == 2)
```

A matrix can also be recorded as indivisible but have a nonzero C matrix:

Results not shown.

```
which(DB$check_nonzero_C + (DB$MatrixSplit == "Indivisible") == 2)
```

Clonality per stage (column sums of `C` matrix). `stageClon` is a list containing the column sums of `matC` (the total sexual reproduction per stage).

```
stageClon <- lapply(matC(DB), colSums)
```

First clonal stage. `firstClon` is a variable with the number of the first stage where `stageClon` is greater than zero.

```
firstClon <- mapply(function(sC, Cna, Cz, Cnz) {
  if (Cna | Cz) {
    return(NA)
  }
  if (Cnz) {
    return(min(which(!(sC %in% 0))))
  }
},
sC = stageClon, Cna = DB$check_NA_C,
Cz = DB$check_zero_C, Cnz = DB$check_nonzero_C
)
```

Extract only clonal stages (those to RHS of first clonal stage). `stageClonMature` is a list containing `stageClon` for only those stages after and including `firstClon`.

```
stageClonMature <- mapply(function(sC, fC) {
  if (!is.na(fC)) sC[fC:length(sC)]
},
sC = stageClon, fC = firstClon
)
```

Find the groups of consecutive average clonality using run length encoding (`rle`)

```
clonGroups_rle <- lapply(
  stageClonMature,
  function(sC) {
    if (!is.null(sC)) rle(sC)
  }
)
```

Lengths only, without corresponding values.

```
clonGroupsSummary <- lapply(clonGroups_rle, function(G) {
  grpSum <- G$lengths
  names(grpSum) <- NULL
```

```
  grpSum
})
```

Find the different 'groups' of averages the stages belong to. `clonGroups` assigns groups to `stageClonMature`. For example, a matrix with `stageClonMature` of `c(0.1, 0.12, 0.5, 0.5, 0.5, 0, 0, 0.9)` would have `clonGroups` of `c(1, 2, 3, 3, 3, 4, 4, 5)`.

```
clonGroups <- lapply(
  clonGroupsSummary,
  function(S) {
    unlist(lapply(seq_along(S), function(G) {
      rep(G, S[[G]])
    }))
  }
)
```

Identify stages with zero clonality (rather than being "averages" of clonality, these have merely not been observed), and flag in the `clonGroups` variable. For example, a matrix with `stageClonMature` of `c(0.1, 0.12, 0.5, 0.5, 0.5, 0, 0, 0.9)` would have `clonGroupsZero` of `c(1, 2, 3, 3, 3, 0, 0, 5)`.

```
clonGroupsZero <- mapply(function(cG, sCM, mCz) {
  if (is.null(cG) | mCz) cGZ <- NULL
  if (!(is.null(cG)) & !mCz) {
    cGZ <- cG
    cGZ[sCM %in% 0] <- 0
  }
  cGZ
},
cG = clonGroups, sCM = stageClonMature,
mCz = DB$check_zero_C
)
```

Collapse these so that zero clonality groups aren't included. For example, a matrix with `stageClonMature` of `c(0.1, 0.12, 0.5, 0.5, 0.5, 0, 0, 0.9)` would have `clonGroupsNoZero` of `c(1, 2, 3, 3, 3, 5)`.

```
clonGroupsNoZero <- lapply(clonGroupsZero, function(cGZ) cGZ[!(cGZ %in% 0)])
# run length encoding on new no-zero groups
clonGroupsNoZero_rle <- lapply(
  clonGroupsNoZero,
  function(cGNZ) {
    if (!is.null(cGNZ)) rle(cGNZ)
  }
)
```

Lengths only, without corresponding values.

```
clonGroupsNoZeroSummary <- lapply(clonGroupsNoZero_rle, function(G) {
  grpSum <- G$lengths
  names(grpSum) <- NULL
```

```
  grpSum
})
```

Find the maximum number of consecutive averaged stages in each matrix including zeroes:

```
maxConsecClon <- sapply(
  clonGroupsSummary,
  function(cGS) {
    if (!is.null(cGS)) mCC <- max(cGS)
    if (is.null(cGS)) mCC <- NA
    mCC
  }
)
```

Excluding zeroes:

```
maxConsecClonNoZero <- sapply(
  clonGroupsNoZeroSummary,
  function(cGNZS) {
    if (!is.null(cGNZS)) mCCNZ <- max(cGNZS)
    if (is.null(cGNZS)) mCCNZ <- NA
    mCCNZ
  }
)
```

Extract number of clonal stages.

```
nCstages <- DB$MatrixDimension - firstClon + 1
```

Extract number of reproducing stages

```
# nRstages <- sapply(stageClon, function(sC) length(which(sC != 0)))
```

Take the lists of consecutive clonality and work out a “traffic light” system to categorise them. Find the length of the consecutive averages for each matrix, then find the maximum for each matrix.

- G If a all stages have different clonality, the matrix is rated “G” (green).
- Y If there is apparent averaging (2 or more consecutive clonality values the same), but the number of stages with the same value of clonality doesn’t exceed half the number of clonal stages, then the matrix is rated “Y” (yellow).
- R If half or more of the clonal stages have the same average value, then the matrix is rated “R” (red), unless there is only one clonal stage.

(NOTE: these could be changed to use `nRstages` instead of `nCstages`)

Including Zeroes:

```
clonRYG <- character(length(maxConsecClon))
clonRYG[maxConsecClon %in% 1] <- "G"
clonRYG[maxConsecClon >= 2 & maxConsecClon <= nCstages / 2] <- "Y"
clonRYG[maxConsecClon >= 2 & maxConsecClon > nCstages / 2] <- "R"
```

Excluding zeroes:

```
clonRYGNoZero <- character(length(maxConsecClonNoZero))
clonRYGNoZero[maxConsecClonNoZero %in% 1] <- "G"
clonRYGNoZero[maxConsecClonNoZero >= 2 & maxConsecClonNoZero <= nCstages / 2]
<- "Y"
clonRYGNoZero[maxConsecClonNoZero >= 2 & maxConsecClonNoZero > nCstages / 2]
<- "R"
```

Make very sure NAs in the correct places

```
stageClon[!DB$check_nonzero_C] <- NA
firstClon[!DB$check_nonzero_C] <- NA
stageClonMature[!DB$check_nonzero_C] <- NA
clonGroups_rle[!DB$check_nonzero_C] <- NA
clonGroupsSummary[!DB$check_nonzero_C] <- NA
clonGroups[!DB$check_nonzero_C] <- NA
clonGroupsZero[!DB$check_nonzero_C] <- NA
clonGroupsNoZero[!DB$check_nonzero_C] <- NA
clonGroupsNoZero_rle[!DB$check_nonzero_C] <- NA
clonGroupsNoZeroSummary[!DB$check_nonzero_C] <- NA
maxConsecClon[!DB$check_nonzero_C] <- NA
maxConsecClonNoZero[!DB$check_nonzero_C] <- NA
nCstages[!DB$check_nonzero_C] <- NA
clonRYG[!DB$check_nonzero_C] <- NA
clonRYGNoZero[!DB$check_nonzero_C] <- NA
```

Add to metadata.

```
DB$CcolSum <- stageClon
DB$FirstC <- firstClon
DB$CnStages <- nCstages

# DB$RnStages <- nRstages
DB$AvClonGrp <- clonGroupsZero
DB$AvClonMaxNoZero <- maxConsecClonNoZero
DB$AvClonRYG <- clonRYG
DB$AvClonRYGNoZero <- clonRYGNoZero

ClonSurv <- fct_count(DB$AvClonRYGNoZero)

ClonSurv <- ClonSurv %>%
  rename("category" = "f") %>%
  as.data.frame()

ClonSurv$f <- c("none", ">50%", "\U2265 50%", NA)

ClonSurv$f <- factor(ClonSurv$f, c("none", "\U2265 50%", ">50%", NA))

ClonSurv["Percentage"] <- round((ClonSurv$n / (sum(ClonSurv$n)) * 100), 1)
```

```
PlotP <- ggplot(data = ClonSurv, aes(x = f, y = n)) +
  geom_bar(stat = "identity", size = 2, fill = c("#25b7bc", "#b7bc25",
"#bc25b7", "black")) +
  theme(
    panel.grid.minor.x = element_blank(),
    panel.grid.major.x = element_blank(),
    axis.text.x = element_text(size = 10)
  ) +
  scale_y_continuous(limits = c(0, 8500)) +
  xlab("") +
  ylab("") +
  NULL

PlotP
```

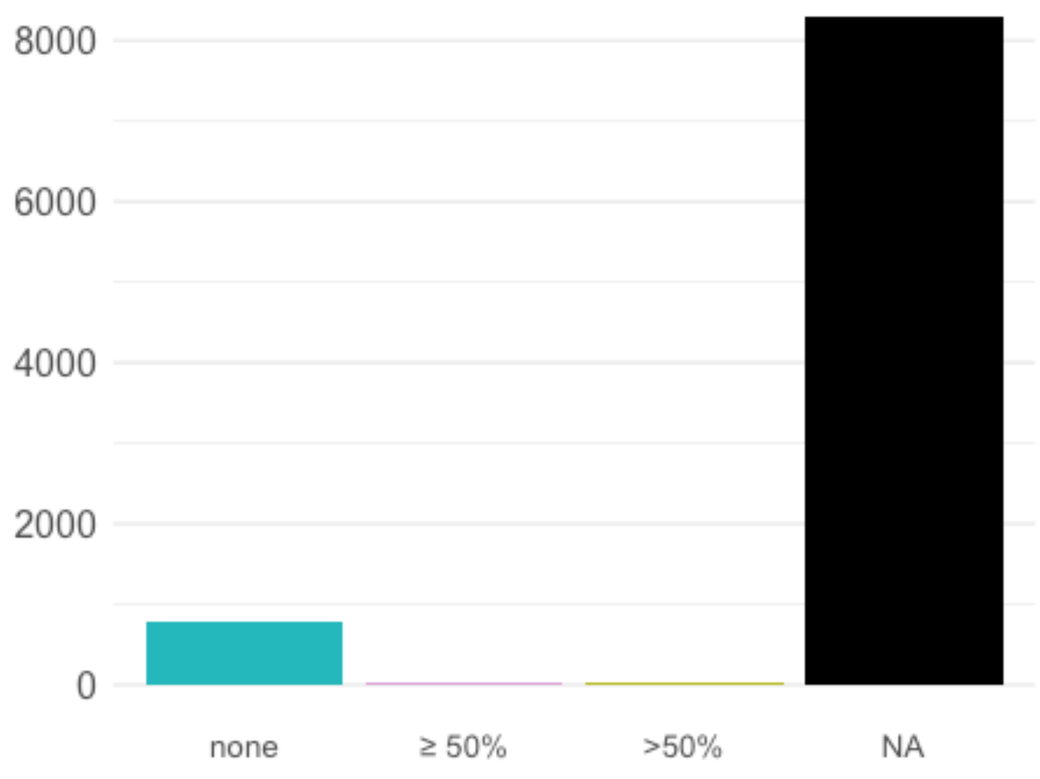


#### F) Appendix

This script contains all the analyses and plot for the appendix of our paper

This includes: - world map of studies in COMPADRE - Raunkiær growth form distribution - violin plots for variation in matrix dimension across growth forms and ecoregions

#### COMPADRE worldmap

```
WorldData <- map_data("world") %>%
  fortify()

unique_locations <- unique(select(db, Lat, Lon))

plot_worldmap <- ggplot() +
  geom_map(
    data = WorldData, map = WorldData,
    aes( map_id = region),
    fill = "grey50", colour = "gray79", size = 0.2
  ) +
  geom_point(
    data = unique_locations,
    aes(x = Lon, y = Lat),
    shape = 21, color = "gray18", fill = "#25b7bc", cex = 1.7
  ) +   coord_map("rectangular", lat0 = 0, xlim = c(-180, 180), ylim = c(-60,
90)) +
  scale_fill_continuous(low = "thistle2", high = "darkred", guide =
"colorbar") +
  scale_y_continuous(breaks = c()) +
  scale_x_continuous(breaks = c()) +
  labs(fill = "legend", title = "", x = "", y = "") +
  NULL

plot_worldmap
```

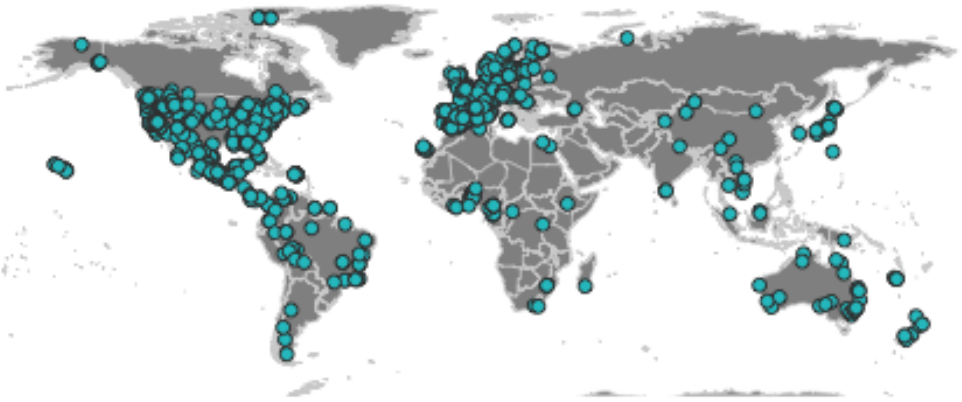

#### Variation across growth forms and ecoregions

First we prepare the data:

```
clean_db_all <- db %>%
  select(Study, SpeciesAuthor, OrganismType, Ecoregion, ecoregionLabel,
StudyDuration, NumberPopulations, MatrixDimension) %>%
  mutate(StudyDuration = ifelse(StudyDuration == "NDY", NA, StudyDuration))
%>%
  mutate(StudyDuration = ifelse(StudyDuration == "FRANK", NA, StudyDuration))
%>%
  mutate(StudyDuration = as.numeric(StudyDuration)) %>%
  mutate(NumberPopulations = ifelse(NumberPopulations == "NDY", NA,
NumberPopulations)) %>%
  mutate(NumberPopulations = as.numeric(NumberPopulations)) %>%
  unique()
```

We also create a new function to create the labels in Tukey test plots:

```
generate_label_df <- function(TUKEY, variable) {

  # Extract labels and factor levels from Tukey post-hoc
  Tukey.levels <- TUKEY[[variable]][, 4]
```

```
  Tukey.labels <- data.frame(multcompLetters(Tukey.levels)["Letters"])

  # Put the labels in the same order as in the boxplot :
  Tukey.labels$treatment <- rownames(Tukey.labels)
  Tukey.labels <- Tukey.labels[order(Tukey.labels$treatment), ]
  return(Tukey.labels)
}
```

#### Growth form

We remove growth forms with very small sample sizes.

```
clean_db <- clean_db_all %>%
  filter(!OrganismType %in% c("Algae", "Bryophyte", "Fern", "Liana",
"Lichen"))
```

##### Temporal replication

```
# average study years per growth form
Temp_growth <- clean_db %>%
  select(OrganismType, Study, SpeciesAuthor, StudyDuration) %>%
  unique()

model <- aov(StudyDuration ~ OrganismType, data = Temp_growth)
out <- HSD.test(model, "OrganismType",
  group = TRUE, console = TRUE,
  main = "temporal replication across growth forms"
)

##
## Study: temporal replication across growth forms
##
## HSD Test for StudyDuration
##
## Mean Square Error:  30.25488
##
## OrganismType,  means
##
##            StudyDuration      std   r Min Max
## Annual          2.868421 1.663130  38   1   9
## Epiphyte        4.836735 3.084551  49   2  14
## Herb            5.900000 5.803098 450   1  35
## Palm            3.958333 2.103273  48   2  12
## Shrub           5.554217 6.055139  83   1  51
## Succulent       4.015873 2.975665  63   2  22
## Tree            6.388489 6.950665 139   1  51
##
## Alpha: 0.05 ; DF Error: 863
## Critical Value of Studentized Range: 4.179435
##
## Groups according to probability of means differences and alpha level( 0.05
```

```
)
##
## Treatments with the same letter are not significantly different.
##
##              StudyDuration groups
## Tree              6.388489      a
## Herb              5.900000      a
## Shrub             5.554217     ab
## Epiphyte          4.836735     ab
## Succulent         4.015873     ab
## Palm              3.958333     ab
## Annual            2.868421      b

plot(out)
```

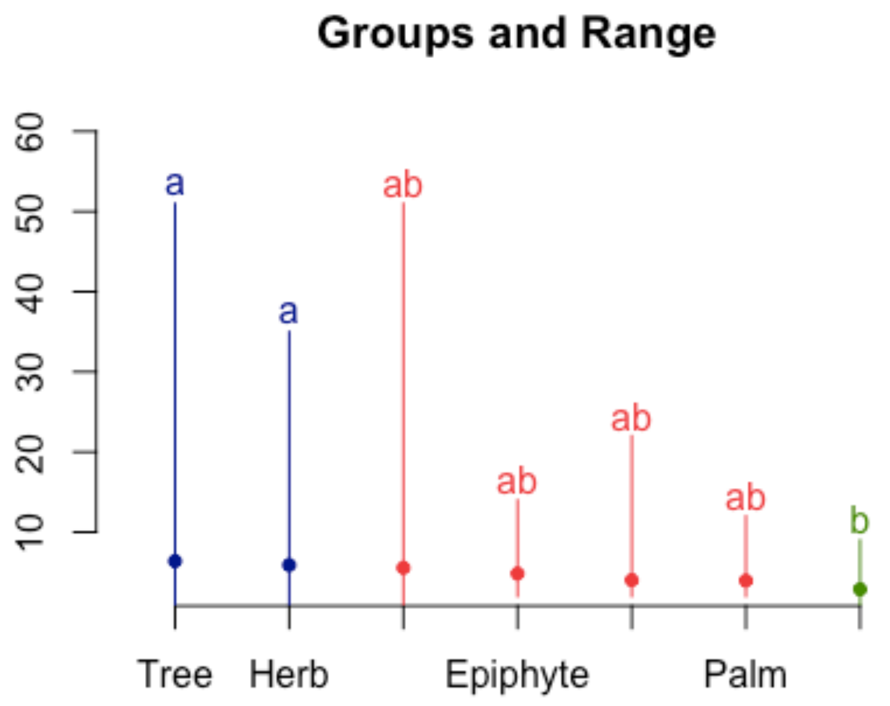


```
Temp_growth <- clean_db %>%
  select(OrganismType, Study, StudyDuration) %>%
  unique() %>%
  group_by(OrganismType)

# adding sample size to the x labels
sampleSize <- Temp_growth %>%
  group_by(OrganismType) %>%
  summarize(n = length(unique(Study)))
```

```
labelsGF <- paste0(sampleSize$OrganismType, "\n(n = ", sampleSize$n, ")")

# add comment that Bryophyte, Liana and Fern were removed due to small sample
size
textbox <- "Bryophyte, Liana and Fern were\nremoved due to small sample size"

# cut off after NumberPopulation = 30 --> !!! this removes 2 data points in
Epiphytes, 1 in Palms and 1 in Trees
Plot_TempG <- ggplot(data = Temp_growth, aes(x = OrganismType, y =
StudyDuration, fill = OrganismType)) +
  geom_violin() +
  scale_y_continuous(limits = c(0, 20), breaks = 1:20) +
  scale_x_discrete(labels = labelsGF) +
  theme_minimal() +
  theme(
    plot.title = element_text(hjust = 0.5),
    legend.position = "none"
  ) +
  ylab("Number of study years per study") +
  NULL

Plot_TempG
```

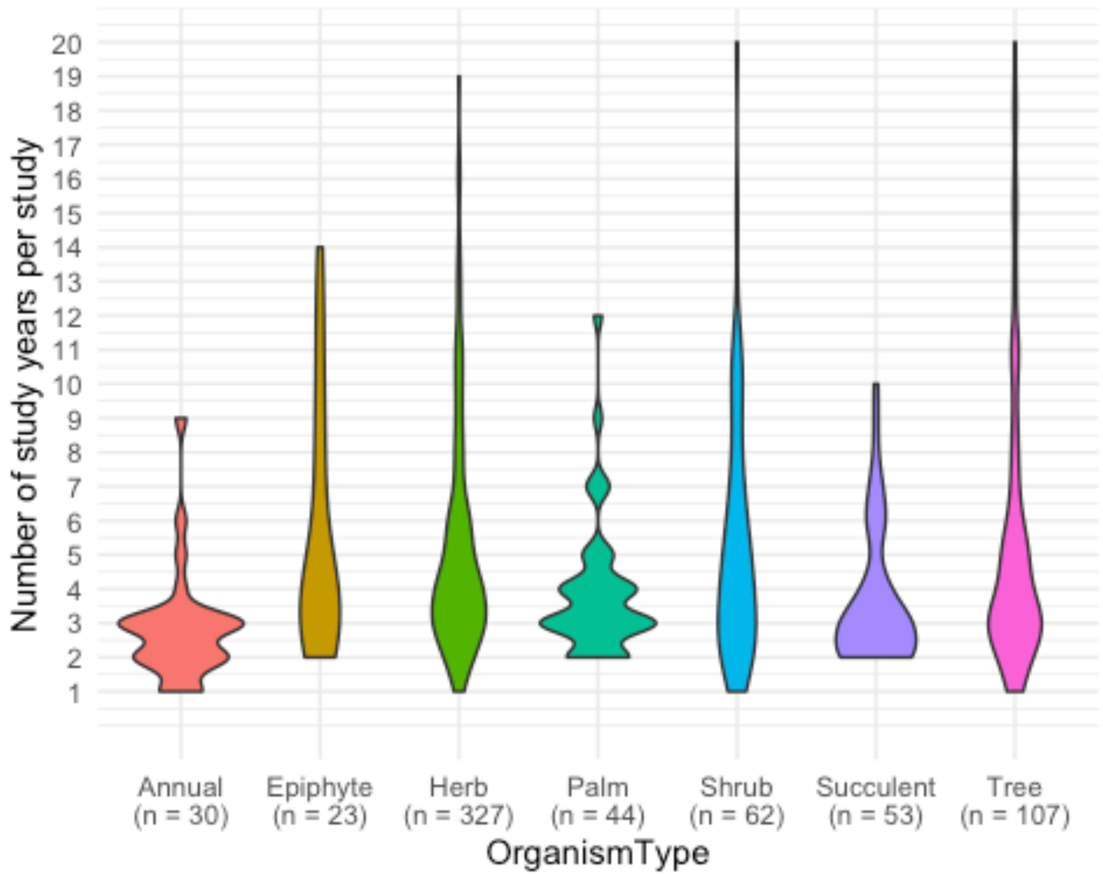

##### Spatial replication

```
# average study years per growth form
Spatial_growth <- clean_db %>%
  select(OrganismType, Study, SpeciesAuthor, NumberPopulations) %>%
  unique()

model <- aov(NumberPopulations ~ OrganismType, data = Spatial_growth)
out <- HSD.test(model, "OrganismType",
  group = TRUE, console = TRUE,
  main = "spatial replication across growth forms"
)

##
## Study: spatial replication across growth forms
##
## HSD Test for NumberPopulations
##
## Mean Square Error:  12.97967
##
## OrganismType,  means
##
##           NumberPopulations      std   r Min Max
## Annual             2.171429 2.093206  35   1  10
## Epiphyte           1.770833 1.812951  48   1   9
## Herb               3.498886 4.554298 449   1  60
## Palm               2.469388 3.221685  49   1  21
## Shrub              2.373494 2.004330  83   1   9
## Succulent          1.887097 1.620614  62   1   9
## Tree               1.914894 2.033608 141   1  12
##
## Alpha: 0.05 ; DF Error: 860
## Critical Value of Studentized Range: 4.17947
##
## Groups according to probability of means differences and alpha level( 0.05
)
##
## Treatments with the same letter are not significantly different.
##
##           NumberPopulations groups
## Herb               3.498886      a
## Palm               2.469388     ab
## Shrub              2.373494     ab
## Annual             2.171429     ab
## Tree               1.914894      b
## Succulent          1.887097      b
## Epiphyte           1.770833      b

plot(out)
```

Groups and Range

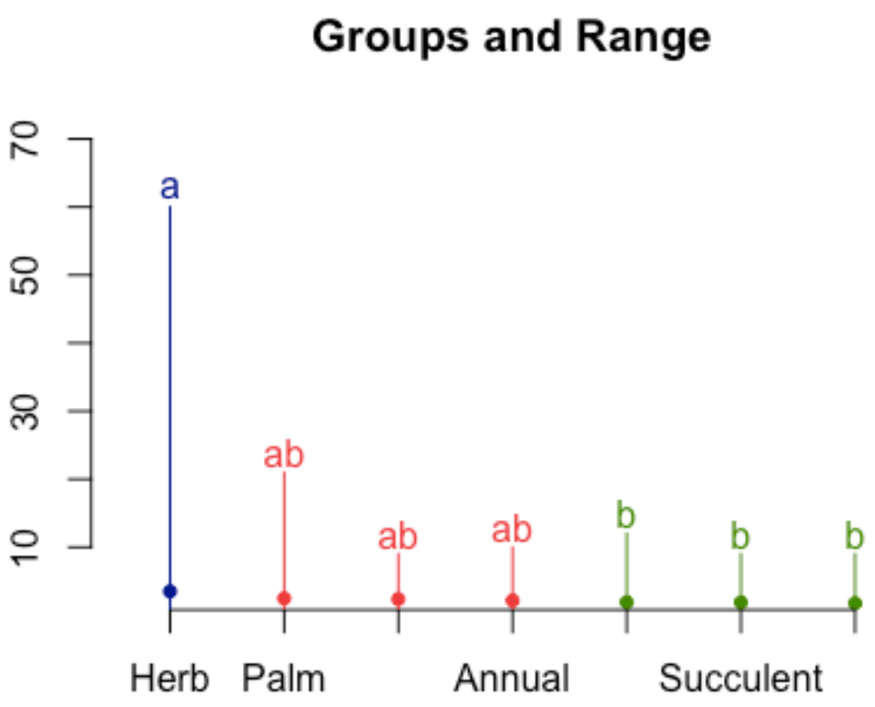


```
Spatial_growth <- clean_db %>%
  select(OrganismType, Study, NumberPopulations) %>%
  unique() %>%
  group_by(OrganismType)

# adding sample size to the x labels
sampleSize <- Spatial_growth %>%
  group_by(OrganismType) %>%
  summarize(n = length(unique(Study)))

labelsGF <- paste0(sampleSize$OrganismType, "\n(n = ", sampleSize$n, ")")

# add comment that Bryophyte, Liana and Fern were removed due to small sample
size
textbox <- "Bryophyte, Liana and Fern were\nremoved due to small sample size"

# cut off after NumberPopulation = 30 --> !!! this removes 2 data points in
Epiphytes, 1 in Palms and 1 in Trees
Plot_SpatialG <- ggplot(
  data = Spatial_growth,
  aes(x = OrganismType, y = NumberPopulations, fill = OrganismType)
) +
  geom_violin() +
```

```
  scale_y_continuous(limits = c(0, 20), breaks = 1:20) +
  scale_x_discrete(labels = labelsGF) +
  theme_minimal() +
  theme(
    plot.title = element_text(hjust = 0.5),
    legend.position = "none"
  ) +
  xlab("OrganismType") +
  ylab("Number of sites per study") +
  NULL

Plot_SpatialG
```

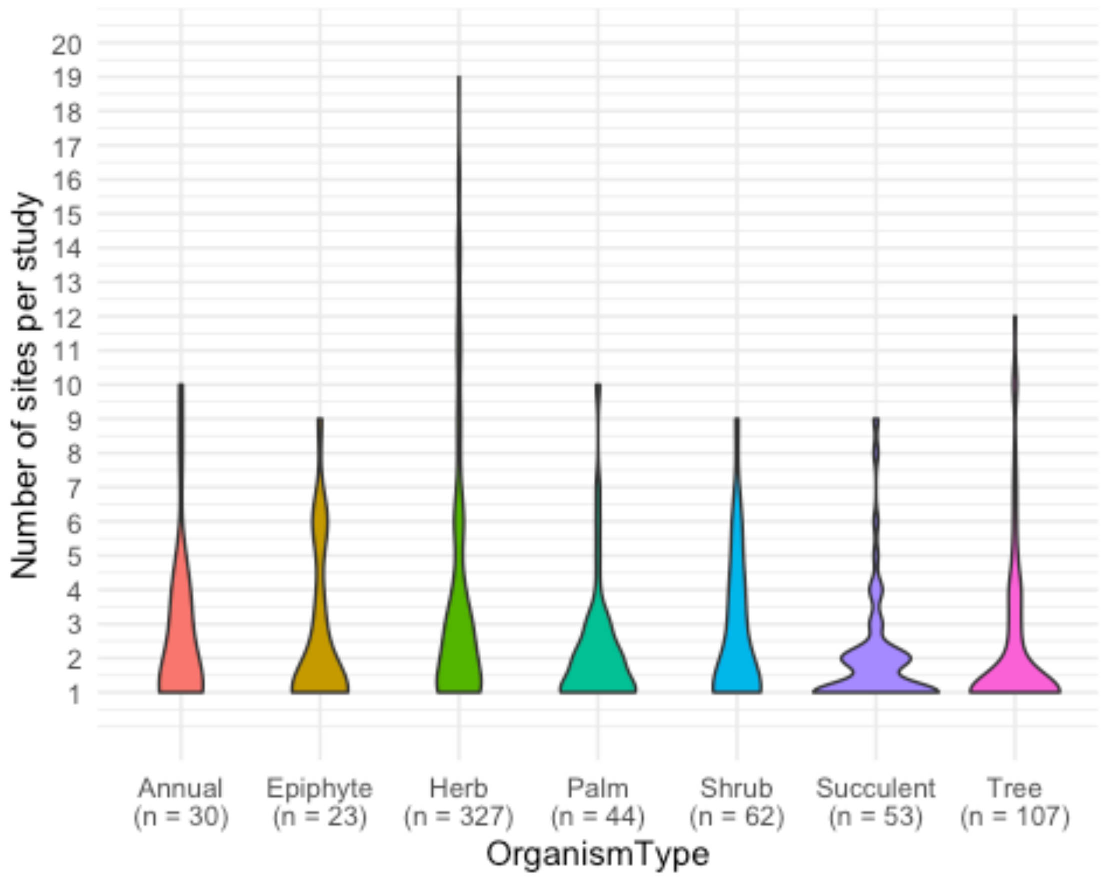


#### Matrix dimension

```
# average dimension per growth form
Dim_growth <- clean_db %>%
  select(OrganismType, Study, SpeciesAuthor, MatrixDimension) %>%
  unique()

model <- aov(MatrixDimension ~ OrganismType, data = Dim_growth)
out <- HSD.test(model, "OrganismType",
  group = TRUE, console = TRUE,
```

```
  main = "matDim across growth forms"
)
```

```
##
## Study: matDim across growth forms
##
## HSD Test for MatrixDimension
##
## Mean Square Error:  22.91226
##
## OrganismType,  means
##
##           MatrixDimension      std   r Min Max
## Annual           3.585366 2.301908  41   2  14
## Epiphyte         7.291667 8.467606  48   3  41
## Herb             5.309474 3.307550 475   1  29
## Palm             8.700000 6.578412  50   3  36
## Shrub            6.422222 3.280784  90   1  18
## Succulent        7.656250 2.918027  64   3  17
## Tree             9.483444 7.533794 151   2  60
##
## Alpha: 0.05 ; DF Error: 912
## Critical Value of Studentized Range: 4.178904
##
## Groups according to probability of means differences and alpha level( 0.05
)
##
## Treatments with the same letter are not significantly different.
##
##           MatrixDimension groups
## Tree             9.483444      a
## Palm             8.700000     ab
## Succulent        7.656250     ab
## Epiphyte         7.291667    abc
## Shrub            6.422222     bc
## Herb             5.309474     cd
## Annual           3.585366      d
```

```
plot(out)
```

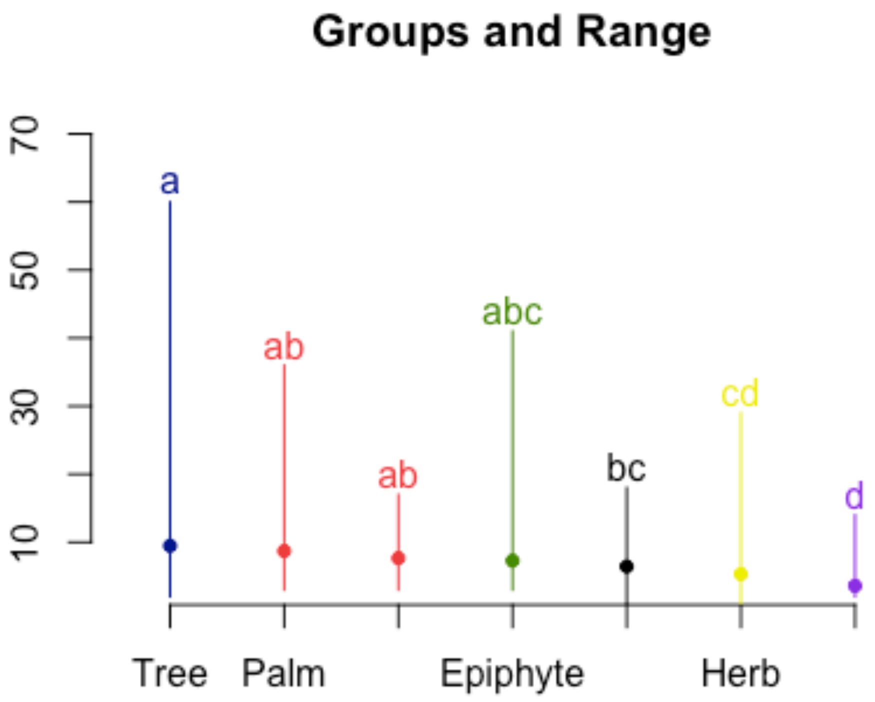


```r
Dim_growth <- clean_db %>%
  select(OrganismType, Study, MatrixDimension) %>%
  unique() %>%
  group_by(OrganismType)

# adding sample size to the x labels
sampleSize <- Dim_growth %>%
  group_by(OrganismType) %>%
  summarize(n = length(unique(Study)))

labelsGF <- paste0(sampleSize$OrganismType, "\n(n = ", sampleSize$n, ")")

# add comment that Bryophyte, Liana and Fern were removed due to small sample
size
textbox <- "Bryophyte, Liana and Fern were\nremoved due to small sample size"

# Cut off after NumberPopulation = 30 --> !!! this removes 2 data points in
Epiphytes, 1 in Palms and 1 in Trees
Plot_DimG <- ggplot(data = Dim_growth, aes(x = OrganismType, y =
MatrixDimension, fill = OrganismType)) +
  geom_violin() +
  scale_y_continuous(limits = c(0, 20), breaks = 1:20) +
  scale_x_discrete(labels = labelsGF) +
```

```
  theme_minimal() +
  theme(
    plot.title = element_text(hjust = 0.5),
    legend.position = "none",
    panel.grid.minor = element_blank()
  ) +
  ylab("Matrix dimension per study") +
  NULL

Plot_DimG
```

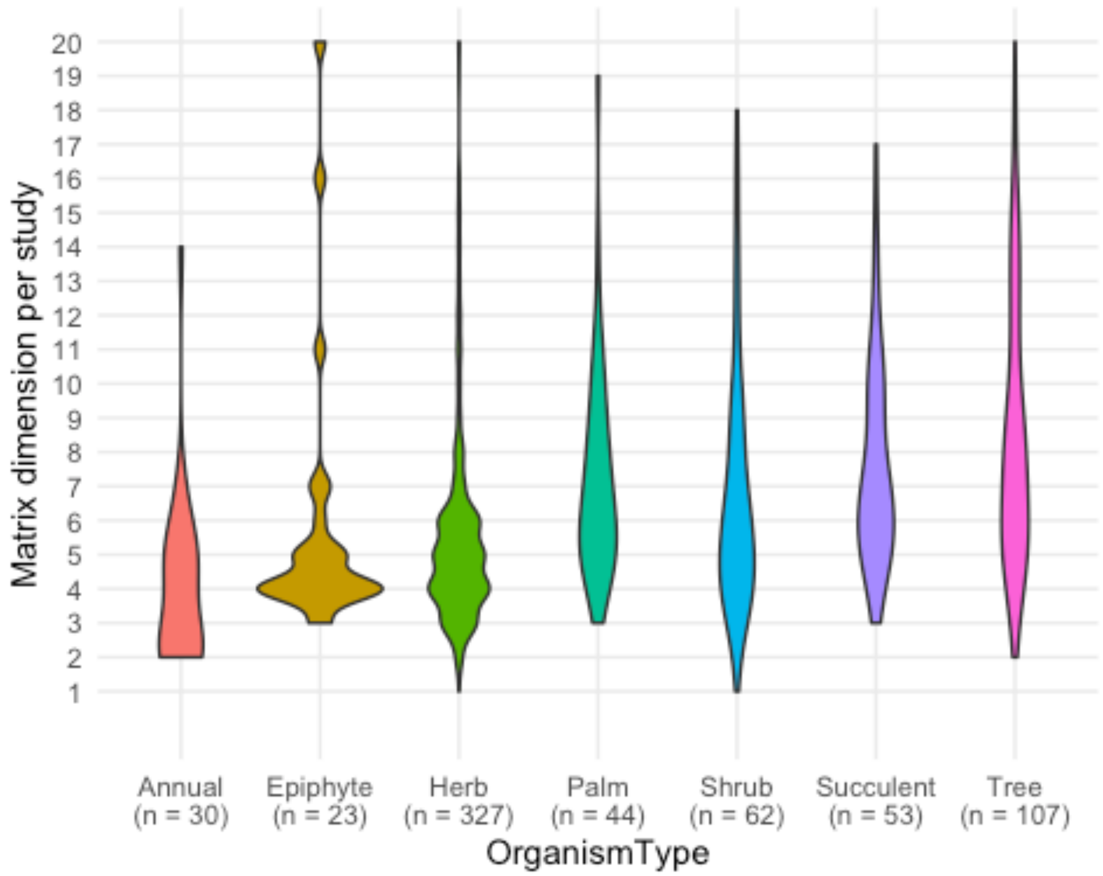


```
#Stats for this
Dim_growth <- Dim_growth %>%
  mutate(logDim = log(MatrixDimension))
mod1 <- lm(logDim~OrganismType,data = Dim_growth)
anova(mod1)

## Analysis of Variance Table
##
## Response: logDim
##               Df  Sum Sq Mean Sq F value    Pr(>F)
## OrganismType   6  32.967  5.4944  22.718 < 2.2e-16 ***
## Residuals    730 176.549  0.2418
```

```
## ---
## Signif. codes:  0 '***' 0.001 '**' 0.01 '*' 0.05 '.' 0.1 ' ' 1
```

#### Ecoregion

```
# get subset
eco <- clean_db_all %>%
  select(Study, SpeciesAuthor, Ecoregion, ecoregionLabel, MatrixDimension,
StudyDuration, NumberPopulations)
```

##### Temporal variation

```
Temp_eco <- eco %>%
  select(Ecoregion, ecoregionLabel, Study, SpeciesAuthor, StudyDuration) %>%
  unique()

model <- aov(StudyDuration ~ ecoregionLabel, data = Temp_eco)
out <- HSD.test(model, "ecoregionLabel",
  group = TRUE, console = TRUE,
  main = "study duration across ecoregions"
)

##
## Study: study duration across ecoregions
##
## HSD Test for StudyDuration
##
## Mean Square Error:  30.72688
##
## ecoregionLabel,  means
##
##                        StudyDuration       std   r Min Max
## Marine                      3.833333 0.9831921   6   3   5
## Mediterranean/desert        5.907895 5.2163209 152   1  51
## Temperate                   6.362319 6.7715493 414   1  51
## Tropical                    4.167401 3.0363861 227   1  30
## Tundra/boreal               5.038462 4.3474651  52   1  17
## Wetlands                    7.000000 4.2130749   9   3  16
##
## Alpha: 0.05 ; DF Error: 854
## Critical Value of Studentized Range: 4.039275
##
## Groups according to probability of means differences and alpha level( 0.05
)
##
## Treatments with the same letter are not significantly different.
##
##                        StudyDuration groups
## Wetlands                    7.000000      a
## Temperate                   6.362319      a
## Mediterranean/desert        5.907895      a
## Tundra/boreal               5.038462     ab
```

```
## Tropical                    4.167401     ab
## Marine                      3.833333     ab

plot(out)
```

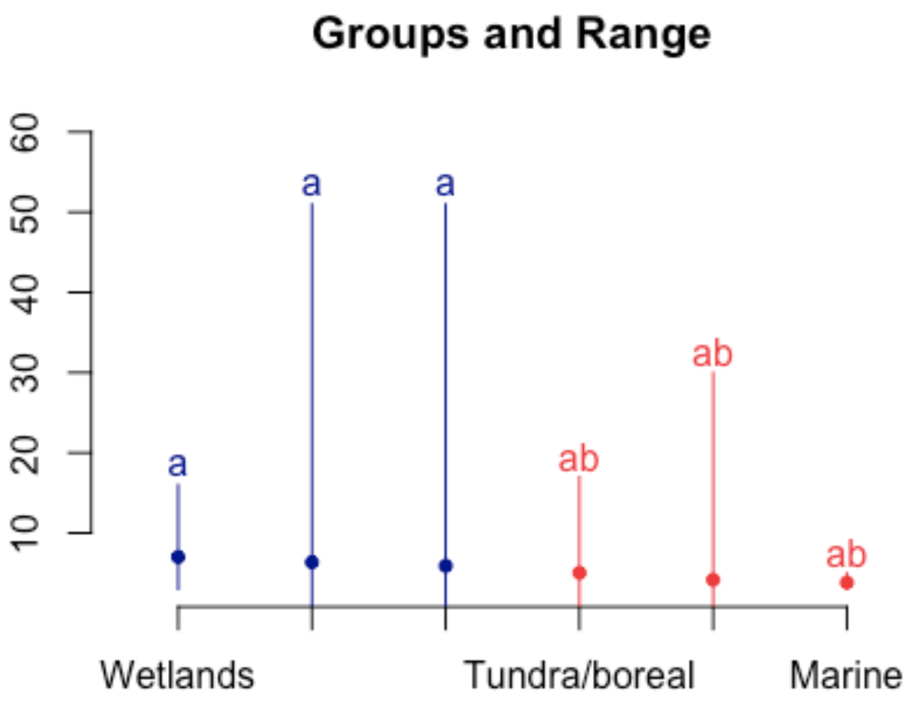


```
Temp_eco <- eco %>%
  select(SpeciesAuthor, ecoregionLabel, Ecoregion, Study, StudyDuration) %>%
  filter(!is.na(ecoregionLabel)) %>%
  unique()

# adding sample size to the x labels
sampleSize <- Temp_eco %>%
  group_by(ecoregionLabel) %>%
  summarize(n = length(unique(Study)))

labelsGF <- paste0(sampleSize$ecoregionLabel, "\n(n = ", sampleSize$n, ")")

# add comment that Bryophyte, Liana and Fern were removed due to small sample
size
textbox <- "Bryophyte, Liana and Fern were\nremoved due to small sample size"

# Cut off after NumberPopulation = 30 --> !!! this removes 2 data points in
Epiphytes, 1 in Palms and 1 in Trees
Plot_TempE <- ggplot(data = Temp_eco, aes(x = ecoregionLabel, y =
```

```
StudyDuration, fill = ecoregionLabel)) +
  geom_violin() +
  scale_y_continuous(limits = c(0, 20), breaks = 1:20) +
  scale_x_discrete(labels = labelsGF) +
  theme_minimal() +
  theme(
    plot.title = element_text(hjust = 0.5),
    legend.position = "none"
  ) +
  ggtitle("MatrixDimension per OrganismType") +
  xlab("Ecoregion") +
  ylab("Number of study years per study") +
  NULL

Plot_TempE
```

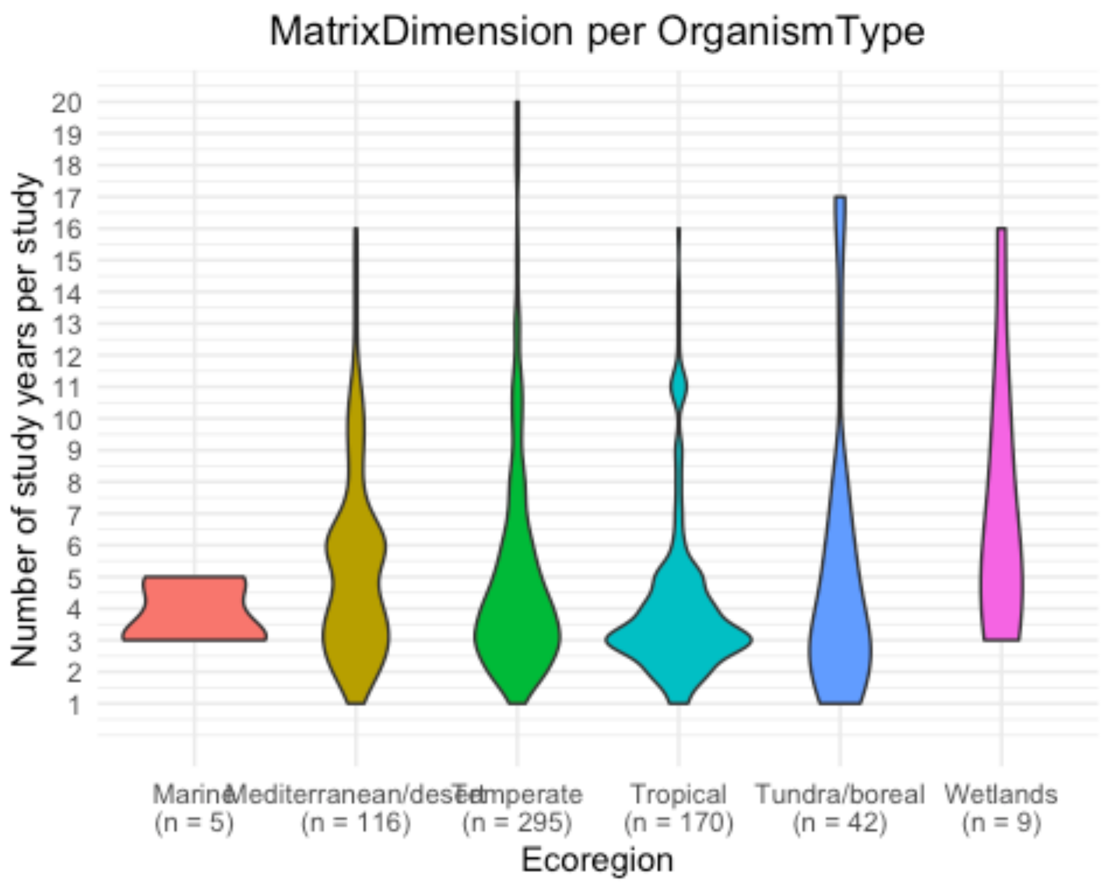


##### Spatial replication

```
Spatial_eco <- eco %>%
  select(Ecoregion, ecoregionLabel, Study, SpeciesAuthor, NumberPopulations)
%>%
  unique()

model <- aov(NumberPopulations ~ ecoregionLabel, data = Spatial_eco)
```

```r
out <- HSD.test(model, "ecoregionLabel",
  group = TRUE, console = TRUE,
  main = "spatial replication across ecoregions"
)

##
## Study: spatial replication across ecoregions
##
## HSD Test for NumberPopulations
##
## Mean Square Error:  9.764689
##
## ecoregionLabel,  means
##
##                        NumberPopulations       std   r Min Max
## Marine                          1.750000 0.9574271   4   1   3
## Mediterranean/desert            2.370130 2.3233717 154   1  12
## Temperate                       3.167076 3.5520891 407   1  36
## Tropical                        2.235808 2.6748228 229   1  21
## Tundra/boreal                   3.150943 2.9377924  53   1  14
## Wetlands                        4.600000 5.5617743  10   1  16
##
## Alpha: 0.05 ; DF Error: 851
## Critical Value of Studentized Range: 4.039307
##
## Groups according to probability of means differences and alpha level( 0.05
)
##
## Treatments with the same letter are not significantly different.
##
##                        NumberPopulations groups
## Wetlands                        4.600000      a
## Temperate                       3.167076      a
## Tundra/boreal                   3.150943     ab
## Mediterranean/desert            2.370130     ab
## Tropical                        2.235808     ab
## Marine                          1.750000     ab

plot(out)
```

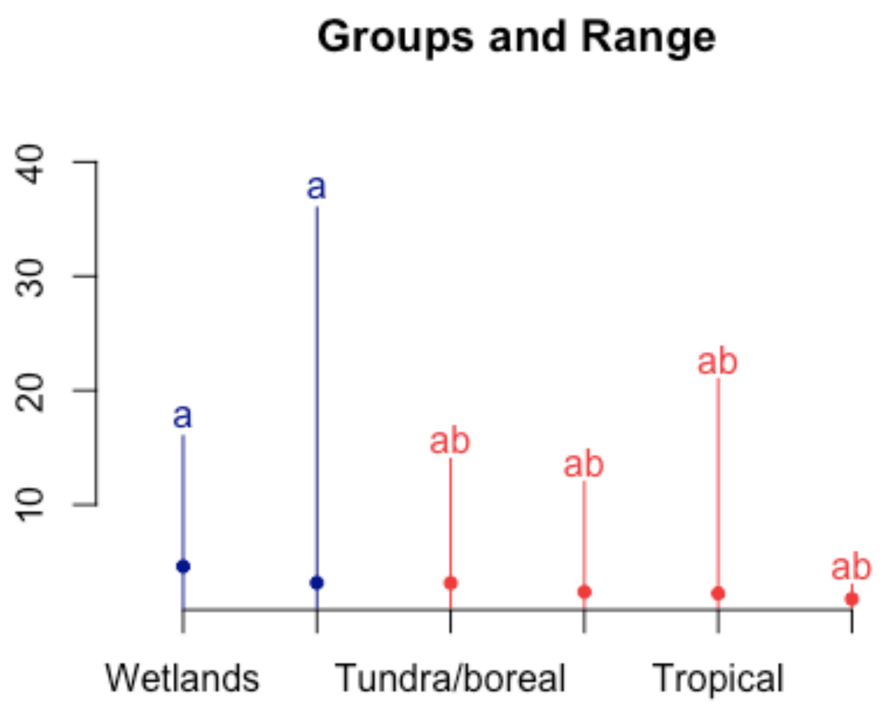


```
Spatial_eco <- eco %>%
  select(ecoregionLabel, Study, SpeciesAuthor, NumberPopulations) %>%
  filter(ecoregionLabel != is.na(ecoregionLabel)) %>%
  unique()

# adding sample size to the x labels
sampleSize <- Spatial_eco %>%
  group_by(ecoregionLabel) %>%
  summarize(n = length(unique(Study)))

labelsGF <- paste0(sampleSize$ecoregionLabel, "\n(n = ", sampleSize$n, ")")

# add comment that Bryophyte, Liana and Fern were removed due to small sample
size
textbox <- "Bryophyte, Liana and Fern were\nremoved due to small sample size"

# Cut off after NumberPopulation = 30 --> !!! this removes 2 data points in
Epiphytes, 1 in Palms and 1 in Trees
Plot_SpatialE <- ggplot(data = Spatial_eco, aes(x = ecoregionLabel, y =
NumberPopulations, fill = ecoregionLabel)) +
  geom_violin() +
  scale_y_continuous(limits = c(0, 20), breaks = 1:20) +
  scale_x_discrete(labels = labelsGF) +
```

```
  theme_minimal() +
  theme(
    plot.title = element_text(hjust = 0.5),
    legend.position = "none"
  ) +
  xlab("Ecoregion") +
  ylab("Number of sites per study") +
  NULL

Plot_SpatialE
```

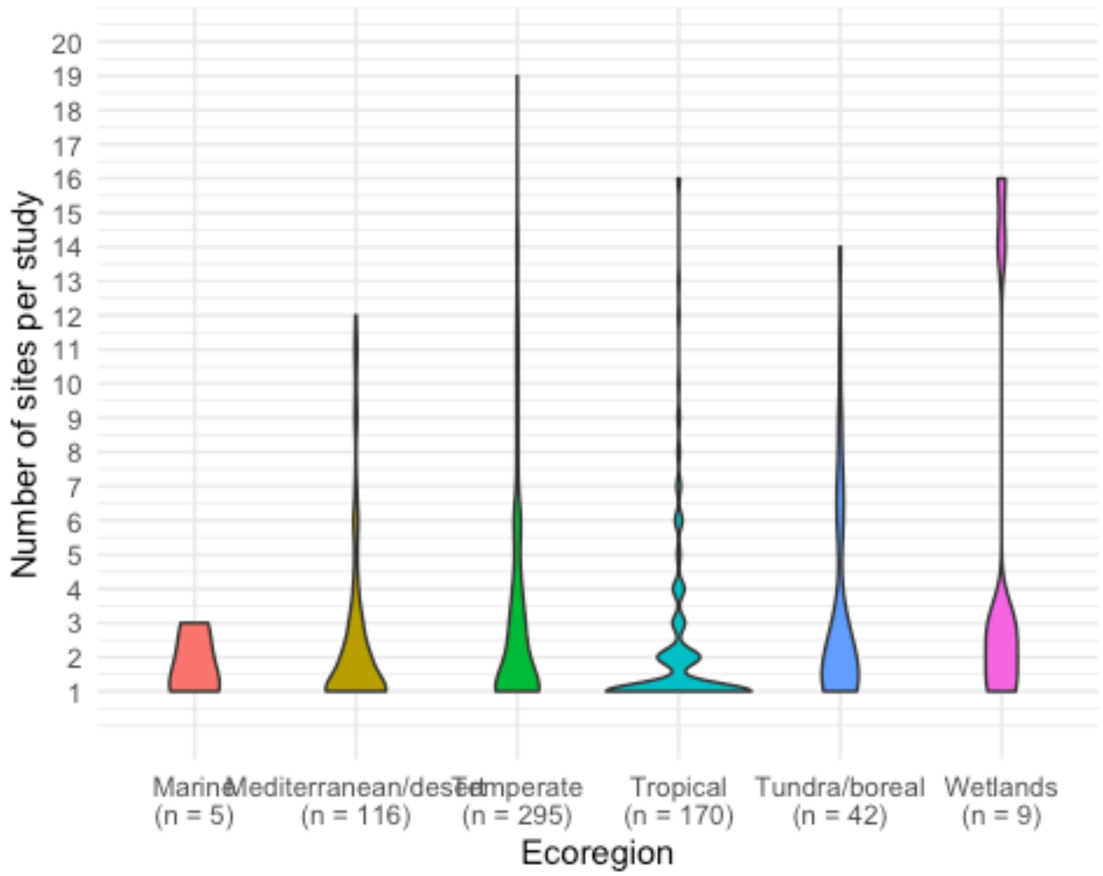


##### Matrix dimension

```
Dim_eco <- eco %>%
  select(Ecoregion, ecoregionLabel, Study, SpeciesAuthor, MatrixDimension)
%>%
  unique()

model <- aov(MatrixDimension ~ ecoregionLabel, data = Dim_eco)
out <- HSD.test(model, "ecoregionLabel",
  group = TRUE, console = TRUE,
  main = "matDim across ecoregions"
)
```

```
##
## Study: matDim across ecoregions
##
## HSD Test for MatrixDimension
##
## Mean Square Error:  24.63544
##
## ecoregionLabel,  means
##
##                        MatrixDimension      std   r Min Max
## Marine                        8.166667 3.125167   6   5  12
## Mediterranean/desert          5.727848 2.816351 158   2  20
## Temperate                     5.911833 5.212831 431   1  60
## Tropical                      8.510638 5.720231 235   2  41
## Tundra/boreal                 6.759259 4.596698  54   3  26
## Wetlands                      5.900000 3.665151  10   3  16
##
## Alpha: 0.05 ; DF Error: 888
## Critical Value of Studentized Range: 4.038923
##
## Groups according to probability of means differences and alpha level( 0.05
)
##
## Treatments with the same letter are not significantly different.
##
##                        MatrixDimension groups
## Tropical                      8.510638      a
## Marine                        8.166667     ab
## Tundra/boreal                 6.759259     ab
## Temperate                     5.911833      b
## Wetlands                      5.900000      b
## Mediterranean/desert          5.727848      b
```

```
plot(out)
```

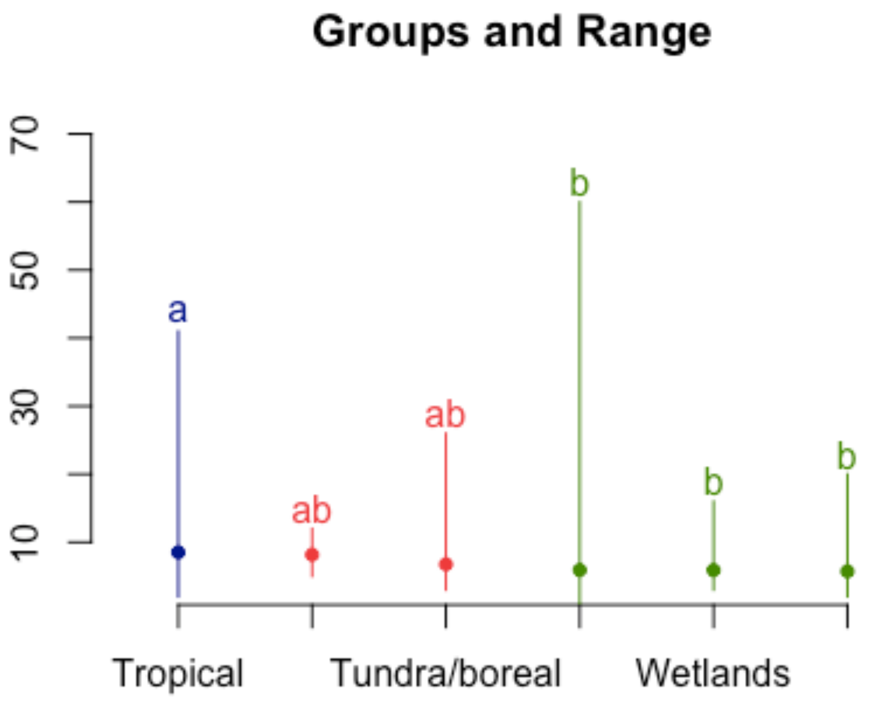


```
Dim_eco <- eco %>%
  select(ecoregionLabel, Study, SpeciesAuthor, MatrixDimension) %>%
  filter(!is.na(ecoregionLabel)) %>%
  unique()

# adding sample size to the x labels
sampleSize <- Dim_eco %>%
  group_by(ecoregionLabel) %>%
  summarize(n = length(unique(Study)))

labelsGF <- paste0(sampleSize$ecoregionLabel, "\n(n = ", sampleSize$n, ")")

# add comment that Bryophyte, Liana and Fern were removed due to small sample
size
textbox <- "Bryophyte, Liana and Fern were\nremoved due to small sample size"

# Cut off after NumberPopulation = 30 --> !!! this removes 2 data points in
Epiphytes, 1 in Palms and 1 in Trees
Plot_DimE <- ggplot(data = Dim_eco, aes(x = ecoregionLabel, y =
MatrixDimension, fill = ecoregionLabel)) +
  geom_violin() +
  scale_y_continuous(limits = c(0, 20), breaks = 1:20) +
  scale_x_discrete(labels = labelsGF) +
```

```
  theme_minimal() +
  theme(
    plot.title = element_text(hjust = 0.5),
    legend.position = "none",
    panel.grid.minor = element_blank()
  ) +
  xlab("Ecoregion") +
  ylab("Matrix dimension per study") +
  NULL

Plot_DimE
```

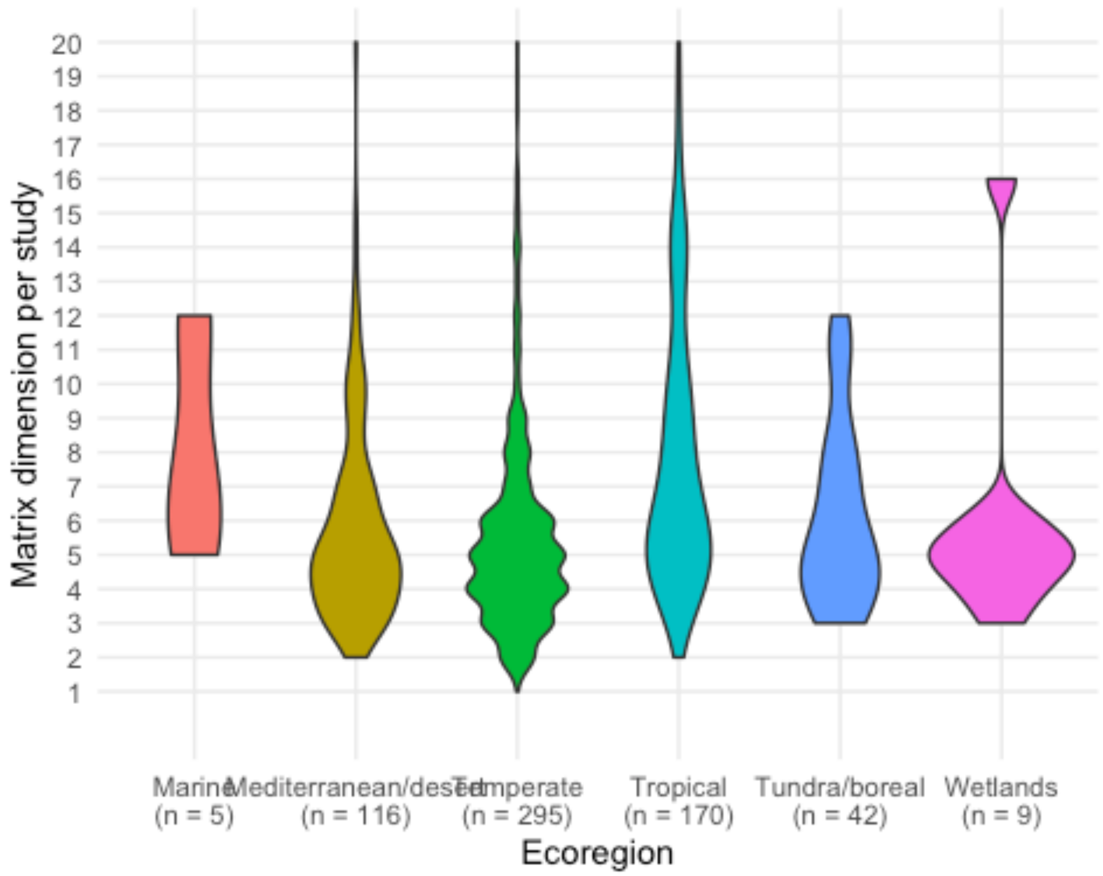


```
#Stats for this plot
Dim_eco<-Dim_eco %>%
  mutate(logDim = log(MatrixDimension))
names(Dim_eco)

## [1] "ecoregionLabel"  "Study"           "SpeciesAuthor"
"MatrixDimension"
## [5] "logDim"

mod1 <- lm(logDim~ecoregionLabel,data = Dim_eco)
summary(mod1)
```

```
##
## Call:
## lm(formula = logDim ~ ecoregionLabel, data = Dim_eco)
##
## Residuals:
##      Min       1Q   Median       3Q      Max
## -1.60706 -0.37273 -0.03624  0.29963  2.48729
##
## Coefficients:
##                                   Estimate Std. Error t value Pr(>|t|)
## (Intercept)                         2.0404     0.2077   9.825   <2e-16
***
## ecoregionLabelMediterranean/desert  -0.3947     0.2116  -1.865   0.0625 .
## ecoregionLabelTemperate             -0.4333     0.2091  -2.072   0.0386 *
## ecoregionLabelTropical              -0.0582     0.2103  -0.277   0.7821
## ecoregionLabelTundra/boreal         -0.2587     0.2193  -1.180   0.2385
## ecoregionLabelWetlands              -0.3739     0.2627  -1.423   0.1550
## ---
## Signif. codes:  0 '***' 0.001 '**' 0.01 '*' 0.05 '.' 0.1 ' ' 1
##
## Residual standard error: 0.5087 on 884 degrees of freedom
## Multiple R-squared:  0.09225,    Adjusted R-squared:  0.08711
## F-statistic: 17.97 on 5 and 884 DF,  p-value: < 2.2e-16
```

```
anova(mod1)
```

```
## Analysis of Variance Table
##
## Response: logDim
##                 Df  Sum Sq Mean Sq F value    Pr(>F)
## ecoregionLabel   5  23.248  4.6496  17.967 < 2.2e-16 ***
## Residuals      884 228.769  0.2588
## ---
## Signif. codes:  0 '***' 0.001 '**' 0.01 '*' 0.05 '.' 0.1 ' ' 1
```

```
t.test(x = Dim_eco$logDim[Dim_eco$ecoregionLabel == "Tropical"],y =
Dim_eco$logDim[Dim_eco$ecoregionLabel == "Mediterranean/desert"])
```

```
##
##  Welch Two Sample t-test
##
## data:  Dim_eco$logDim[Dim_eco$ecoregionLabel == "Tropical"] and
Dim_eco$logDim[Dim_eco$ecoregionLabel == "Mediterranean/desert"]
## t = 6.8561, df = 375.36, p-value = 2.92e-11
## alternative hypothesis: true difference in means is not equal to 0
## 95 percent confidence interval:
##  0.2399879 0.4329962
## sample estimates:
## mean of x mean of y
##  1.982169  1.645677
```
